# Local adaptations of Mediterranean sheep and goats through an integrative approach

**DOI:** 10.1101/2021.01.22.427461

**Authors:** Bruno Serranito, Marco Cavalazzi, Pablo Vidal, Dominique Taurisson-Mouret, Elena Ciani, Marie Bal, Eric Rouvellac, Bertrand Servin, Carole Moreno-Romieux, Gwenola Tosser-Klopp, Stephen J. G. Hall, Johannes A. Lenstra, François Pompanon, Badr Benjelloun, Anne Da Silva

## Abstract

In a context of climate change, identifying the genes underlying adaptations to extreme environments is essential. Small ruminants are adapted to a wide variety of habitats and thus, are promising study models. Here, we considered 17 goat and 25 sheep local Mediterranean breeds, in Italy, France and Spain. We proposed, and empirically tested a new methodology to highlight selection signatures in relation to the environments. Based on historical archives, we selected the breeds potentially most linked to a territory and defined their original geographical cradle, including transhumant pastoral areas. For both species, the cradles were arranged along latitudinal gradient of aridity and altitude. Then we used the programs PCAdapt and LFMM. Considering cradles, instead of current GPS coordinates, markedly improved the sensitivity of the LFMM analyses. All the results combined with a systematic literature review, revealed a set of genes with potentially key adaptive roles. Some of these genes, have been found implicated in lipid metabolism (SUCLG2, BMP2), hypoxia/heat stress (UBE2R2/UBAP2), lung function (BMPR2), seasonal patterns (SOX2, DPH6) or neuronal function (TRPC4, TRPC6). For the intergenic region between PCDH9 and KLH1, as well as NBEA, identified in both species, the adaptive role could be strong. We found for RXFP2, associated with sheep horn development, and MSRB3 and SLC26A4, both associated with hearing of sheep, a strong association with the environment gradient. We conclude that our new methodology, which considers breeds in their historical, environmental and anthropological context, provides novel and essential information on local breeds and their unique adaptations.

## INTRODUCTION

Sheep and goats were among the first mammalian livestock species to be domesticated, in a process that started at ca. 10,000-9,500 B.P. in Mesopotamia (Vigne, 2015). The subsequent expansion across the Mediterranean Basin via maritime transport included different waves of colonization. The remains testifying to these founding periods along the northern coasts of the western Mediterranean are dated from 8,100-7,700 B.P. (Zeder, 2008).

Waves of dispersal, which have dispatched small groups of individuals (founder effect) to very diverse habitats, are the starting point of the process that led to the emergence of breeds (Sponenberg & Bixby, 2007). Traditional pastoralism, characterized by limited human intervention (i.e., “soft” artificial selection, Taberlet et al., 2008), and in counterpart by a strong influence of natural selection, led over the millennia, to populations genetically differentiated and adapted to various agro-climatic conditions (Berihulay et al., 2019). They provide an interesting model for deepening our knowledge of adaptive mechanisms under extreme environmental conditions. In industrialized countries, artificial breeding has intensified over the past 200 years (Taberlet et al., 2008), focusing mainly on agronomic traits related to milk, meat production and fiber quality. Very many breeds have by crossbreeding and/or intensive artificial selection, largely lost the link with their original territory. In contrast, the so-called local breeds have remained at least partially, connected to their original environments. We suggest it is among these breeds that selection signatures related to environmental adaptations will be most detectable. For example, studies on local sheep in Ethiopia (Edea et al., 2019) and Tibet (Yang et al.; 2016, Wei et al.; 2016) have identified several genes involved in adaptation to thermal stress and hypoxia. Kim et al. (2016) considering both local goats and sheep of hot and arid environment, have highlighted genes implicated in the adaptation to thermal stress.

This study aimed at identifying the genomic bases of adaptation, in local breeds of goats and sheep, distributed along the Mediterranean arc, in Italy, the south of France and Spain. The area under consideration harbours contrasting environments, such as high altitude areas (Alps, Pyrenees, Corsica), coastal areas, wetlands and arid areas, with closely related sheep and goat breeds (Ciani et al., 2020; Colli et al., 2018; Kijas et al., 2009). All these features provide an ideal situation for the identification of adaptation signatures.

We hypothesize that selection signatures relevant to environmental adaptation, will be more reliably identified by reference to the environmental conditions of the breed’s cradle of origin, rather than to those of the locations where the animals were actually sampled. Indeed, breeds of domestic animals are at the very interface between anthropized and natural environments, which implies specificities with regard to wild animals. First, (i) sheep and goat breeds have originally emerged within socially structured human groups, sometimes referred to as clans or tribes, occupying a well-defined territory, and within which herds were transmitted from generation to generation (Brisebarre, 2009; Hall, 2019). In industrialized countries, the traditional links between territory, social group and breeds has become distorted. Thus, herds of a given breed can be found outside of its original cradle. Hence, the search for a selection signature may be optimized by considering the environmental conditions of the breed’s cradle, where the genomes have been shaped over time, rather than the current location of herds. Moreover, (ii) we should consider that traditional pastoralism, was largely based on transhumance. During these periods, the animals are outdoor, most often in mountain altitude summer pastures. While human intervention is minimal, the strength of natural selection is particularly pronounced. Even if transhumance practices are increasingly abandoned (Caballero et al., 2011; Collantes, 2009), they have fundamentally shaped the genetic adaptations of sheep and goats. It is clearly important that this process be methodologically investigated.

In this study, our objective is to test empirically our method on datasets that have been previously investigated, in order to assess if the use of historical and anthropological dimensions of breeds improves the detection of selection signatures. We reasoned that (i) selection of authentic local breeds, to be included in the analysis, will limit the background noise caused by breeds for which the link to the environment is weak or has eroded, for instance by crossbreeding or migration; and (ii) targeting the geographical area in which populations have evolved rather than the present distribution will optimize the identification of adaptive genomic regions. Next, the proposed methodology was implemented to search for selection signatures underlying local adaptation. We attached particular importance to the selection signatures involving homologous genomic regions in sheep and goats.

## MATERIAL AND METHODS

### 1) Methodology

A summary diagram of the main steps of the proposed approach is shown in Supplementary Figure 1.

#### a) Selection of breeds

The aim was to keep breeds with the highest probability of having retained strong adaptations to the local environment, i.e. breeds least affected by the disruptions inherent to the intensification of agricultural practices. For the study, we have targeted the Mediterranean arc in France, Italy and Spain. This region has the advantage of harboring closely related sheep and goat breeds, in spite of large environmental contrasts (Ciani et al., 2020; Colli et al., 2018; Kijas et al., 2009). We initially, considered 32 breeds of goats and 51 breeds of sheep (Supplementary Table 1). Subsequently, we selected 17 goat breeds and 25 sheep breeds (Supplementary Figure 2), on the basis of historical and anthropological documentation and according to the following criteria: (i) the breeds are local (*i.e*. cosmopolitan breeds were excluded); (ii) the breeds have featured in the local tradition for at least one century; (iii) any recorded introgression of exotic breeds was limited and occurred more than 100 years ago. If available, recent evaluations of admixture were also used to select breeds to include (see Ciani et al. 2014; Manunza et al., 2016; Rochus et al., 2018; Bertolini et al., 2018; Oget, Servin & Palhière, 2019). Location and outline descriptions of the selected breeds are in Supplementary Figure 2 and Supplementary Table 2.

#### b) Geographic definition of cradle of origin

We determined cradles of origin on the basis of historical and anthropological documentation. Particular attention has been paid to transhumance practices (see Supplementary Figure 2 and Supplementary Table 2 for details). We used ArcGIS (ESRI, 2011) to map the cradles, in such a way to include areas of summer pastures when so-called vertical transhumance was recorded. The horizontal transhumance, involving movements covering long distances without significant altitude differences (Ruiz & Ruiz, 1986) cannot be directly taken into account because a large horizontal range would complicate the definition of the cradles.

#### c) Environmental characterization

Bioclimatic data were obtained from the WorldClim database (v.1.4, Hijmans et al., 2005) covering the period from 1960 to 1990, with a spatial resolution of 30 arc-seconds in the WGS84 datum. We considered the bioclimatic variables most relevant to highlight the major contrasts in temperature and humidity between the cradles, i.e.: BIO1=Annual Mean Temperature, BIO5=Maximum Temperature of Warmest Month, BIO_6_=Minimum Temperature of Coldest Month, BIO_12_=Annual Precipitation, BIO_13_=Precipitation of Wettest Month, BIO_14_= Precipitation of Driest Month, BIO_15_=Precipitation Seasonality (Coefficient of Variation). Altitude information was collected from the SRTM 90 m Digital Elevation Database (v.4.1) (Jarvis, Reuter, Nelson & Guevara, 2008). Finally, latitude was taken as a proxy for luminosity and seasonality. This procedure was performed in R 3.5.2 (R Core Team, 2018) using the R package RSAGA (Brenning, 2008).

Because our objective was not to estimate allele-environment correlations between each SNP and each environmental variable at a time, but to capture the relation between the genome and the environment considered as a whole, the cradles were clustered according to their climatic proximity, following this procedure:

For each cradle, the environmental variables of 10,000 random location points were extracted, using R packages sp (Pebesma & Bivand, 2005; Bivand, Pebesma, & Gomez-Rubio, 2013) and rgdal (Keitt, Bivand, Pebesma, & Rowlingson, 2010).

In order to identify groups of breeds with relatively close environmental conditions in their cradle area, principal component analysis (PCA) was performed on the environmental variables. Prior to the analysis, Spearman correlation coefficient identified highly correlated variables (*i.e*. correlation coefficient r□0.9), in which case the biologically less relevant variable was removed to facilitate graphical presentation. The PCA analysis was followed by a hierarchical clustering on principal components (HCPC), using Euclidian distances and Ward’s method. Multivariate analyses were performed with R software, using the FactoMineR package (Le, Josse, & Husson, 2008).

The LFMM (Frichot et al., 2013) analyses were conducted following the classification of the cradles obtained through the PCA/HCPC procedure.

#### d) Genotyping

For goats, we used the AdaptMap dataset, including worldwide breeds genotyped with the Caprine SNP50 BeadChip (available via Dryad: https://doi.org/10.5061/dryad.v8g21pt), which contains genotypes for 53,547 SNPs. For sheep, we merged the French dataset obtained with the Illumina Ovine HD SNP Beadchip (Zenodo repository 10.5281/zenodo.237116), with the Italian dataset (Ciani et al., 2014, provided by the authors) and the Spanish dataset (Manunza et al., 2016, provided by the authors) obtained with the Illumina Ovine SNP50 BeadChip. SNP data from French breeds were extracted from the 600K variation using the Ovine SNP50 BeadChip coordinates of SNPs on the OAR v3.1 reference genome assembly using Vcftools (Danecek et al., 2011). Merging with the SNP50 data resulted in 40,455 genotypes. SNPs and animals were pruned with PLINK v1.07 (Purcell et al., 2007) using the following filtering thresholds: (i) SNP call rate ≤97%; (ii) SNP minor allele frequency (MAF) ≤1%; (iii) animals displaying ≥10% of missing genotypes. After filtration of the merged datasets we retained 50,329 genotypes for 416 goats and 32,168 genotypes for 555 sheep (initial datasets were respectively of 420 and 576 individuals).

#### e) Identification and analysis of genomic selection signatures

We identified loci that may support local adaptation by two methods: (i) the individual-based multivariate approach PCAdapt (Luu et al., 2017), via the pcadapt R package and (ii) the latent factor mixed models LFMM, as implemented in the R package LEA (Frichot & François, 2015) for testing genotype-environment association.

PCAdapt scans the genome to detect outliers with respect to the population structure. Unlike population-based approaches, the package deals with admixed individuals and does not assume prior knowledge of population structure. As recommended by Luu et al. (2017), the optimal number of principal components (*i.e*., K, the optimal number of genetic groups) was determined using the graphical PCAdapt function, by varying K from 1 to 35, and following “Cattell’s rule”, i.e., keeping PCs that correspond to eigenvalues to the left of the lower straight line in the screeplot (Cattell, 1966). This led us to choose K=15 for goats and K=20 for sheep (see Supplementary Figure 8). Candidate SNPs were identified by calculating the False Discovery Rate (FDR, α□=□0.05) of the p-values associated with Mahalanobis distance estimated by PCAdapt, using the R package qvalue (Storey & Tibshirani, 2003).

LFMM software assesses associations between genetic variation (response variable) and environmental factors (explaining variables) using a linear mixed model and controlling for neutral genetic structure, such as population history and isolation-by-distance, with (random) latent factors, *via* an MCMC algorithm. The number of latent factors was chosen in accordance with the Admixture (Lee & Seung, 1999) analyses. We ran LFMM using 1000 sweeps for burn-in and 9000 additional sweeps. Since LFMM uses a stochastic algorithm, ten runs with different seeds were performed. We then chose significant associations based on FDR (α□=□0.05) using the R package qvalue. LFMM was used to test for the correlation between the genetic variations and the environment taken as a whole (*i.e*. the variables were not considered one by one, except to test the method), but the analysis was performed once using the PCA/HCPC cradle clustering results (see details in section 1.c).

For both PCAdapt and LFMM, we applied a stringent screen to identify selection signatures: candidate-selected regions were required to have at least 3 SNPs ≤ 500 kb apart, previously identified based on FDR and showing p-value ≤ 10^-9^. The window was chosen on the basis of previous evidence that LD in sheep and goats does not exceed 500 kb (see Supplementary Figure 3). It should be noted that the size of the window considered is widely used and that the selection criteria retained are among the most stringent (see Chen et al. 2016 and 2018; Cheruiyot et al., 2018; Lopez et al., 2019; Avila et al., 2018; Ablondi et al., 2019). For each analysis, genes within a region spanning 100 kb upstream and downstream of the candidate selection regions were annotated.

The chromosomal regions under selection pressure were inspected using NCBI Genome Data Viewer (https://www.ncbi.nlm.nih.gov/genome/gdv/), Oar_v3.1 for sheep and ARS1 for goats. Biological functions of the genes identified were inferred from the literature. Studies focusing on environmental adaptation signatures involving the same genes as those identified here, were compiled. As top candidate genes for adaptation, we took genes identified by our study and by at least one other study focusing on environmental adaptation. These genes, were explored for biological processes through Gene Ontology (GO) enrichment analyses, using GOrilla algorithm (Eden et al., 2009). The 19,932 goat genes associated with a GO term were used as background reference for the goat analysis; for the sheep analysis, 22,680 sheep genes were used as background reference. The criterion for selection of significantly enriched GO terms was q-value < 0.05 (Benjamini & Hochberg, 1995).

### 2) Experimental test of the method

We compared the cradle-based method with the use of the individual GPS coordinates. This comparison was carried out entirely for the goat dataset, for which the AdaptMap Consortium provided us with the geographical locations for all breeds considered. However, for sheep, geographical coordinates were only available for the French breeds.

For each GPS coordinate, the environmental variables were extracted as described in section 1.c, according to a circling area with a radius of 5 km centered on the GPS point. We considered the most discriminating variables: Annual Mean Temperature, Annual Mean Precipitation and Altitude, in order to compare the average information obtained by breed, from the GPS coordinates with that obtained from the cradles. Multiple comparisons with FDR adjusted p-values (α=0.05) were then realized with R, using the package rstatix.

The LFMM analysis proceeded as follows (see Figure 1):

- The individual GPS method: we assessed the LFMM correlation between individual genotypes and the variables, Annual Mean Temperature, Annual Mean Precipitation and Altitude, considered separately and extracted for each individual GPS coordinate.
- The GPS area method: we considered environmental data for the set of individual GPS coordinates recorded for the breed. (i) The LFMM correlation was calculated between each genotype and the average value obtained for the different variables considered separately (Annual Mean Temperature, Annual Mean Precipitation and Altitude). (ii) As alternative approach, we used all the environmental variables (see section 1.c) recorded at the set of the breed GPS coordinates, in order to group the different GPS areas according to their similarity, following the PCA/HCPC process as explained in section 1.c. Then we tested LFMM correlation between the genotypes and the resulting cluster ranking.
- The novel cradle method: finally, we used the method we propose. For each breed, we considered environmental data at the cradle level. The LFMM correlation was tested in two steps, as for the GPS area method.

**Figure 1.**
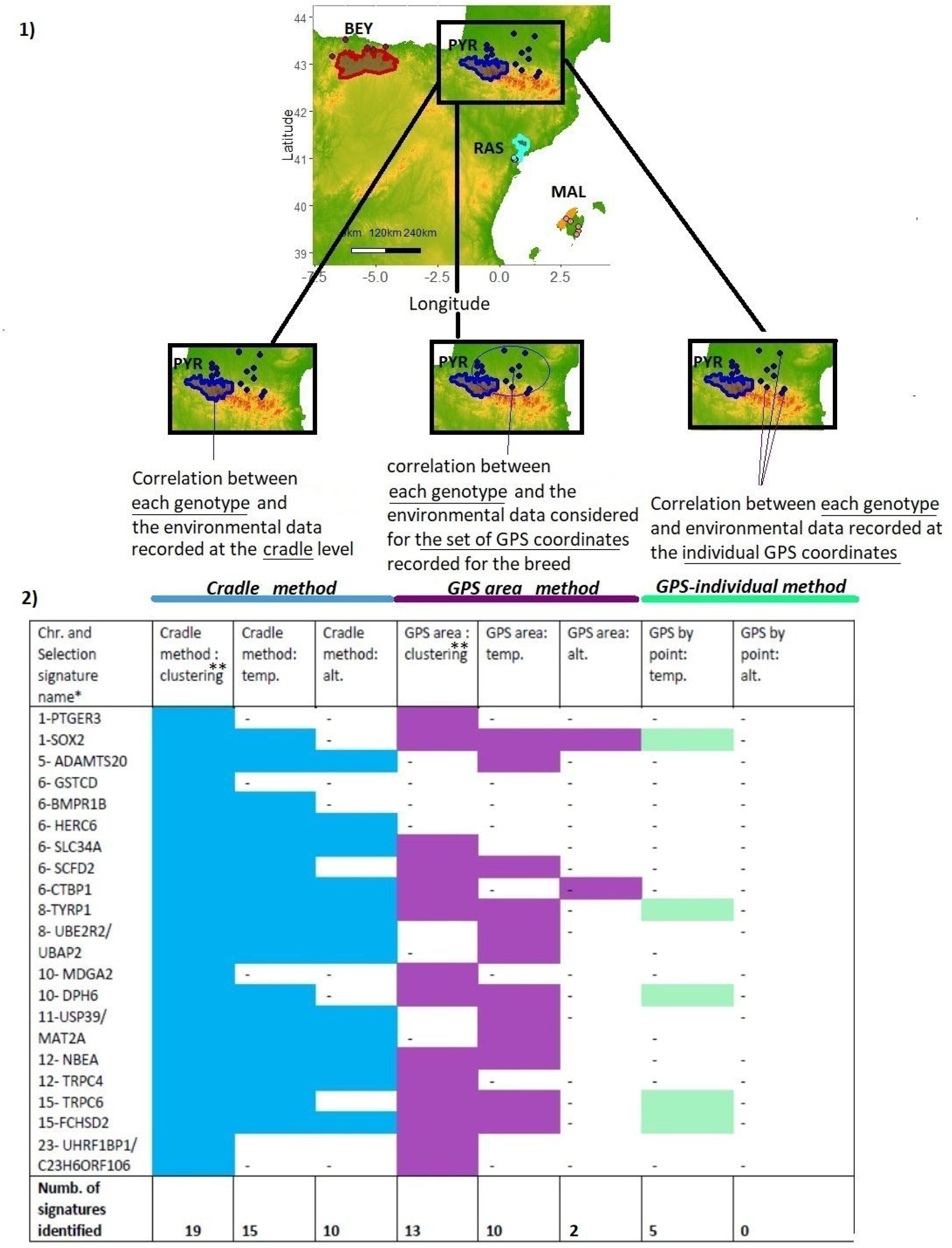
Experimental test of the method. The signature search was done via LFMM. The environment was characterized either at the level of the GPS coordinates (individual GPS method), or considering all the GPS points recorded for a given breed (GPS area method), or at the level of the cradle (cradle method). The environmental variables Altitude and Mean Annual Temperature were considered individually. Chr.: chromosome number. * Selection signature are named according to the gene(s) most likely targeted by the selection process, taking into account the distance of the SNPs from the genes, the associated p-values and the number of SNPs near or in the gene(s); it is possible to know in detail the genes associated with each selection signature (see supplementary table 3). ** Clustering: the environment was characterized considering all the variables at the same time and according to the PCA/HCPC process.

### 3) Complementary analyses

#### a) LD decay

To provide an insight into the overall levels of LD in the different breeds, genome-wide pairwise r^2^ values of SNPs separated by a maximum distance of 10 Mb, were calculated with PLINK software.

#### b) Admixture analyses

To assess population genetic structure we used ADMIXTURE software (Alexander, Novembre & Lange, 2009) Prior to these analyses, SNP pruning was used to select a subset of SNPs with minimal linkage disequilibrium via the --indep option of PLINK with the following parameters: 50 SNPs per window, a shift of five SNPs between windows, and a Variation Inflation Factor’s threshold of two (corresponding to r^2^>0.5). SNPs identified by the study as potentially selected were removed. For K=2 to K=25, ten independent runs were performed. The entropy criterion was calculated via the sNMF function implemented in the R package LEA to assess the number of ancestral populations that best explains the genotypic data (Alexander and Lange, 2011; Frichot et al., 2014). We used the program CLUMPAK (Kopelman et al., 2015, http://clumpak.tau.ac.il) to analyze the multiple independent runs at a single K and visualize the results.

#### c) Mantel test

Finally, we assessed the correlation between genetic distances and geographic distances using the Mantel test via the mantel. randtest function, implemented in the R adegenet package (Jombart, 2008).

## RESULTS

### 1) Breed analyses

For 15 of the 42 selected breeds we found documented evidence dating back several millennia (Supplementary Table 2). The remaining breeds have a history of at least several centuries, with the exception of 4 Italian goat breeds (DIT, CCG, GAR, ARG) and 3 Italian sheep breeds (DEL, BIE, VAL), that have an origin of about one century. Seventeen breeds were classified as “critical” or “endangered”. 30 breeds have been described as showing adaptations to stressful environments and 29 are, or were, transhumant. For most transhumant breeds the transhumance was, or is, vertical, so we included their summer mountain pasture in the geographical definition of the cradle (see Supplementary Figure 2 and Supplementary Table 2).

The Admixture analysis is detailed in Supplementary text 1. The entropy curves obtained, as well as the Mantel tests postulated distance isolation patterns. Therefore, we used K=1 and K=2 to correct for the limited genetic structure in the LFMM analyses (Dalongeville et al., 2018; De Kort et al., 2015; Capblancq et al., 2018).

### 2) Experimental testing of the method

#### a) Environmental characterization

For eight French sheep breeds we could compare Mean Annual Precipitation, Mean Annual Temperature and Altitude obtained via the GPS area method and the cradle method, respectively (Supplementary Figure 4), generating 21 comparisons. All differences between the GPS area and the cradle were significant except for the variable, Mean Annual Precipitation, for the breeds Causse du Lot (CDL) and Préalpes du Sud (PAS). Overall, significantly higher mean Altitude values were provided by the cradle method, which resulted in higher Mean Annual Precipitation and lower Mean Annual Temperatures, compared to that obtained by the GPS area method. The differences were particularly strong for transhumant breeds such as Rava (RAV), Tarasconnaise (TAR) and Mourerous (MOUR), for which GPS sampling had mainly been carried out on the plains.

For goats, we compared the same three environmental parameters obtained by the GPS area and the cradle methods for all 17 breeds, (Supplementary Figure 4), which resulted in 45 out of 51 significant differences. As found for sheep, significantly higher mean Altitude was always found for the cradle method, and resulted in lower Mean Annual Temperature. The rainfall pattern was less clear-cut, with 56% of the comparisons showing higher Mean Annual Precipitation when extracted at the level of cradle of origin.

These analyses showed that the environmental characterization depends on the definition of the distribution range based on the cradle or on the individual/breed GPS coordinates. The cradle method takes into account summer grazing areas, which implies the extraction of climatic data at higher altitudes than with the GPS method, generally resulting in lower temperatures and higher precipitations. In addition, for some breeds such as Garganica (GAR) and Mourerous (MOUR), individuals were sampled more than 100 km from their established cradle (Supplementary Figure 4), which led to the consideration of different climatic environments depending on the method used.

#### b) Selection signatures in relation to environment

In order to assess whether these differences in the environmental characterization affect the identification of the selection signature, we performed the LFMM analyses on the goat dataset for the different methods (see Material and Methods and Figure 1). The poor results obtained with the Annual Mean Precipitation are not shown. Notably, for the Altitude, individual GPS method did not detect any signature; the GPS area method identified only two signatures, whereas 10 signatures were recorded with the cradle method. For the Mean Annual Temperature, 5 signatures were observed with the individual GPS method, 10 with the GPS area method, and 15 by the cradle method. The process of clustering the environments allowed the detection of 13 signatures via the GPS areas and 19 (maximum value) via the cradles (see details in Supplementary Figures 5 and 6).

### 3) Goat dataset: track for selection signatures

From the 37 signatures, 21 were identified by PCAdap, 19 by LFMM; 3 signatures were supported by both methods (Table 1; Supplementary Table 5 and 6). Seven signatures were observed on chromosome 6 only, and four on chromosome 12. These signatures were documented in the literature (Supplementary Table 3). Of the 21 signatures highlighted by PCAdapt, 15 were found to be involved in adaptive processes and 15 by LFMM out of 19.

**Table 1.**
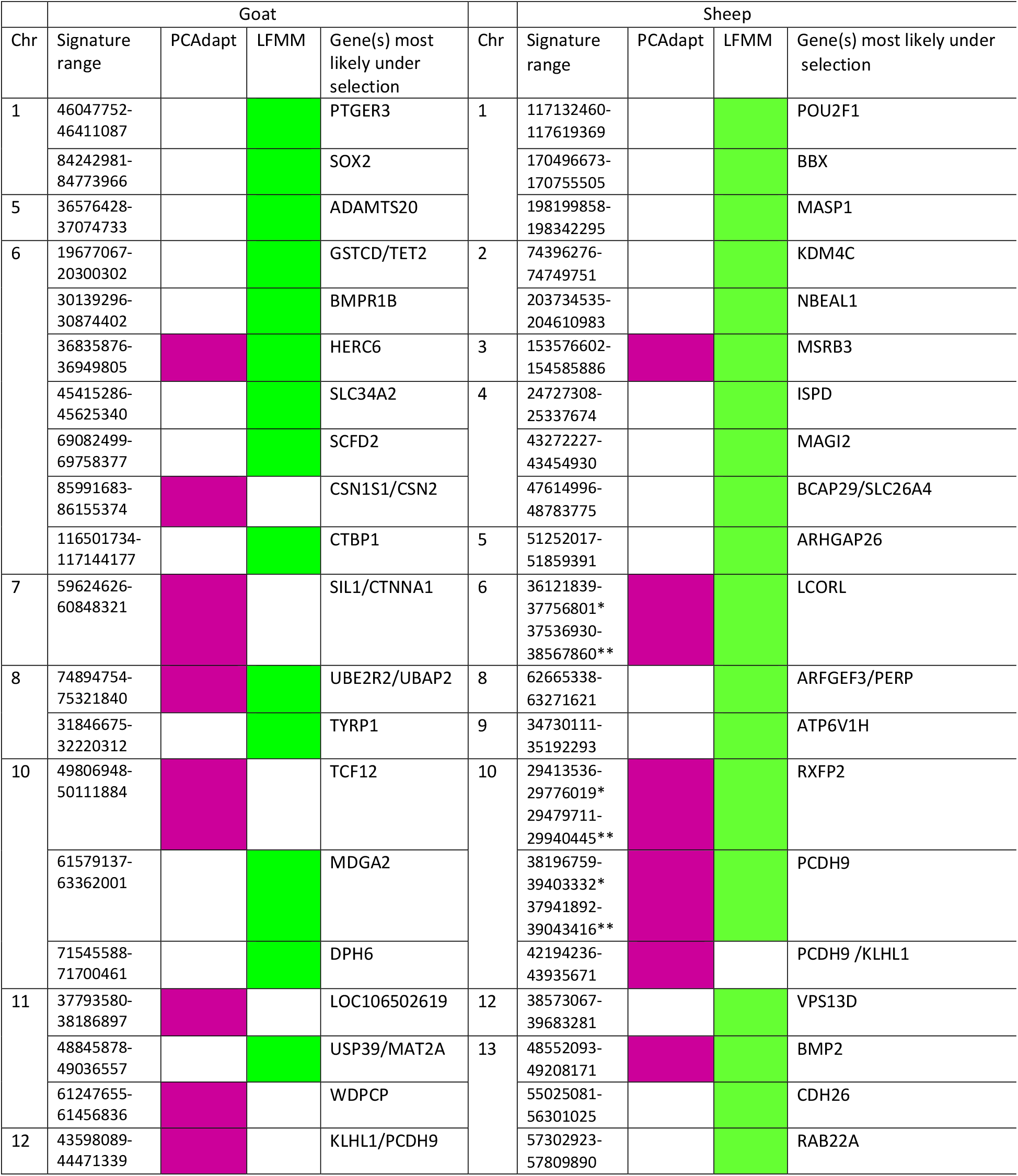
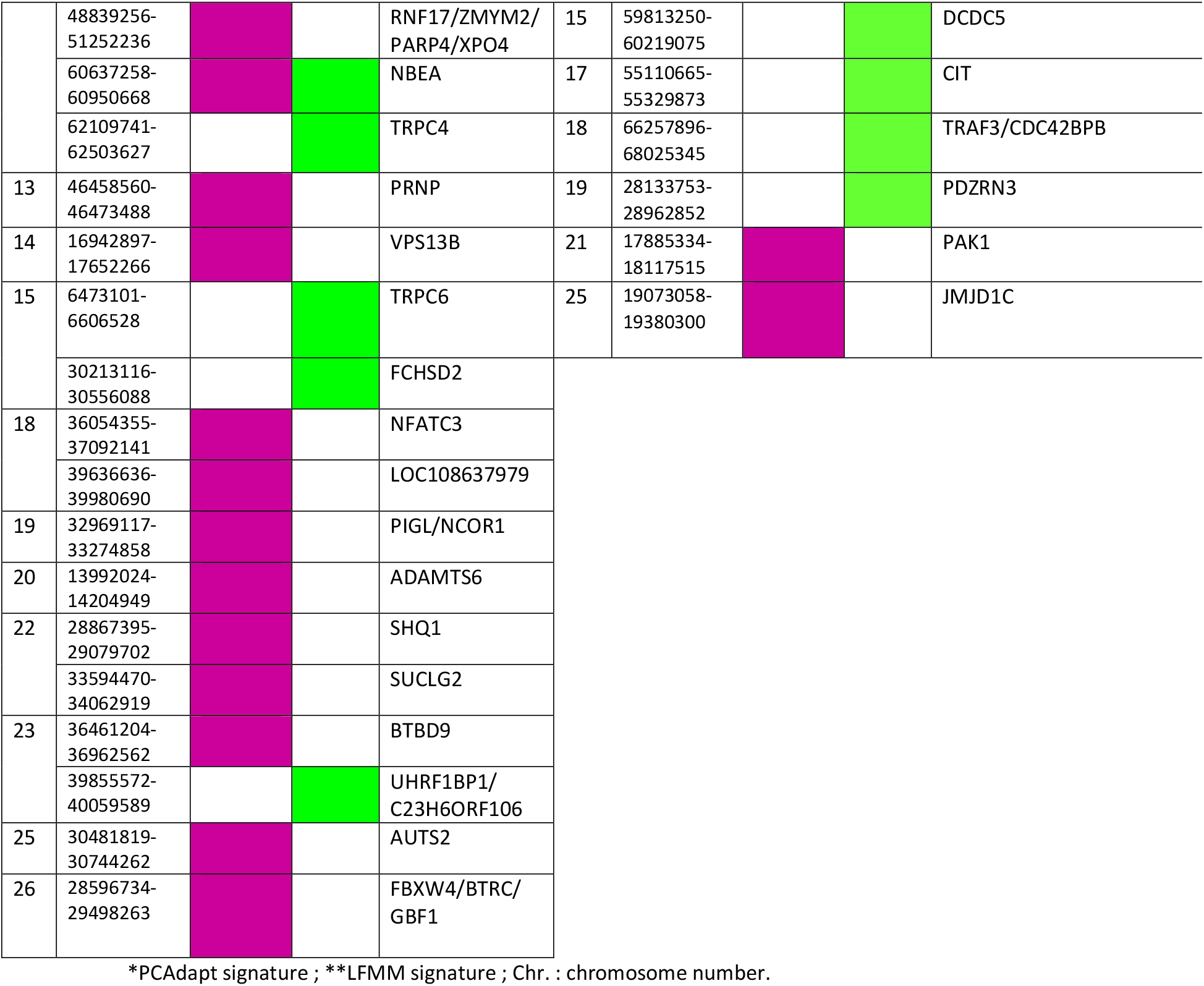
Selection signatures identified in goats and sheep by LFMM and PCAdapt using the cradle method. Each signature was named according to the gene(s) most likely targeted by the selection process, taking into account the distance of the SNPs from the genes, the associated p-values and the number of SNPs near or in the gene(s); All annotated genes associated with each selection signature are highlighted in supplementary tables 3 and 4.

The LFMM analysis was based on the PCA/HCPC ranking in which the cradles were clustered according to the similarity of their environments (Figure 2). The PCA analysis was largely driven by the first component (76.9% of variance retained by PC1). The cradles appeared thus distributed along a South-North axis, following a decreasing temperature and aridity gradient on one side, and an increasing altitudinal gradient on the other side. HCPC clustering, indicating K=5 as the optimal number of clusters, was as follows: (1) breeds from southern Italy, Sicily, southern Spain and the island of Majorca, *i.e*. regions with high annual mean temperatures and very low rainfall during the dry season. The group (2) clustered the Spanish (Blanca de Rasquera, RAS) and Italian (Garganica, GAR) breeds, raised under low rainfall also during the wet season. The third group (3) clustered Spanish, French and Italian goat breeds, in an intermediate position with regard to the aridity gradient during the dry season. The groups (4) and (5) corresponded to Italian breeds in regions of high altitude, low average temperatures and high precipitation (Figure 2). PCA/HCPC processed with environmental data recorded at the GPS area level (see Supplementary Figure 7), also showed five clusters that followed a temperature and precipitation gradient, but without altitude gradient (Supplementary Figure 7.3), which implied a different clustering than the one obtained by the cradle method (Figure 2) and a lower number of signatures highlighted by LFMM (Figure 1).

**Figure 2.**
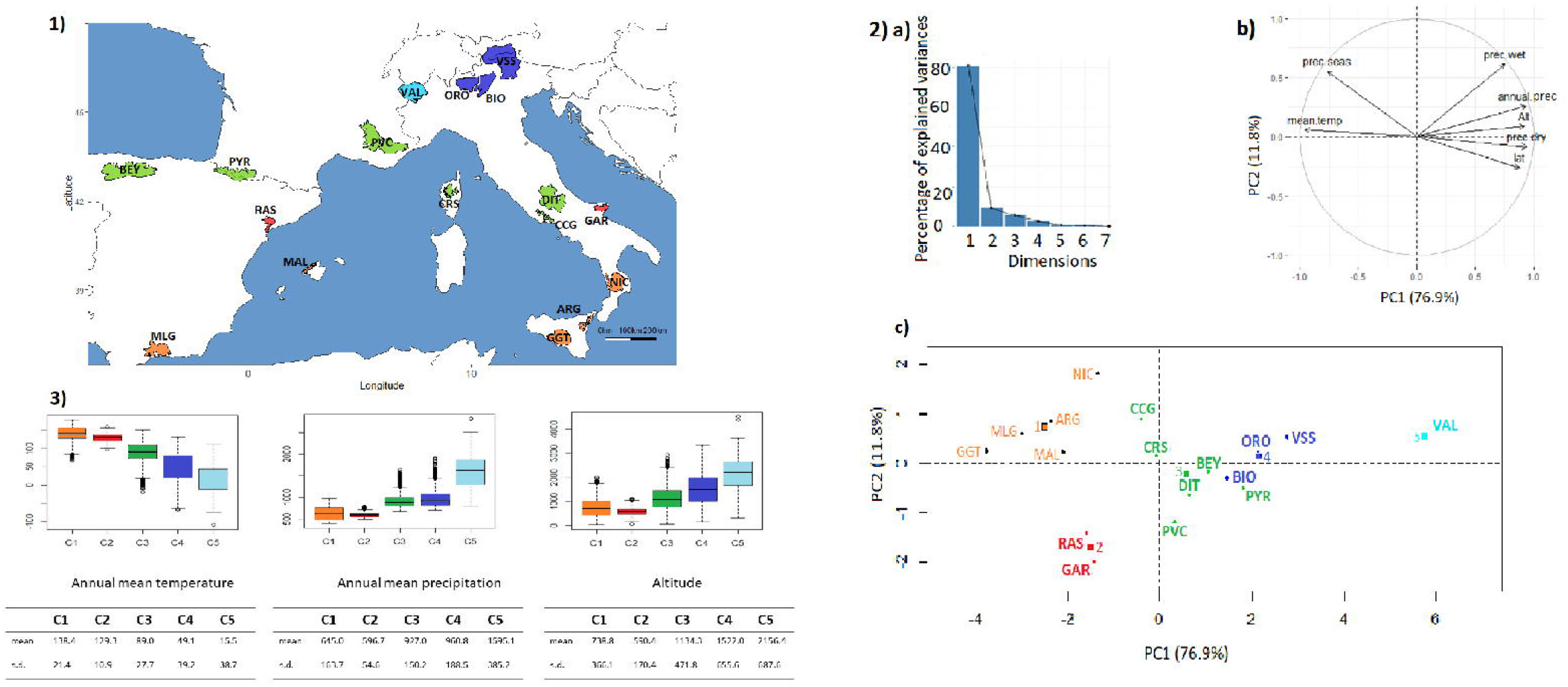
Goat cradles analyses as a function of environmental variables. 1) Mapping display of the cradles coloured according to the result of the clustering. 2). HCPC analysis. a) Scree plot. b) PCA correlation circle. c) PCA score plot of goat cradles. Colours represent the main clusters obtained by HCPC (K=5). PC 1 and PC 2 represent respectively principal components 1 and 2; numbers in brackets show the variance explained by each PC. 3) Boxplot representation for the variables Mean Annual Temperature, Mean Annual Precipitation and Altitude for the different HCPC groups and associated numerical values. Alt.: Altitude in meters, T°: Temperature in °C×10, prec.: precipitation in millimeters, min.: minimal, max.: maximal, s.d.: standard deviation.

The LFMM analysis, based on this cluster ranking (Figure 3), showed several strong signals near genes that previously have been implicated in adaptation: SOX2, ADAMTS20, UBE2R2/UBAP2, DPH6, NBEA, TRPC4, TRPC6, and UHRF1BP1/C23H6ORF106. Strong signals detected by PCAdapt analysis (Supplementary Table 6) and also found in the literature as implicated in adaptive processes, corresponded to the genes SIL1/CTNNA1, UBE2R2/UBAP2, SHQ1, SUCLG2 as well as a large area on chromosome 12 containing RNF17, ZMYM2, PARP4 and XPO4 and the vast intergenic area between KLH1 and PCDH9. A particularly strong signal was found at locus LOC108637979.

**Figure 3.**
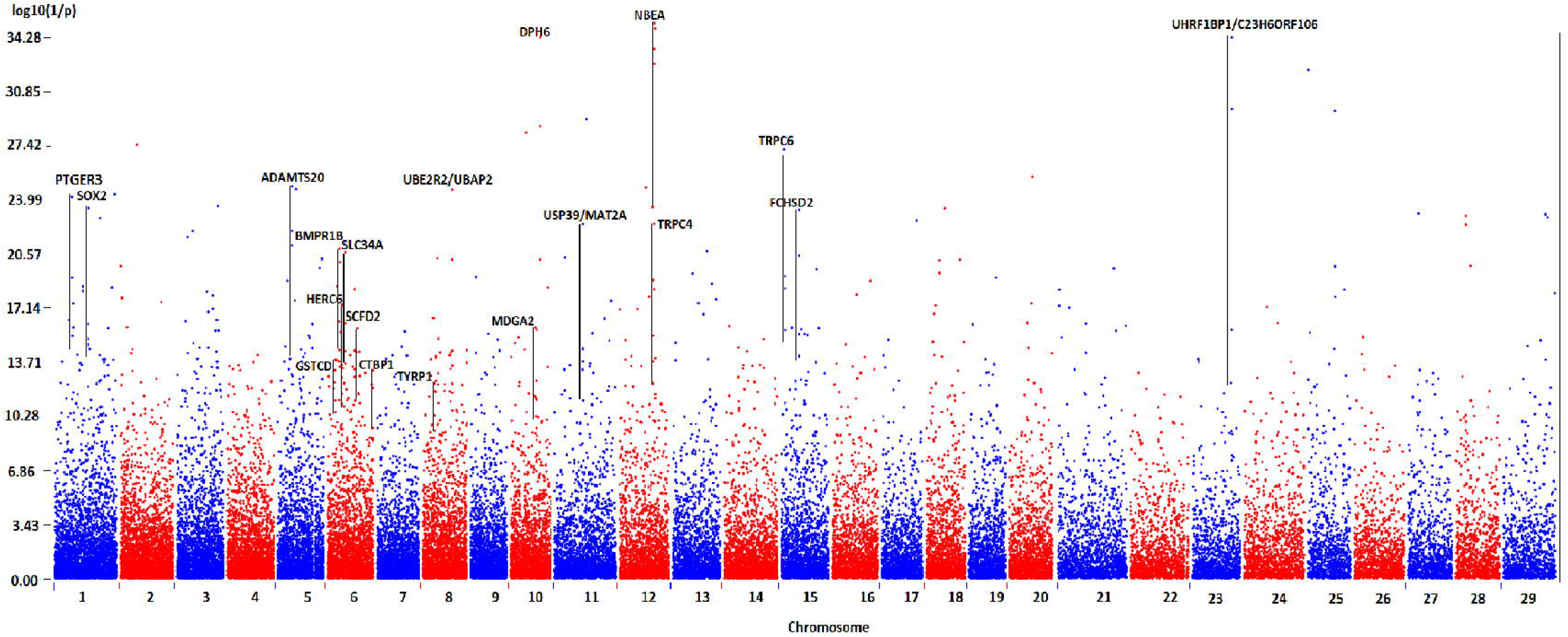
Genome scan for selection signatures in local goat breeds obtained by the LFMM approach and via the cradle method. Each signature was named according to the gene(s) most likely targeted by the selection process, taking into account the distance of the SNPs from the genes, the associated p-values and the number of SNPs near or in the gene(s).

A GO enrichment analysis of the genes identified, highlighted biological processes linked to transport via genes AQP3 and AQP7 and to gap junction assembly, via the genes CTNNA1 and GJB2 (Supplementary Figure 9).

### 4) Sheep dataset: track for selection signatures

From the 26 selection signatures, 8 were identified by PCAdapt, 23 by LFMM; 5 signatures were evidenced by both methods (table 1; Supplementary Table 5 and 7). These signatures were documented in the literature (Supplementary Table 4), all PCAdapt signatures and 18 out of 23 LFMM signatures could be linked to environmental adaptations.

Figure 4 shows the PCA/HCPC results used for the LFMM analysis. As in Figure 2 for goats, PC1 accounted for a large part of the variation (76.0%) and showed similar South-North gradients of temperature and aridity (Figure 4). The optimal K=5 generated the following clusters: group (1) concerned regions with highest aridity and temperature levels, including the Sicilian, Southern Italian, the island of Majorca and Central Spanish breeds. Groups (2) and (3) were in an intermediate position, in term of dryness and wetness. Group (2) corresponded to regions showing slightly higher average temperatures than group (3) and high levels of precipitation during the wet season (especially for Gallega GAL, Manech Tête Rousse MTR and Latxa LATX), while group (3) included regions of higher altitudes (except for Causse du Lot CDL). Groups (4) and (5), including the Pyrenean and Alpine breeds of the three countries, corresponded to areas of very high altitudes and significant precipitation levels (Figure 4). LFMM on the basis of this cluster ranking (Figure 5) showed several strong signals near genes that previously have been implicated in adaptation: MASP1, NBEA, MSRB3, RXFP2, BMP2.

**Figure 4.**
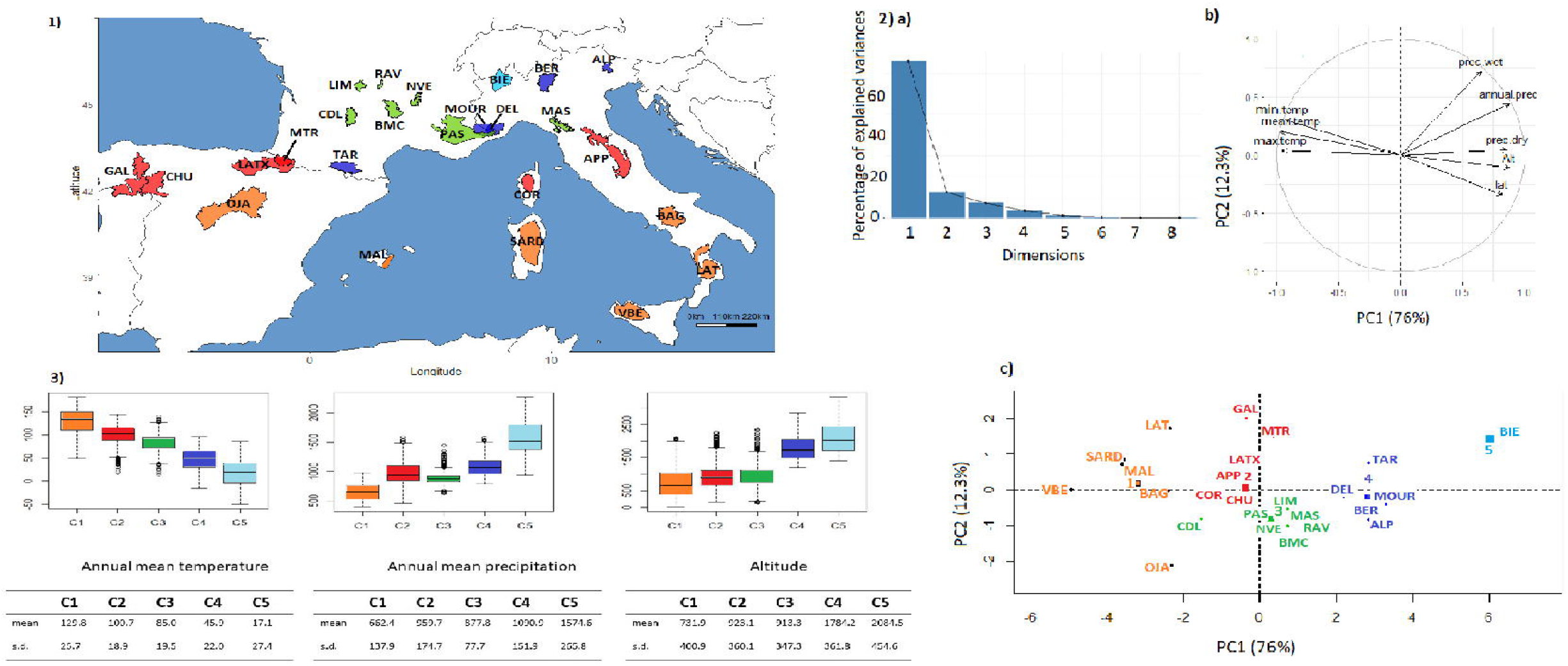
Sheep cradles analyses as a function of environmental variables. 1) Mapping display of the cradles coloured according to the result of the clustering. 2) HCPC analysis. a) Scree plot. b) PCA correlation circle. c) PCA score plot of goat cradles. Colours represent the main clusters obtained by HCPC (K=5). PC 1 and PC 2 represent respectively principal components 1 and 2; numbers in brackets show the variance explained by each PC. 3) Boxplot representation for the variables Mean Annual Temperature, Mean Annual Precipitation and Altitude for the different HCPC groups and associated numerical values. Alt.: Altitude in meters, T°: Temperature in °C×10, prec.: precipitation in millimeters, min.: minimal, max.: maximal, s.d.: standard deviation.

**Figure 5.**
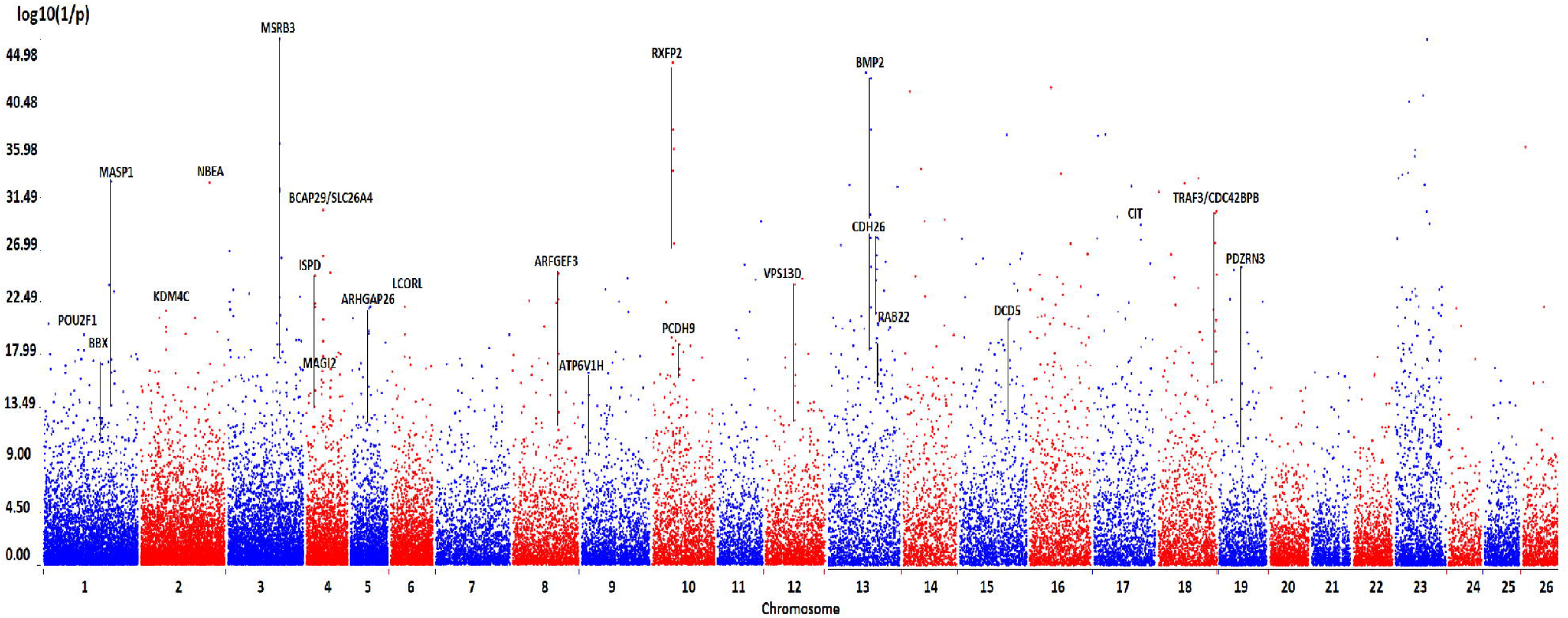
Genome scan for selection signature in local sheep breeds obtained by the LFMM approach and via the cradle method. Each signature was named according to the gene(s) most likely targeted by the selection process, taking into account the distance of the SNPs from the genes, the associated p-values and the number of SNPs near or in the gene(s).

The strongest PCAdapt signals (Supplementary Table 7) corresponded to: PAK1 and the KLHL1/PCDH9 intergenic region, both described previously as being involved in adaptation. A broad signal was on chromosome 6 and implied the genes HERC6, SPP1, LAP3, also previously involved in adaptation.

A GOrilla analysis of the candidate adaptation genes detected enriched biological processes linked to (i) heart development and regulation, (ii) cell growth regulation, (iii) cellular response to organic substance, and (iv) regulation of catalytic activity (Supplementary Figure 10).

### 5) Comparison of sheep and goat signatures

The genes NBEA, SHQ1 and VPS13 genes (goat VPS13B and sheep VPS13D), the regions containing HERC6 and ABCG2 as well as the KLHL1/PCDH9 intergenic region, appeared to be under selection, in both goat and sheep (Table 1, Supplementary Tables 3 and 4). Most of the selection signatures (74%), or genes included in the signatures, were found in the literature on mammals and birds as being related to environmental adaptations (Supplementary Tables 3 and 4). Noteworthy (Supplementary Tables 3 and 4), 16 goat and 14 sheep selection signatures were also identified in Chinese sheep adapted to extreme environments (Yang et al., 2016); four signatures in goats and three in sheep, were found in Ethiopian sheep kept at high-altitude (Edea et al., 2019). Finally, three goat and six sheep signatures were detected in Egyptian spatially differentiated sheep and goats in arid environments (Kim et al., 2016).

## DISCUSSION

This study proposed and tested a new genomic landscape approach for the identification of selection signatures in livestock breeds. The key points are (i) filtering the breeds to be included in the study and (ii) matching the genotypes to the environmental conditions recorded not at the current coordinates of the individuals, but at the level of the areas identified as their cradle of origin.

### 1) Performance of cradle Versus GPS method

We targeted French, Spanish and Italian breeds with strong historical links to their territory of origin, in order to build original datasets. For most of the breeds, this link was attested for several centuries. Most of the total of 63 signatures we identified, had not been detected in the initial studies (Manunza et al., 2016; Fariello et al., 2014; Rochus et al., 2018; Bertolini et al., 2018; Ruiz-Larrañaga et al., 2018; Oget, Servin & Palhière, 2019). A high proportion (74%) of the highlighted selection signatures were previously identified by studies on environmental genetic adaptation in birds and mammals, from different parts of the world, providing a validation of our method. Further corroboration came from the many signatures identified in our study that have been previously linked, in sheep or goats from different parts of the world, to extreme environmental adaptations (Yang et al., 2016; Edea et al., 2019; Kim et al., 2016; Wang et al., 2016; Supplementary Tables 3 and 4 for a complete list).

Our methodological tests, allowed highlighting a few points. First, the vast majority of small ruminant breeds have been shaped by traditional transhumant breeding practices. However, the individual GPS method will only be able to consider summer grazing areas, (i) for breeds and herds always managed according to these practices (ii) and that, only if the sampling is carried out during the summer period. Consistent with this, we observed that GPS methods gave poor results with the altitude variable, whereas with the cradle method it was possible to detect 53% of all signatures, by the consideration of this single variable. Secondly, considering the variable, mean annual temperature, (i) it appeared less sensitive to the location method than the altitude variable, and (ii) the correlation of genotypes with environmental variables recorded at the level of the area defined by the set of the breed’s GPS coordinates (GPS area method), yielded better results than considering the environmental variable at individual GPS coordinates (individual GPS method). This demonstrated the benefits of smoothing environmental data at the area level rather than considering it at individual points. Finally, the greatest number of selection signatures was scored by considering all the environmental variables at the same time and using the cradle method including transhumant pasture areas.

Thus, the strengths of our novel cradle method are: (i) taking into account the history of the breeds avoids including breeds, which could interfere with the signal (*i.e*. breeds which would not have a strong link with a given environment or whose link would have been distorted). Moreover, (ii) in industrialized countries, agricultural intensification in recent decades led to major shifts in practices, particularly the abandonment of transhumance, and also significantly changed the distribution areas of herds. Considering the cradle of the breed rather than the current distribution thus links with more accuracy, the genetic determinants of adaptation to the environment that has mainly shaped the breed’s genome. (iii) Our methodological framework combined the PCAdapt algorithm, which is particularly suited to heterogeneous populations (Luu, Bazin & Blum, 2017), such as local breeds, with LFMM, which identifies the links between genome and environment. PCAdapt approach followed by elimination of signatures related to agronomic selections on the basis of the literature, indicated potential environmental adaptations, while the LFMM approach detected areas of the genomes that followed the environmental gradients of aridity and altitude characterizing the studied datasets.

It should be noted that the breeds, considered today, come from ancestral populations which life history is largely unknown. Indeed, the various traces and historical documents indicating the existence of breeds on territories for several centuries, or even millennia, refer to the ancestral populations from which they originated. Hence, selection signatures identified may predate the formation of the breed or conversely, reflect rapid recent changes. In addition, environmental features of a cradle may not be unique and may also have changed. The environmental characterization was inferred from climatic data recorded during the 20th century. This constitutes a bias, which in the context of current climate change, must be carefully considered. We expect that our work will contribute to the synthesis of Rellstab et al. (2015) by focusing on the particularities of livestock at the interface of natural and anthropized environment.

### 2) Selection signatures identified in this study

The highlighted selection signatures can be categorized according to physiology and phenotypes. In Supplementary Text 2, we discussed the genes that seem to be involved (i) in lipid storage, (ii) in seasonal patterns and circadian behaviors, (iii) in coat colour and horns, (iv) in hypoxia and/or heat stress response, (v) in immunity, (vi) in lung function, and (ii) in neuronal function. We also discussed selection signatures that seemed difficult to classify because of their length and the diversity of the contained genes. Here we focus on the most robust signals with the most relevant information from the previous studies.

#### a) Goat selection signatures

We identified SUCLG2, in goat by the PCAdapt approach. It was found under selection in sheep in arid Egyptian environments (Kim et al., 2016) and in the Chinese desert (Yang et al., 2016), but also in yaks adapted to extreme altitudes (Qi et al., 2019). The gene is involved in the propanoate metabolic pathway and was significantly associated with growth in pigs (Yang et al., 2012). Moreover, the latter study suggested that SUCLG2 plays a key role in the regulation of POU1F1, which is well known to be involved in growth function, and belongs to the same family of POU2F1, detected by our study, in sheep. The study by Schmidt et al. (2017) in the blind subterranean mole rat, known to show remarkable tolerance to hypoxia and cancer resistance, also highlighted SUCLG2 and POU1F1 in the liver transcriptome. It was hypothesized that the energy-saving responses triggered in hepatic metabolic pathways may be crucial adaptations to low oxygen levels. Finally, the study of Tian et al. (2017) assessing energy metabolism-related genes in hypoxia-tolerant mammals, identified SUCLG2 as significantly differentially expressed in the liver of cetaceans.

SOX2 and DPH6 were found correlated to the environmental gradient in goats by LFMM. In mice, SOX2 is expressed in adult SCN neurons and positively regulates transcription of the core clock gene, Period2, implicated in behavioral rhythms linked to environmental light cycles (Cheng et al., 2019). A link between SOX2 and cold adaptation has been found in marmots (Bai et al., 2019). Interestingly, BBX another Sox proteins that belongs to the HMG box superfamily of DNA-binding ones, was identified by us, in sheep. DPH6 belongs to the circadian rhythm-related GO categories. Moreover it was found under selection in northern European cattle (Stronen et al., 2019), in yaks (Lan et al., 2018), and associated with plateau conditions in Chinese sheep (Yang et al., 2016).

Both PCAdapt and LFMM highlighted the signature near UBE2R2/UBAP2 and bordered by AQP7/AQP3. UBE2R2/UBAP2, have been found to be associated with hypoxia tolerance in humans (Udpa et al., 2014) and with highlands in Chinese sheep (Yang et al., 2016). AQP7 was linked to high altitudes in yaks (Qi et al., 2019). The aquaporins AQP7 and AQP3, which belong to the hypoxia-related genes, have been identified in thermal adaptation through the regulation of evapotranspiration and cryoprotectant transport (Wollenberg Valero et al., 2014).

Interesting to note that TRPC4 gene, on chromosome12, and TRPC6 gene, on chromosome 15, are among the strongest signatures highlighted by the LFMM approach. Transient receptor potential channel (TRPC) proteins have been characterized as molecular substrates mediating receptor-activated cation influx. TRPC4 and TRPC6, in particular, have been shown to strongly contribute to synaptic information transfer in neuronal dendrites via the Ca^2+^-dependent release of neurotransmitter (Chen et al., 2017). The increase in dendritic γ-aminobutyric acid (GABA) release from thalamic interneurons appears critically dependent on these TRP proteins (Munsch et al., 2003). Hence, these genes may mediate hot and cold sensation and affect endothelial dependent regulation of vascular tone (Duan et al., 2018). Moreover, there is strong evidence for an important function of TRPC6 in pulmonary vascular (Malczyk et al., 2017), and a functional role in hypoxia induction, potentially via the metabolism regulation of the HIF-1α (Wang et al, 2016; Li et al., 2015; Xu et al., 2014). TRPC6 was associated with high altitude adaptation in Tibetan highlanders (Deng et al., 2019) and TRPC4 was implicated in body temperature regulation in cattle (Howard et al., 2013).

#### b) Sheep selection signatures

Both PCAdapt and LFMM approach identified the gene MSRB3. This gene has been subject to high selection pressure in sheep (Kijas et al., 2012) and is commonly assumed to be involved in ear morphology. It has been reported in large-eared sheep (Wei et al., 2015). Its link with ears was also highlighted in pig (Chen et al., 2018) and dogs (Webster et al., 2015). Our study showed a correlation between SNPs near this gene and the environmental gradient. Interestingly, a recent study (Mastrangelo et al., 2019) associated genetic variation in this gene and fat deposition in sheep. Furthermore, Webster et al. (2015) revealed that this gene was in genetic linkage with variants in HMGA2 governing body mass in dogs and sheep (Kijas et al. 2012). HMGA2 and MSRB3 are neighbors on sheep chromosome 5. Moreover, HMGA2 is known to be involved in adipose tissue, development and obesity in mouse (Xi et al., 2016). The MSRB3 gene appears to be of central importance in terms of adaptation, because it was identified under selection in highland Ethiopian sheep (Edea et al., 2019), in Tibetan sheep (Wei et al., 2016), in Tibetan Yaks (Qi et al., 2019) and in Tibetan dogs (Gou et al., 2014).

Like MSRB3, both methods (PCAdapt and LFMM) implicated BMP2, which initiates osteoblast and adipocyte differentiation. It has previously been found in association with dry conditions in Chinese (Yang et al., 2016), and Egyptian sheep (Kim et al., 2016). It was also found under selection at high altitude, in Tibetan sheep (Wei et al., 2016) and in Yaks (Qi et al., 2019). Adaptive mechanisms governed by BMP2 could be linked to lipid storage capacities, particularly at the level of the tail (Zhu et al., 2019; Yuan et al., 2017; Mastrangelo et al., 2019).

Finally, RXFP2, well known to be under strong selection due to its role in horn development (Johnston et al., 2011; Allais-Bonnet et al., 2013; Kijas et al., 2012; Fariello et al., 2014), was identified by both PCAdapt and LFMM approaches. It is the receptor of the peptide hormone INSL3 and plays a regulatory role in testis and ovaries of various mammalian species (Hanna et al., 2010). In the seasonally breeding roe deer, with a periodic variation in the volume of the testicles, the seasonal expression of INSL3/RXFP2 was correlated with the differentiation of Leydig cells in the testis (Hombach-Klonisch et al., 2004). Interestingly, RXFP2 was associated with latitude, precipitation and temperature in free-ranging bighorn sheep (Roffler et al., 2016); and identified as being under selection in Tibetan sheep (Wei et al., 2016). Moreover, RXFP2, as MSRB3 and SLC26A4 (also identified by our study) showed signs of rapid evolution in semi-feral breeds’ sheep (Pan et al., 2018). This suggests an adaptive role in the wild or, potentially, in traditional livestock farming with limited human intervention.

#### c) Selection signatures highlighted in both species

An interesting result concerns the area between PCDH9 and KLHL1 gene that was found under selection in both sheep and goats. This region appears to be a genetic desert that extends over a syntenic segment conserved on bovine chromosome 12, goat chromosome 12 and sheep chromosome 10. Kim et al. (2016) also found this genomic region to be under selection in both sheep and goats that are adapted to arid environments, and suggested a major role for this region. PCDH9 has been found to be implicated in autism disorders (Oksenberg et al., 2013). Protocadherins are thought to be implicated in various aspects of neuronal development and functions. PCDH9 in particular is involved in synaptic cell adhesion (Hayashi & Takeichi, 2015; Seong, Yuan, & Arikkath, 2015). KLHL1 was found related to neuron motion and neuromuscular process (Shin et al., 2014), and it may play a role in organizing the actin cytoskeleton of the brain cells.

Another selection signature common to both species and most interesting is the NBEA signal. NBEA was reported to be associated with high altitude in Ethiopian sheep (Edea et al., 2019), Chinese sheep (Yang et al., 2016), cattle at high altitude (Zeng, 2017), and yaks (Qi et al., 2019). Furthermore, it was previously associated to body temperature regulation in cattle (Howard et al., 2013) and was found under selection in Ugandan and Moroccan goats (Onzima et al., 2018; Benjelloun, 2015). NBEA is a brain specific A-kinase anchor protein (AKAP), which is required for synaptic surface expression of glutamate and GABA receptors. Neurons lacking the BEACH (beige-Chediak/Higashi) domain protein Neurobeachin (NBEA) show strongly reduced synaptic responses caused by a reduction in surface levels of glutamate and GABAA receptors. Hence, NBEA plays an essential role in thermal adaptation through the regulation of synaptic transmission (Farzana et al., 2016; Nair et al., 2013). In the present study, LFMM revealed a correlation of NBEA and the environmental gradient, in both goats and sheep; in goat it was also identified by PCAdapt.

Interestingly, Alberto et al. (2018) identified NBEA, but also, HERC6 and SLC34A2, all detected by us, involved in selective sweeps that differentiate domestic from wild sheep and goat populations, indicating a predomestic selection on these genomic areas.

Finally, the signature near NBEA is bordered in sheep, by BMPR2, for which mutations have been associated with high Altitude Pulmonary Hypertension (APH), in Kyrgyz Highlanders (Iranmehr et al., 2019) and also in cattle (Newman et al., 2011). Moreover, this gene was found associated with desert and plateau conditions in Chinese sheep (Yang et al., 2016). BMP2, another BMPs, belonging to the TGF-β superfamily and able to activate BMPR2, was also identified by us in sheep, as a major signal (see above). Hypoxia may act on BMPR2 activity (Yang et al., 2019) and on BMP2 signaling in the pulmonary vasculature (Anderson et al., 2009).

## CONCLUSION

Our study provided a methodology for integrating ecological, historical and cultural approaches in the search for selection signatures in small ruminants, which as domestic animals, are at the very interface of natural and anthropized environments. One of the key step is the characterization of the environment at the cradle level, thus targeting the geographic area, including transhumant summer pastures, where the breed has evolved by traditional practices. Local breeds are derived from populations selected for centuries and sometimes millennia; they have thus developed singular links between genome and environment. Today, local breeds are increasingly neglected and have become endangered or have even disappeared. For those that remain, traditional practices, including the transhumance that has been one of the main driving forces in their development, are being abandoned (Caballero et al., 2011; Collantes, 2009). This study emphasizes that local breeds are invaluable resources of environmental adaptation in the context of climate changes.

## DATA ACCESSIBILITY

Data availability: For goats, we used the AdaptMap dataset, shared on Dryad: https://doi.org/10.5061/dryad.v8g21pt. For sheep, we used the French dataset online at the Zenodo repository 10.5281/zenodo.237116, and the Italian dataset partially available in the Figshare repository via https://doi.org/10.23644/uu.8947346 (for the few missing breeds please contact Elena Ciani: and Johannes A. Lenstra).

## AKNOWLEDGMENTS

The authors would like to thank the AdaptMap consortium; Vincenzo Landi, Philippe Teinturier, Claire Jouannaux and the “Collectif CORAM” for their valuable insights into the Spanish and French local breeds. The calculations presented in this article were performed on (i) the CALI calculator of the University of Limoges, funded by the Limousin region, the European Union, the XLIM institutes, IPAM, GEIST, and the University of Limoges, and on (ii) the genotoul bioinformatics platform Toulouse Occitanie (Bioinfo Genotoul, doi: 10.15454/1.5572369328961167E12). This work was supported by the French region Nouvelle-Aquitaine (PAPAGENO project, 2016-2018), and the AAP CNRS 2020: “Adaptation du Vivant à son Environnement” (EMPREINTES project, 2020-2021).

## AUTHOR CONTRIBUTIONS

B.S. and M.C.: contributed to data analyses; P.V.: performed anthropological analyses; D. T.- M.: performed historical analyses; E.C., B.S., C.M.-R and G.T.-K: acquisition of data; M.B. and E.R.: contributed to the conception of the study; J.A.L.: acquisition of data / review & editing; S.J.G.H.: review and editing; F.P.: contributed to the conception of the study / review & editing; B.B.: contributed to data analyses / review & editing; A.D.S.: designed research / analyzed data / wrote the paper; all authors contributed to the final manuscript.

## SUPPLEMENTARY MATERIAL

Supplementary information (see pdf document)

Supplementary data (excel document). Summarize of environmental data extracted at the cradle level for the sheep and goat datasets, and extracted at the level of the GPS area for the goat dataset; PCA dimensions; s.d.: standard deviation.

## Notes

### Competing Interest Statement

The authors have declared no competing interest.

