## SupplementalFiles for "Local adaptations of Mediterranean sheep and goats through an integrative approach"

### Table of Contents:

| | |
|---|---|
| **Supplementary Table 1.** Initial list breeds for the constitution of the datasets. | Page 1 |
| **Supplementary Figure 1.** Synthetic schema of the main steps for the proposed approach. | Page 2 |
| **Supplementary Figure 2.** Goat and sheep cradles description as a function of environmental variables: Synthetic descriptions of cradles and some characteristic breed phenotypic traits. | Pages 3-4 |
| **Supplementary Table 2.** Description of the breeds included in the sheep and goat datasets. Number of individuals considered, country of origin, information on the breed history, geographical definition of the cradle, description of the breed, use and status. | Pages 5-16 |
| **Supplementary Figure 3.** LD analyses for sheep and goats. | Page 17 |
| **Supplementary Text 1.** Genetic structure assessment: admixture analyses, sNMF cross-entropies and Mantel tests. | Pages 18-20 |
| **Supplementary Figure 4.** Mapping display for sheep and goats, of the geographical cradles and the GPS coordinates of the sampled points; statistical comparisons for the variables Annual Mean Temperature, Annual Mean Precipitation and Altitude between the distributions obtained by the cradle method and the GPS area method, display of the results via boxplots. | Pages 21-22 |
| **Supplementary Figure 5.** Display of the selection signatures obtained with the goat dataset for the different methods tested, *i.e.* cradle method, GPS area method and individual GPS method, using the LFMM approach and | Pages 23-25 |

| | |
|---|---|
| considering individually the variables: Annual Mean Temperature, Annual Mean Precipitation and Altitude | |
| **Supplementary Figure 6.** Display of the selection signatures obtained on the goat dataset for the cradle method and the GPS area method, using the LFMM approach; cradles and GPS areas were characterized by all the environmental variables available; a PCA followed by an HCPC allowed to group areas according to their similarity in terms of environment. | Page 26 |
| **Supplementary Figure 7.** Goat "GPS areas" analyzed as a function of environmental variables. 1) Mapping display of the "GPS areas" coloured according to the result of the clustering. 2) Hierarchical Clustering on Principal Component (HCPC) analysis. a) Scree plot. b) PCA correlation circle. c) PCA score plot of goat cradles. Colours represent the main clusters obtained by HCPC for K=5. PC 1 and PC 2 represent respectively principal components 1 and 2; numbers in brackets show the variance explained by each PC. 3) Boxplot representation for the variables Annual Mean Temperature, Annual Mean Precipitation and Altitude for the different groups and associated numerical values. Alt.: Altitude in m (meters), T°: Temperature in °C×10, prec.: Precipitation in mm (millimeters), min.: minimal, max.: maximal, s.d.: standard deviation. | Page 27 |
| **Supplementary Figure 8.** PCAdapt scree plots, a) considering the goat dataset, b) considering the sheep dataset. | Page 28 |
| **Supplementary Table 3.** Selection signatures identified in goats according to the LFMM and PCAdapt approach and using the cradle method: detailed literature review. | Pages 29-41 |
| **Supplementary Table 4.** Selection signatures identified in sheep according to the LFMM and PCAdapt approach and using the cradle method: detailed literature review. | Pages 42-53 |
| **Supplementary Table 5**. Details concerning the selection signatures identified in goats and sheep: name and position of the SNPs. | Page 54-69 |
| **Supplementary Table 6**. Display of the selection signatures identified in the goat dataset. The figures have been modified from outputs obtained via NCBI Genome Data Viewer (https://www.ncbi.nlm.nih.gov/genome/gdv/): ARS1. SNPs under selection (see their name and position in Supplementary Table 5) are symbolized by stars; red stars = SNPs identified by PCAdapt (and plots associated), green stars = SNPs identified by LFMM. | Page 70-86 |
| **Supplementary Table 7.** Display of the selection signatures identified in the sheep dataset. The figures have been modified from outputs obtained via NCBI Genome Data Viewer (https://www.ncbi.nlm.nih.gov/genome/gdv/): Oar_v3.1. SNPs under selection (see their name and position in Supplementary Table 5) are symbolized by stars; red stars = SNPs identified by PCAdapt (and plots associated), green stars = SNPs identified by LFMM. | Page 87-99 |
| **Supplementary Figure 9.** Gorilla enrichment analysis for goats. The colored areas correspond to the identified processes. | Page 100 |

| | |
|---|---|
| **Supplementary Figure 10.** Gorilla enrichment analysis for sheep. The colored areas correspond to the identified processes. | Page 101 |
| **Supplementary Text 2**. Discussion and classification of identified genes according to the literature. | Page 102-115 |

| Goat breed | country |
|---|---|
| Alpine | France |
| Angora | France |
| Corse | France |
| Fosses | France |
| Poitevine | France |
| Provencale | France |
| Pyrenean | France |
| Saanen | France |
| Alpine | Italy |
| Argentata | Italy |
| Aspromontana | Italy |
| Bionda dell'Adamello | Italy |
| Ciociara Grigia | Italy |
| Di Teramo | Italy |
| Garganica | Italy |
| Girgentana | Italy |
| Jonica | Italy |
| Maltese sarda | Italy |
| Maltese | Italy |
| Nicastrese | Italy |
| Orobica | Italy |
| Rossa Mediterranea | Italy |
| Saanen | Italy |
| Sarda | Italy |
| Valdostana | Italy |
| Valpassiria | Italy |
| Bermeya | Spain |
| Mallorquina | Spain |
| Malaguena | Spain |
| Murciano-Granadina | Spain |
| Palmera (Canaria island breed) | Spain |
| Blanca de Rasquera | Spain |

| Sheep breed | country |
|---|---|
| Mourerous Alpes du Sud | France |
| Mérinos d'Arles | France |
| Lacaune (lait) | France |
| Blanche du Massif Central | France |
| Berrichon du Cher | France |
| Romane | France |
| Rouge de l'Ouest | France |
| Mérinos de Rambouillet | France |
| Tarasconnaise | France |
| Rava | France |
| Noire du Velay | France |
| Limousine | France |
| Lacaune (viande) | France |
| Préalpes du Sud | France |
| Manech tête rousse | France |
| Corse | France |
| Romanov | France |
| Suffolk | France |
| Vendéen | France |
| Charollais | France |
| Causses du Lot | France |
| Ouessant (Bretagne) | France |
| Charmoise | France |
| Roussin de la Hague | France |
| Île-de-France | France |
| Texel | France |
| Altamurana | Italy |
| Comisana | Italy |
| Leccese | Italy |
| Sarda | Italy |
| Appenninica | Italy |
| Sopravissana | Italy |
| Gentile Di Puglia | Italy |
| Delle Langhe | Italy |
| Pinzirita | Italy |
| Bergamasca | Italy |
| Bagnolese | Italy |
| Laticauda | Italy |
| Alpagota | Italy |
| Altamurana | Italy |
| Massese | Italy |
| Fabrianese | Italy |
| Ripollesa | Spain |
| Xisqueta | Spain |
| Roja Mallorquina | Spain |
| Canaria de pelo | Spain |
| Gallega | Spain |
| Segureña | Spain |
| Castellana | Spain |
| Churra | Spain |
| Ojalada | Spain |

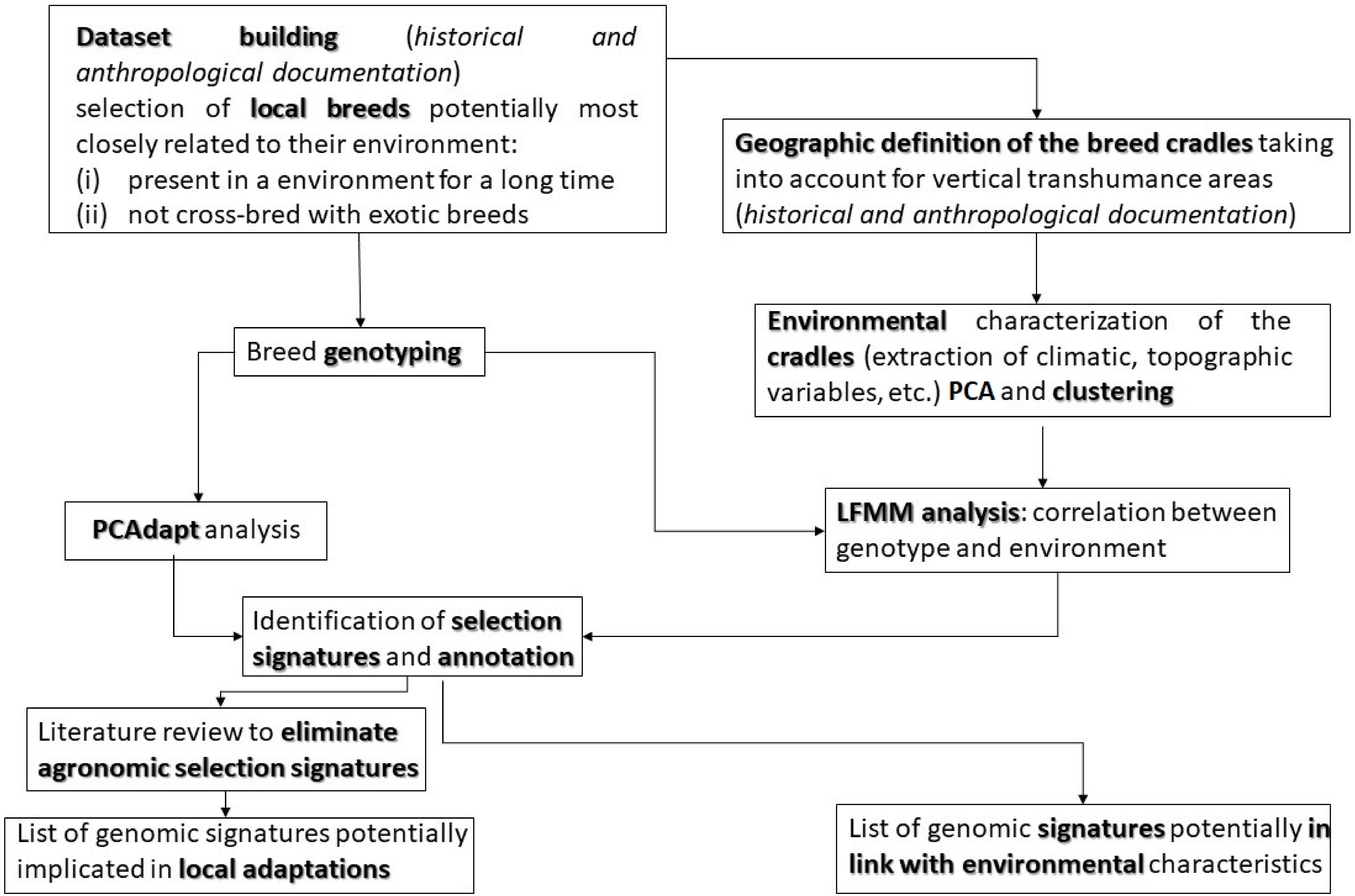
Dataset building (historical and anthropological documentation)
selection of local breeds potentially most closely related to their environment:
(i) present in a environment for a long time
(ii) not cross-bred with exotic breeds
Geographic definition of the breed cradles taking into account for vertical transhumance areas (historical and anthropological documentation)
Breed genotyping
Environmental characterization of the cradles (extraction of climatic, topographic variables, etc.) PCA and clustering
PCAdapt analysis
LFMM analysis: correlation between genotype and environment
Identification of selection signatures and annotation
Literature review to eliminate agronomic selection signatures
List of genomic signatures potentially implicated in local adaptations
List of genomic signatures potentially in link with environmental characteristics

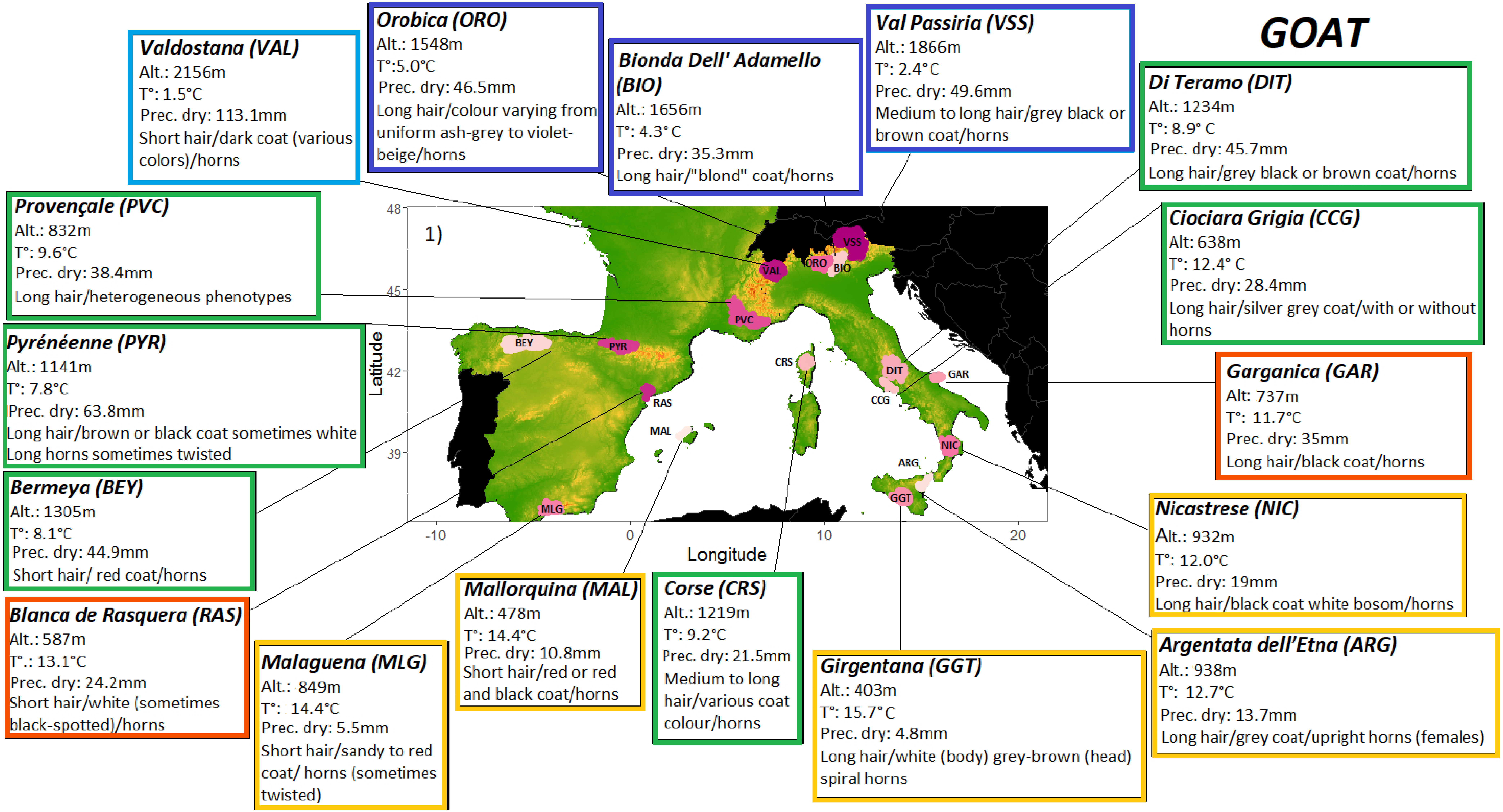
GOAT
Valdostana (VAL)
Alt.: 2156m
T°: 1.5°C
Prec. dry: 113.1mm
Short hair/dark coat (various colors)/horns
Orobica (ORO)
Alt.: 1548m
T°:5.0°C
Prec. dry: 46.5mm
Long hair/colour varying from uniform ash-grey to violet-beige/horns
Bionda Dell' Adamello (BIO)
Alt.: 1656m
T°: 4.3° C
Prec. dry: 35.3mm
Long hair/"blond" coat/horns
Val Passiria (VSS)
Alt.: 1866m
T°: 2.4° C
Prec. dry: 49.6mm
Medium to long hair/grey black or brown coat/horns
Di Teramo (DIT)
Alt.: 1234m
T°: 8.9° C
Prec. dry: 45.7mm
Long hair/grey black or brown coat/horns
Provençale (PVC)
Alt.: 832m
T°: 9.6°C
Prec. dry: 38.4mm
Long hair/heterogeneous phenotypes
Ciociara Grigia (CCG)
Alt: 638m
T°: 12.4° C
Prec. dry: 28.4mm
Long hair/silver grey coat/with or without horns
Pyrénéenne (PYR)
Alt.: 1141m
T°: 7.8°C
Prec. dry: 63.8mm
Long hair/brown or black coat sometimes white
Long horns sometimes twisted
Garganica (GAR)
Alt: 737m
T°: 11.7°C
Prec. dry: 35mm
Long hair/black coat/horns
Bermeya (BEY)
Alt.: 1305m
T°: 8.1°C
Prec. dry: 44.9mm
Short hair/ red coat/horns
Nicastrese (NIC)
Alt.: 932m
T°: 12.0°C
Prec. dry: 19mm
Long hair/black coat white bosom/horns
Blanca de Rasquera (RAS)
Alt.: 587m
T°.: 13.1°C
Prec. dry: 24.2mm
Short hair/white (sometimes black-spotted)/horns
Malaguena (MLG)
Alt.: 849m
T°: 14.4°C
Prec. dry: 5.5mm
Short hair/sandy to red coat/ horns (sometimes twisted)
Mallorquina (MAL)
Alt.: 478m
T°: 14.4°C
Prec. dry: 10.8mm
Short hair/red or red and black coat/horns
Corse (CRS)
Alt.: 1219m
T°: 9.2°C
Prec. dry: 21.5mm
Medium to long hair/various coat colour/horns
Girgentana (GGT)
Alt.: 403m
T°: 15.7° C
Prec. dry: 4.8mm
Long hair/white (body) grey-brown (head) spiral horns
Argentata dell'Etna (ARG)
Alt.: 938m
T°: 12.7°C
Prec. dry: 13.7mm
Long hair/grey coat/upright horns (females)
1)
VSS
ORO
BIO
VAL
PVC
BEY
PYR
CRS
DIT
GAR
CCG
RAS
MAL
NIC
ARG
GGT
MLG
48
45
42
39
Latitude
-10
0
10
20
Longitude

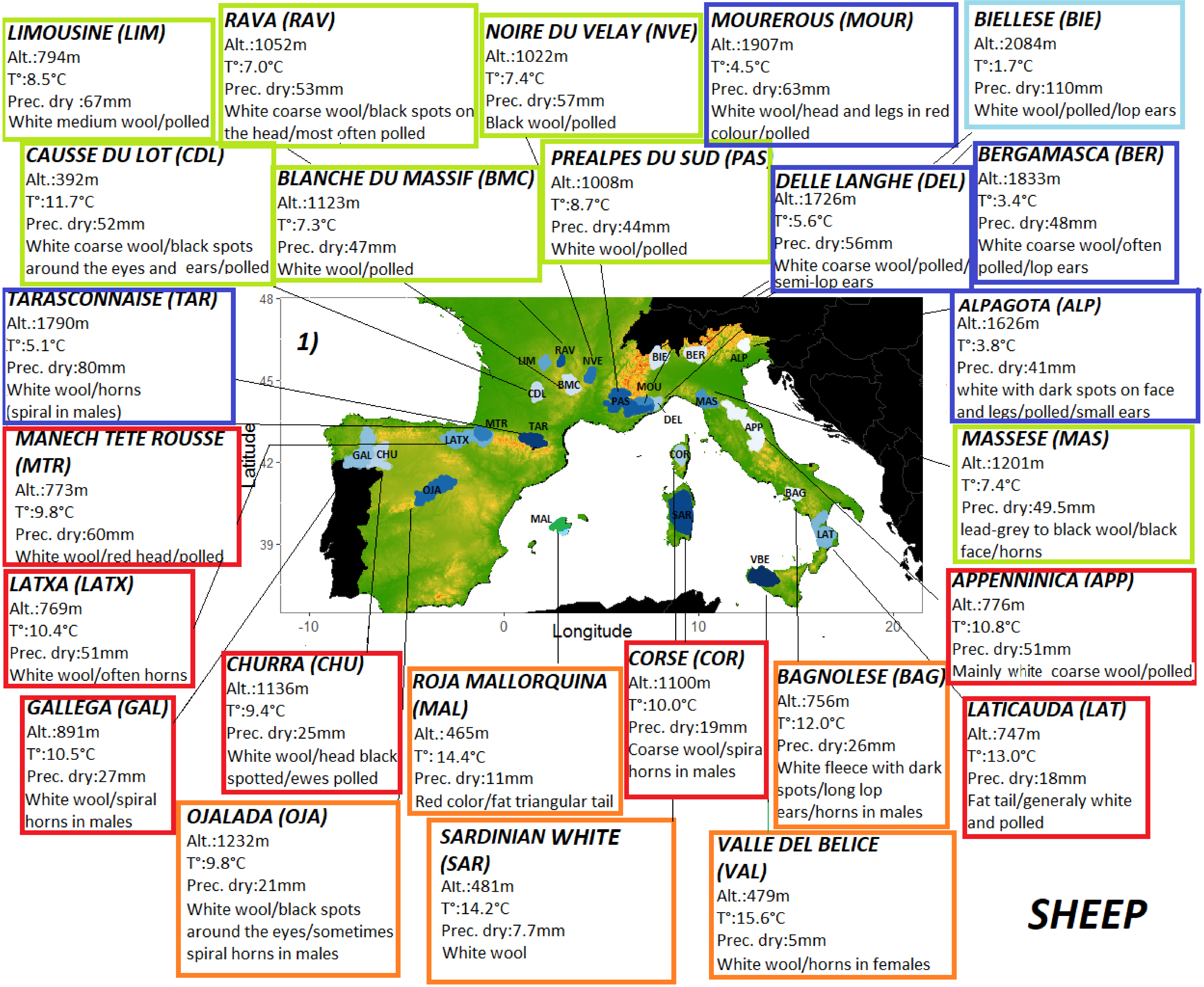

LIMOUSINE (LIM)
Alt.:794m
T°:8.5°C
Prec. dry :67mm
White medium wool/polled
RAVA (RAV)
Alt.:1052m
T°:7.0°C
Prec. dry:53mm
White coarse wool/black spots on the head/most often polled
NOIRE DU VELAY (NVE)
Alt.:1022m
T°:7.4°C
Prec. dry:57mm
Black wool/polled
MOUREROUS (MOUR)
Alt.:1907m
T°:4.5°C
Prec. dry:63mm
White wool/head and legs in red colour/polled
BIELLESE (BIE)
Alt.:2084m
T°:1.7°C
Prec. dry:110mm
White wool/polled/lop ears
CAUSSE DU LOT (CDL)
Alt.:392m
T°:11.7°C
Prec. dry:52mm
White coarse wool/black spots around the eyes and ears/polled
BLANCHE DU MASSIF (BMC)
Alt.:1123m
T°:7.3°C
Prec. dry:47mm
White wool/polled
PREALPES DU SUD (PAS)
Alt.:1008m
T°:8.7°C
Prec. dry:44mm
White wool/polled
DELLE LANGHE (DEL)
Alt.:1726m
T°:5.6°C
Prec. dry:56mm
White coarse wool/polled/semi-lop ears
BERGAMASCA (BER)
Alt.:1833m
T°:3.4°C
Prec. dry:48mm
White coarse wool/often polled/lop ears
TARASCONNAISE (TAR)
Alt.:1790m
T°:5.1°C
Prec. dry:80mm
White wool/horns (spiral in males)
ALPAGOTA (ALP)
Alt.:1626m
T°:3.8°C
Prec. dry:41mm
white with dark spots on face and legs/polled/small ears
MANECH TETE ROUSSE (MTR)
Alt.:773m
T°:9.8°C
Prec. dry:60mm
White wool/red head/polled
MASSESE (MAS)
Alt.:1201m
T°:7.4°C
Prec. dry:49.5mm
lead-grey to black wool/black face/horns
LATXA (LATX)
Alt.:769m
T°:10.4°C
Prec. dry:51mm
White wool/often horns
APPENNINICA (APP)
Alt.:776m
T°:10.8°C
Prec. dry:51mm
Mainly white coarse wool/polled
1)
48
45
42
39
-10
0
10
20
Latitude
Longitude
LIM
RAV
NVE
BIE
BER
ALP
CDL
BMC
MOU
PAS
DEL
MAS
APP
MTR
TAR
LATX
GAL
CHU
COR
OJA
SAR
MAL
BAG
LAT
VBE
GALLEGA (GAL)
Alt.:891m
T°:10.5°C
Prec. dry:27mm
White wool/spiral horns in males
CHURRA (CHU)
Alt.:1136m
T°:9.4°C
Prec. dry:25mm
White wool/head black spotted/ewes polled
ROJA MALLORQUINA (MAL)
Alt.: 465m
T°: 14.4°C
Prec. dry:11mm
Red color/fat triangular tail
CORSE (COR)
Alt.:1100m
T°:10.0°C
Prec. dry:19mm
Coarse wool/spira horns in males
BAGNOLESE (BAG)
Alt.:756m
T°:12.0°C
Prec. dry:26mm
White fleece with dark spots/long lop ears/horns in males
LATICAUDA (LAT)
Alt.:747m
T°:13.0°C
Prec. dry:18mm
Fat tail/generaly white and polled
OJALADA (OJA)
Alt.:1232m
T°:9.8°C
Prec. dry:21mm
White wool/black spots around the eyes/sometimes spiral horns in males
SARDINIAN WHITE (SAR)
Alt.:481m
T°:14.2°C
Prec. dry:7.7mm
White wool
VALLE DEL BELICE (VAL)
Alt.:479m
T°:15.6°C
Prec. dry:5mm
White wool/horns in females
SHEEP

| Goat Breeds (code and number of individuals) | country | Origin/History | Cradle of the breed/transhumant area if practiced in the past and/or today | Description/adaptation | Production/ use | Status (according to FAO 2007 if not mentioned) |
|---|---|---|---|---|---|---|
| Bionda dell Adamello (BIO, n=24) | Italy | The breed seems to be ancient. A painting from around 1760 by the Milanese painter Francesco Londonio shows a goat of the Bionda dell'Adamello type (Bigi & Zanon, 2008). | The Val Camonica in the province of Brescia, in Lombardy in northern Italy. It takes its name from the massif of the Adamello, part of the Adamello-Presanella subsection of the Rhaetian Alps.<br><br>Transhumance in Alpine pasture (Caballero et al., 2009). | Medium to large size<br>long hair ranging from light brown to blonde<br>two white streaks on the muzzle<br><br>Adaptation to pasture in the mountains | Milk also meat<br>Extensive management | Not at risk |
| Valdostana (also called Chamoisée Valdôtaine) (VAL, n=24) | | Publications dating back to 1917 mention its existence, and suggest an older origin. It would have been the result of hybridation between wild and domesticated animals (http://www.agraria.com). | Aosta Valley in northwestern Italy, particularly in the lower Ayas and Lys valleys.<br><br>Transhumance in Alpine pasture, (Caballero et al., 2009). | Medium-large-sized animals<br>Short dark blond hair<br>large horns (selection for the traditional "Batailles de Chèvres", Talenti et al., 2017)<br><br>Adaptation to pasture in the mountains | Milk/meat<br>Extensive management | Critical |
| Orobica (ORO, n=24) | | Ancient breed: The Milanese painter (17th century) Lonati Verri represents goats of the Orobica type. A Milanese lithograph from the early 19th century depicts a goatherd with some of his animals near the Eastern Porta (now Porta Venezia). Official documents mention the existence of the breed as early as 1829 (Corti et al., 1997; Corti, 2006). | The Val Gerola in the province of Sondrio, in the Bergamo Alps of northern Italy.<br><br>Transhumance in Alpine pasture (Caballero et al., 2009). | Medium size<br>long horns with flat section in males and erect ears<br>fine long hair, with a colour varying from uniform ash-grey to violet-beige<br><br>Adaptation to pasture in the mountains | Milk<br>Extensive management | Endangered |

| | | | | | | |
|---|---|---|---|---|---|---|
| Val Passiria<br>(also called Passiria/ Camosciata delle Alpi/Passeirer Gebirgsziege/Chèvre Chamoisée)<br>(VSS, n=24) | | Breed that seems to have originated from Switzerland (Porter, 2002). | Adapted to the valleys bordering Austria (Val Passiria, Upper Isarco, Sarntal and Val Senales)<br><br>Transhumance in Alpine pasture (Caballero et al., 2009). | Medium size<br>Medium to long hair<br>dark coat (from black to grey)<br><br>Adaptation to pasture in the mountains/ good adaptation to the most extreme areas | Meat<br>Extensive management | Unkown |
| Di Teramo<br>(DIT, n=24) | | Indigenous breed of the province of Teramo, unkown origin but probable crossbreeds with Garganica goat (Porter, 2002; http://www.agraria.com). | Province of Teramo, Abruzzi Region.<br><br>Transhumance in the mountain pastures of Abruzzo (Caballero et al., 2009). | Medium size<br>horns in both sexes<br>Long dark hair (black, brown, grey) | Milk | Critical |
| Ciociara Grigia<br>(CCG, n=19) | | Indigenous breed of Lazio in central Italy, unkown origin but probable crossbreeds with Garganica goat (Porter, 2002). | It is thought to have originated in the area of the Monti Aurunci and the Monti Ausoni. It takes its name from the Ciociaria, the area around Frosinone.<br><br>Transhumance in the mountain pastures of Abruzzo and Campania plains (Caballero et al., 2009). | Medium sized, dark silvery grey and light coat | Milk/meat | Endangered |
| Garganica<br>(GAR, n=20) | | Indigenous to the Gargano, it derives from cross-breeding of local animals with goats imported from western Europe (Porter, 2002; http://www.agraria.com). | Gargano promontory in the Puglia region of southern Italy.<br><br>Transhumance in Puglia. | Medium size<br>black, with long hair<br>Horns are quite big<br><br>Hardy and gregarious animal, adapted to very difficult habitat (Bigi & Zanon, 2008) | Milk/meat | Endangered |
| Argentata Dell'Etna<br>(ARG, n=25) | | Indigenous breed of Mount Etna (Porter, 2002). | The area of Mount Etna in the province of Catania and the | Medium size<br>silvery grey coat, medium-long hair | Raised primarily for milk | Not at risk |

| | | | | | | |
|---|---|---|---|---|---|---|
| | | | Monti Peloritani in the province of Messina, in Sicily. | horns of the females are upright<br><br>Adaptation to scarce pastures of Sicily (Rubino, 1993) | production, and also for meat | |
| Nicastrese<br>(NIC, n=25) | | Byzantine origin, photographic evidence suggests that it may be closely connected to the old "Araba" breed of the area (Bigi & Zannon, 2008). | The province of Catanzaro in southern Italy. | Small size<br>Black coat (keshmir undercoat), long hair | Milk/meat | Critical |
| Girgentana<br>(GGT, n=30) | | Would have arrived in Sicily with the Greeks in the 8th century (http://www.agraria.com; Porter, 2002). | The Agrigento coast in Sicily. | Medium size<br>Spiral horns, long and white hair<br><br>Hardy breed | Milk<br>High prolificity | Threatened by extinction, 400 heads recorded in 1950 (Bigi & Zannon, 2008) |
| Corse<br>(CRS, n=29) | France | The presence on the island has been documented for several millennia (https://www.racesdefrance.fr). | Corsican massifs, in particular the valleys of Caccia and Niolo.<br><br>Double transhumance between plains in winter and mountains in summer (Ravis-Giordani, 2001). | Small size<br>curved parallel horns<br>long hair, heterogeneous coat color<br><br>Hardy breed/ adaptation to scarce and rugged pastures | Milk/meat<br>Extensive management | Not at risk |
| Provençale<br>(PVC, n=18) | | Indigenous breed of Provence registered since the 19th century (https://www.racesdefrance.fr; Babo, 2000). | The cradle of the breed is located in the "Provence of hills" (Alpes de Haute Provence).<br><br>This breed was not transhumant. | Large size<br>large long hanging ears<br>medium long hair, heterogeneous coat color<br><br>Adaptation to scarce and dry pastures | Milk | Endangered<br>This breed, which almost disappeared, was taken over in 1993 by a breeders' association (http://www.capgenes.com) |
| Pyrénéenne<br>(PYR, n=25) | | Very old origin in the Pyrenees (Fournier, 2006). | Pyrenees - Atlantic to Ariège. | Medium size<br>Long hair, dark coat | Milk/meat | Endangered |

| | | | | | | |
|---|---|---|---|---|---|---|
| | | | Transhumance in Pyrenean pastures (Fanica, 2008) some herds moved with the Pyrenean sheep across the Landes to Gironde and Dordogne. | | Extensive management | It almost disappeared in the second half of the 20th century (https://www.chevredespyrenees.org) |
| Bermeya (BEY, n=24) | Spain | Indigenous breed of the North of Spain, probably descending from Capra aegagrus (https://www.mapa.gob.es). | The southern part of the Principality of Asturias, in northern Spain.<br><br>Short transhumance (transtermitance). | Medium size<br>Short hair, red coat<br><br>suitable for mountainous areas and humidity | Mainly meat and also milk | Endangered |
| Blanqua De Rasquera (RAS, n=20) | | Municipal Ordinances of Vila de Rasquera (Tarragona) refers to the goat population in 1573 (Sabaté et al., 2011). | Several regions in southern Catalonia (Baix Ebre, Ribera d'Ebre and Terra Alta), its name comes from the municipality of Rasquera, in the Ebro.<br><br>Horizontal transhumance to the Pyrenean pastures (http://ipcite.cat). | Medium size<br>short hair, white coat with sometimes wide black spots large, drooping ears<br><br>Hardy breed | Meat and also milk<br>Extensive management | Endangered |
| Malaguena (MLG, n=41) | | Certainly very old origin of the breed; there are traces of its existence that date back to the 4th millennium BC (Garcia-Dory et al., 1990). | Malaga in the south of the Iberian Peninsula. | Medium size<br>Short hair, sandy to red coat, horns sometimes twisted | Milk/meat | Not at risk |
| Mallorquina (MAL, n=20) | | Ancient breed that was introduced to the island around 2500/1400 BC (Segui et al., 2005). | Majorca, mainly in the mountains: Sierra de tramuntana and Sierra de Llevant. | Medium size<br>Short hair, red or red and black coat<br><br>Adaptation to scarce, dry and rugged areas | Meat<br>Extensive management | Endangered |

| **Sheep Breeds (code)** | **country** | **Origin/History** | **Cradle of the breed/transhumant area if practiced in the past and/or today** | **description** | | **status** |
|---|---|---|---|---|---|---|
| Manech Tête Rousse (MTR, n=25) | France | Very ancient Basque origin (https://www.races-montagnes.com; Babo, 2000). | Basque hillsides (Basse Navarre, Basse Soule).<br><br>Transhumance in Pyrenean pasture. | Medium size<br>White wool, red head, long lopping ears<br><br>Ability to breed out of season<br>Hardy breed, good walker | Milk | Not at risk |
| Tarasconnaise (TAR, n=15) | | Very old breed derived from an original population of the Pyrenees (https://www.races-montagnes.com). | Central Pyrenees and particularly in the Ariège, Hautes-Pyrénées and Haute-Garonne.<br><br>Transhumance in Pyrenean pasture. | Medium size<br>White wool, spiral horns in males<br><br>Hardy breed, good walker, ability to breed out of season | Meat | Not at risk |
| Causse Du Lot (CDL, n=20) | | It is believed to come from a very old sheep strain that lived on the northern slope of the Garonne. Its origin goes back at least to the Gallo-Roman period (https://www.races-montagnes.com; Clozier, 1932). | Limestone plateaus of the Lot (the Causses of Quercy)<br><br>Horizontal transhumance linking the Lot to the Auvergne; winter transhumance was used during periods of extreme cold (Leveau, 2016; Clozier, 1932; Delhoume, 2005). | Medium to large size<br>White coarse wool, black spots around the eyes and ears<br><br>Hardy breed, it would be resistant to piroplasmosis; ability to breed out of season | Meat/milk<br>Extensive management | Not at risk |
| Limousine (LIM, n=18) | | The breed would have emerged at the end of the 19th century as a result of crossbreeding between different breeds of the Massif Central (https://www.races-montagnes.com). | Millevaches plateau, in the mid-mountain area. | Medium size<br>White medium wool<br><br>Adapted to a difficult territory with acid soil and harsh climate, ability to breed out of season | Meat | Not at risk |

| | | | | | | |
|---|---|---|---|---|---|---|
| Rava<br>(RAV, n=20) | | Breed of the volcanic plateaus of Auvergne; it was first described in 1826 (https://www.races-montagnes.com). | The Chaîne des Puys in the Massif Central.<br><br>Transhumance in the Massif Central. | Medium size<br>White coarse wool, black spots on the head<br><br>Adapted to a difficult territory with acid soil and harsh climate, ability to breed out of season | Meat | Not at risk |
| Blanche Du Massif<br>(BMC, n=20) | | Old breed that belongs to the Caussenard group (https://www.races-montagnes.com). | The Margeride in the South part of the Massif Central.<br><br>Horizontal transhumance linking the Langedoc to the Auvergne (Leveau, 2016). | Medium size<br>White wool<br><br>Adapted to a difficult territory, ability to breed out of season | Meat | Not at risk |
| Noire Du Velay<br>(NVE, n=19) | | Very old breed of Auvergne; it seems to have arrived in France around 1500 BC when the Celtic peoples settled in the Massif Central; writings mention its existence as early as the 17th century (https://www.races-montagnes.com; Brunelin, 2012). | Velay volcanic plateau, in the Auvergne region.<br><br>Horizontal transhumance linking the Languedoc to the Auvergne (Leveau, 2016). | Medium size<br>black skin and wool of black-brownish colour<br><br>Good walker, adapted to a difficult territory with acid soil and harsh climate, ability to breed out of season | Meat | Not at risk<br>It almost disappeared at the beginning of the 20th century with the arrival of better conformed breeds |
| Préalpes Du Sud<br>(PAS, n=17) | | It would be the result of the old mixing of the Sahune, Quint and Savournon breeds (https://www.races-montagnes.com; Duclos & Mallen, 1998). | The Southern Alps at the border between the Dauphiné and Provence.<br><br>Transhumance in Alpine pastures. | Medium size<br>White wool<br><br>Adaptation to dry mountain, ability to breed out of season | Meat | Not at risk |
| Mourerous, also called "La Rouge", "Péone" ou "Rouge de Guillaumes" | | It could have been crossbred with breeds imported from North Africa (https://www.races-montagnes.com). | Péone in the department of Alpes Maritimes.<br><br>Transhumance in Alpine pastures. | Medium size<br>White wool, red head and legs<br><br>Cold and drought resistance, ability to walk in the | Meat | Not at risk<br>For a long time known as a "threatened" or "small breed" |

| | | | | | | |
|---|---|---|---|---|---|---|
| (MOUR, n=16) | | | | mountain, ability to breed out of season | | |
| Corse<br>(COR, n=16) | | Ancestral breed of Corse; It would originate from a Pyrenean breed imported on the island several centuries ago (https://www.races-montagnes.com; Babo, 2000). | Mountainous areas of Corse.<br><br>Double transhumance between plains in winter and mountains in summer (Ravis-Giordani, 2001). | Medium size<br>Coarse wool, spiral horns in males<br><br>Adaptation to mountain pastures, ability to breed out of season | Milk | Not at risk |
| Delle Langhe<br>(DEL, n=24) | Italy | Indigenous breed of the Piedmont (Porter, 2002). | The mountainous area of the Alta Langa in Piedmont (province of Cuneo), in northwestern Italy.<br><br>Transhumance in Alpine pasture (Caballero et al., 2009). | Medium to large size<br>White coarse wool, polled, semi lop hears<br><br>Adaptation to mountain pastures | Milk/meat | Endangered |
| Biellese<br>(BIE, n=21) | | Indigenous breed of the province of Biella in Piedmont (Porter, 2002). | The province of Biella, in Piedmont in north-western Italy.<br><br>Transhumance in Alpine pasture, (Caballero et al., 2009, Battaglini 2014)<br>still practice vertical trasterminance (3/4 of total) between the mountains and the plain areas of Lombardy or Emilia Romagna (Corti, 2007). | Medium to large size<br>White coarse wool, polled, lop hears<br><br>Adaptation to mountain pastures | Meat and also wool in the past | Not at risk |
| Bergamasca<br>(BER, n=24) | | Very ancient breed, we can trace its existence back to the 13$^{th}$ century (Astori, 1963). | The mountainous part of the province of Bergamo, in Lombardy in northern Italy. | Medium to large size<br>White coarse wool, polled, lop hears | Meat | Not at risk |

| | | | | | | |
|---|---|---|---|---|---|---|
| | | | Transhumance in Alpine pasture, (Caballero et al. 2009)<br>still practice vertical trasterminance (3/4 of total) between the mountains and the plain areas of Lombardy or Emilia Romagna (Corti, 2007). | Adaptation to mountain pastures | | |
| Alpagota<br>(ALP, n=24) | | Ancient breed, the sheep's emblem is found on the coat of arms of Chies d'Alpago (15th century) (Pastore, 2005). | The historic region of the Alpago in northern Italy<br><br>Transhumance in Alpine pasture (Caballero et al., 2009). | Small to medium-size<br>white with dark spots, polled small ears<br><br>Adaptation to hills and mountains | Meat/milk | Not at risk |
| Massese<br>(MAS, n=24) | | Indigenous breed that belongs to the Apennine group, with an ancient origin: Machiavelli refers to the breed in his writings (early 16th century). | The Alpi Apuane mountains of the province of Massa Carrara, in Tuscany, central Italy.<br><br>Transhumance in the mountain pastures of Appennins (Caballero et al., 2009). | Small to medium size<br>lead-grey to black<br><br>Adaptation to mountain pastures | Milk/meat | Not at risk |
| Appenninica<br>(APP, n=24) | | The origin of the breed can be traced back to the end of the 19th century, when sheep of the Bergamasque breed were imported into Tuscany, Umbria, Romagna, Marche and Abruzzo. These subjects were crossed with the pre-existing Apennine populations in order to intensify meat production (http://biodiversita.umbria.parco3a.org). | The central Apennine mountains of Italy.<br><br>Transhumance in the mountain pastures of Appennins and Abruzzo (Caballero et al., 2009). | Medium size<br>White, coarse wool, polled | Meat and also wool | Not at risk |

| | | | | | | |
|---|---|---|---|---|---|---|
| Laticauda<br>(LAT, n=24) | | It should result from ancient hybridisation of local breeds with Barbary (or Barbarin) sheep of Maghrebi origin. The Bourbon king Charles VII of Naples could have brought it in the area (Bigi & Zannon 2008). | Campania and Calabria, in southern Italy, in particular the provinces of Avellino, Benevento and Caserta.<br><br>This breed was not transhumant. | Medium to large size<br>Generally white, fat tail, polled | Milk/meat | Unkonwn |
| Bagnolese<br>(BAG, n=23) | | A native campanian breed, that probably derives from the crossbreeding of the Barbaresca breed and the local breeds of the Appennines (http://www.agraria.com). | The area surrounding Bagnoli Irpino in the province of Avellino, in Campania in southern Italy.<br><br>Transhumance to the mountain pastures in summer. | Medium size<br>white fleece with dark spots, coarse wool, long hanging ears<br><br>Well adapted to pasture in rough conditions | Milk/meat | Endangered-maintained |
| Valle Del Bellice<br>(VAL, n=24) | | It appears to result from the three-way hybridization of the Sicilian Pinzirita and Comisana breeds with Sarda stock brought from Sardinia (Porter, 2002). | The valley of the Belice river in south-western Sicily. | Medium size<br>White coat (Sarda), black coat (Sardinian black), long coarse wool | Milk | Not at risk |
| Sardinian White<br>(SAR, n=24) | | Indigenous to the island of Sardinia with very ancient origin (http://www.agraria.com). | Mountains and plains of Sardinia.<br><br>Transhumance to mountain pastures. | Medium size<br>White coat long coarse wool | Milk | Not at risk |
| Latxa<br>(LATX, n=24) | Spain | Basque breed called Manech in France, and one of the oldest breed in Spain. There are engravings dating back to 1872 that represent the breed (https://www.mapa.gob.es, Porter 2002; Garcia-Dory, 1990). | the north-west of Navarre and in the east and south of the province of Guipúzcoa, in the north of Spain<br><br>Transhumance in Pyrenean pasture, (Caballero et al., 2009) still practice vertical trasterminance (3/4 of total) between the mountains and the plain areas. | Medium size<br>White long wool, the colour of the head depends on the variety, twisted horns in males<br><br>Adapted to rugged terrain and humidity | Milk and also meat | Not at risk |

| | | | | | | |
|---|---|---|---|---|---|---|
| Gallega<br>(GAL, n=27) | | Very ancient Galician breed, originary from Ovis aries celtibericus, it would go back 6,000 years (Garcia-Dory, 1990). | South east of Gallicia. | Small to medium size<br>White or black coat, spiral horns in males | Meat and also milk and wool | Unknown, according to FAO<br>Undangered, according to gouvernemental Decret |
| Churra<br>(CHU, n=30) | | Primitive Iberian breed, it comes from *Ovis aries celticus* and gives its name to the Churro trunk. Traces of its existence date back to pre-Roman times (Garcia-Dory, 1990). | Zamora province in Castile and León. | Medium size<br>White wool, head black spotted | Milk/meat | Not at risk |
| Ojalada<br>(OJA, n=24) | | Ancient breed that belongs to the Iberian Trunk, whose ancestral representative is the *Ovis aries Ibericus* (Garcia-Dory, 1990). | The province of Soria in central Spain. | Medium size<br>White wool, black spots around the eyes, sometimes spiral horns in males | Milk/meat/wool | Not at risk according to FAO;<br>Endangered according to the website of the Spanish Ministry of Agriculture |
| Roja Mallorquina<br>(MAL, n=28) | | Ancient breed, resulting from the crossing of two sheep trunks, one from the south of the European continent and the other from the North African countries (https://www.mapa.gob.es). | Island of Mallorca, especially the southern area. | Medium to large size<br>Red coat, fat triangular tail<br><br>Adapted to dryness | Meat/wool | Endangered |

a)

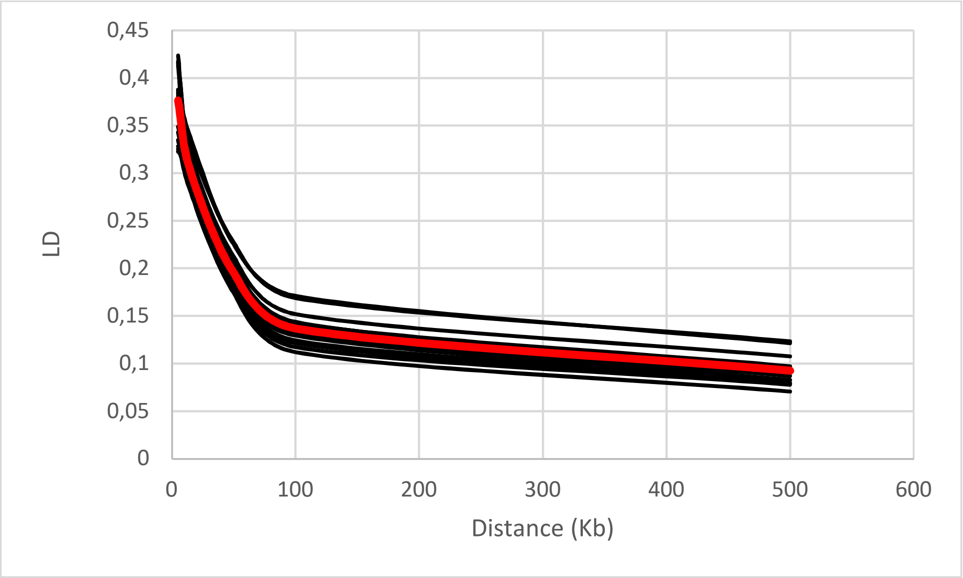

Mean LD for sheep in red: 5Kb=0.37 ; 15Kb=0.30 ; 50Kb=0.19 ; 100Kb=0.13 ; 500Kb=0.09

b)

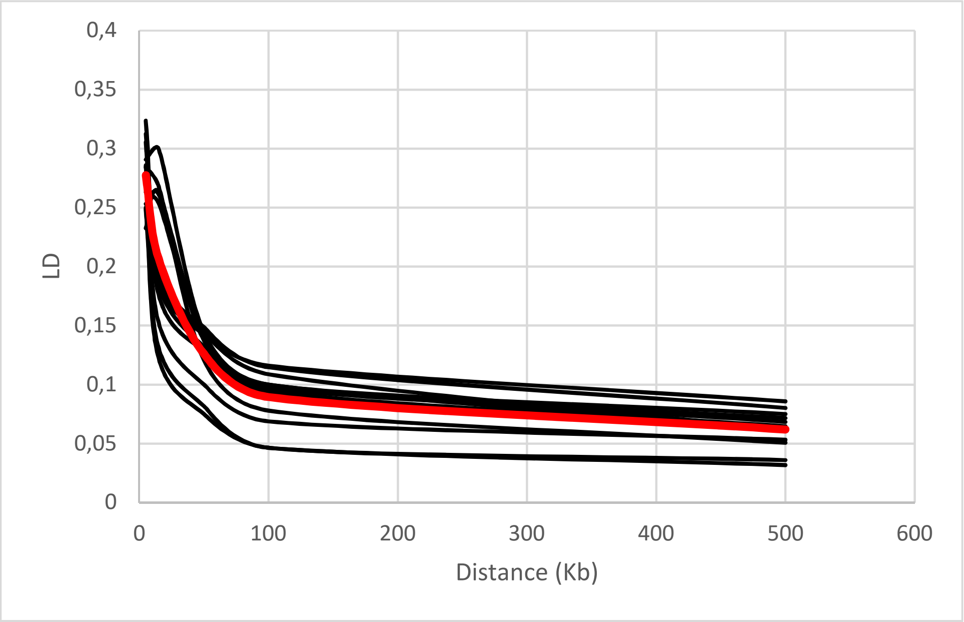

Mean LD for goat in red: 5Kb=0.28 ; 15Kb=0.21 ; 50Kb=0.13 ; 100Kb=0.09 ; 500Kb=0.06

### Genetic relationships between the Mediterranean goat breeds

For K=2 (Figure 1), the admixture analysis reveals a gradient that follows the geographical disposition of the goat breeds along the Mediterranean arc. Spanish breeds show a predominance of orange; the proportion of orange gradually decreases to give place to a dominant blue within the breeds of southern Italy. At K=3, the Italian DIT breed is individualized from GGT, then at K=4 it is the alpine ORO breed that stands out. At K=6, the proximity of the Spanish breeds is obvious, with a clear link between BEY and PYR, the breeds close to the Pyrenees, respectively Spanish and French. In the Italian group, the VAL and GAR breeds show their originality. At K=17, all breeds are individualized, with the exception of alpine breeds, BIO and VSS, which remain very close, as well as breeds from southern Italy, CCG, NIC and ARG.

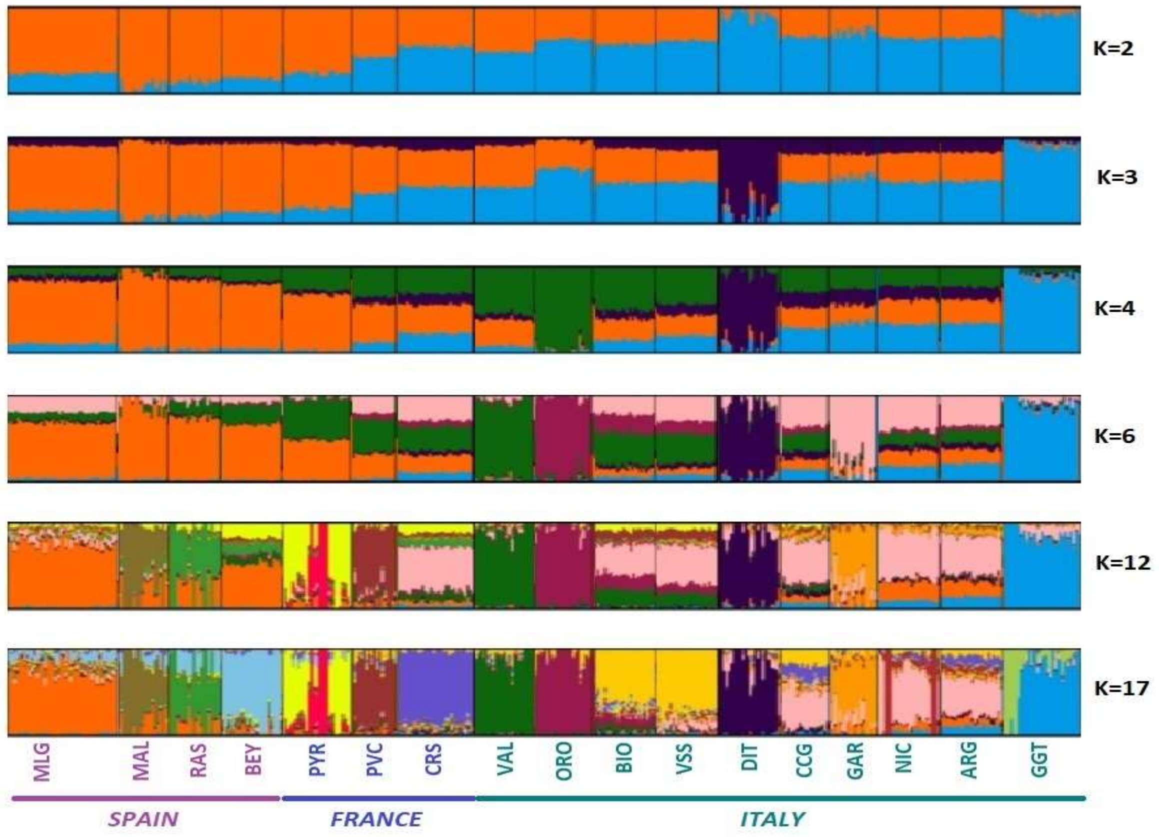

Figure 1. Bayesian clustering performed with ADMIXTURE software on Mediterranean goat breeds. K = number of clusters; see the correspondence between breed names and codes in Supplementary table 1.

### Genetic relationships between the Mediterranean sheep breeds

For K=2 (Figure 2), the major contrast is established between the Italian Alpine breeds BIE, BER, ALP and the others. Within the Spanish group, the MAL breed is very clearly individualized from K=8. The Spanish Pyrenean breed LATX shows a strong proximity with the French MTR, also Pyrenean, and this even for K=25. As for the French breeds, the first to individualize are LIM, CDL and COR. At K=25, a clear link between PAS, MOUR and NVE is maintained. The Italian breeds, DEL, then SAR, VAL and MAS, are individualized between K=4 and K=8. At K=25, the analysis shows a very strong proximity between BIE and BER, as well as a persistent link between LAT and BAG.

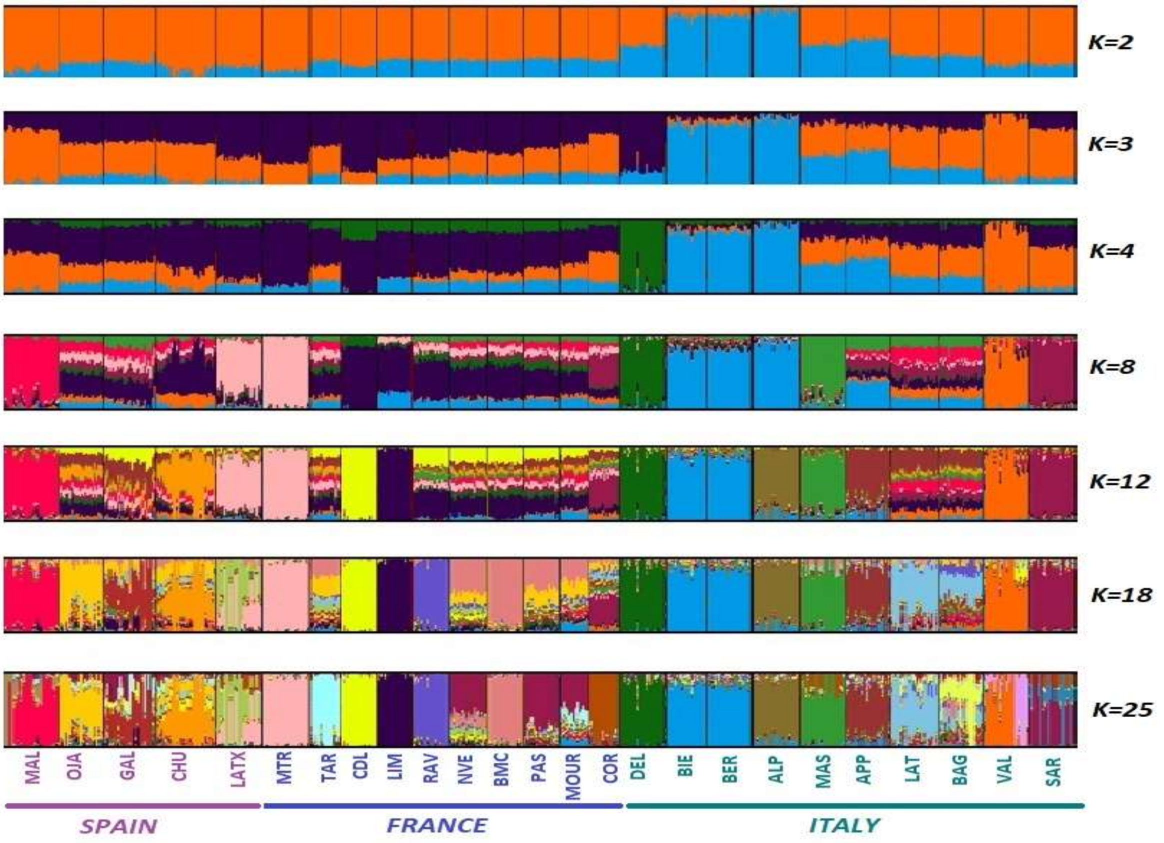

Figure 2. Bayesian clustering performed with ADMIXTURE software on Mediterranean sheep breeds. K = number of clusters; see the correspondence between breed names and codes in Supplementary table 1.

### K choice

For both goats and sheep (Figure 3), the cross-entropy curve decreases as K increases, which is characteristic of a distance isolation pattern (François, 2016). A plateau reached at K=10 for goats and K=9 for sheep, indicating that fine structuring could be apparent from this threshold. In sheep, a dropout at K=3 could be the reflection of an under-structuring by country.

The Mantel tests showed correspondence between geographic and genetic distances (p-value = 0.049 for goats and 0.001 for sheep), supporting the Isolation By Distance (IBD) pattern.

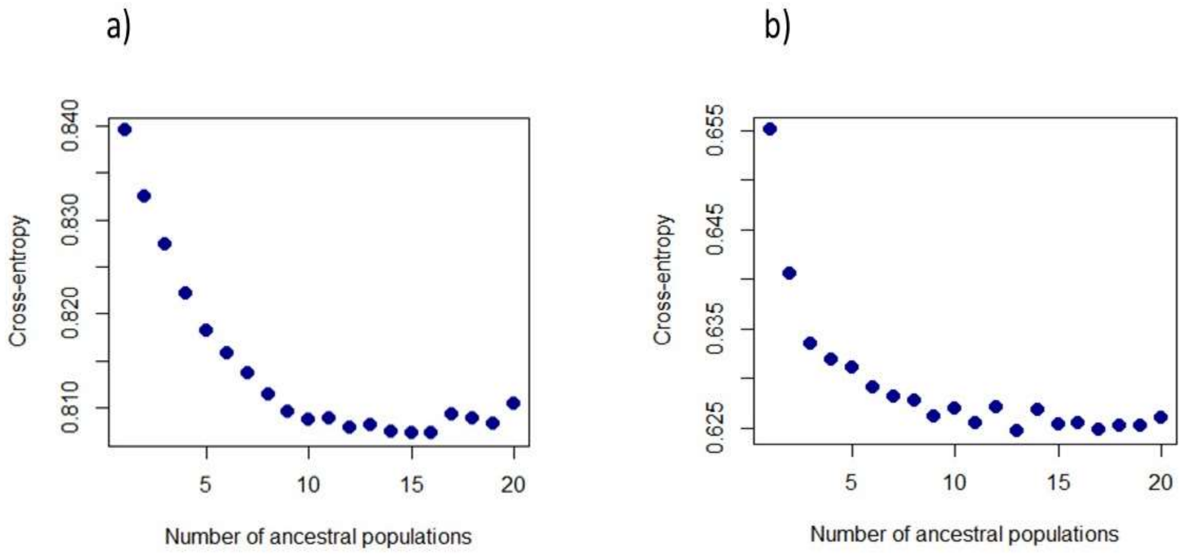

Figure 3. Cross-entropy plot for the number of cluster K = 1-20, considering: a) the goat dataset and b) the sheep dataset.

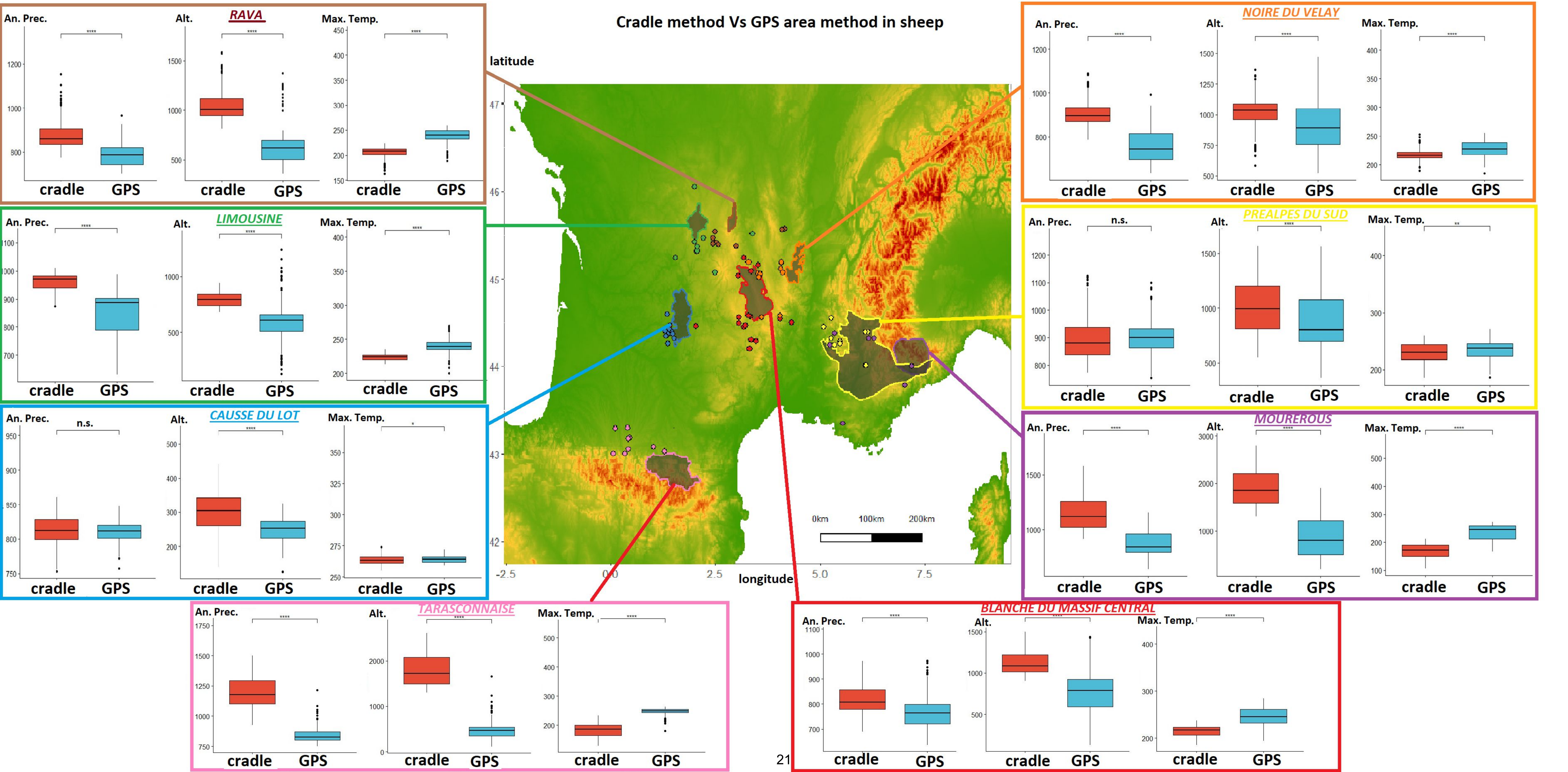
Cradle method Vs GPS area method in sheep
latitude
longitude
RAVA
NOIRE DU VELAY
LIMOUSINE
PREALPES DU SUD
CAUSSE DU LOT
MOUREROUS
TARASCONNAISE
BLANCHE DU MASSIF CENTRAL
An. Prec.
Alt.
Max. Temp.
cradle
GPS
n.s.
0km
100km
200km

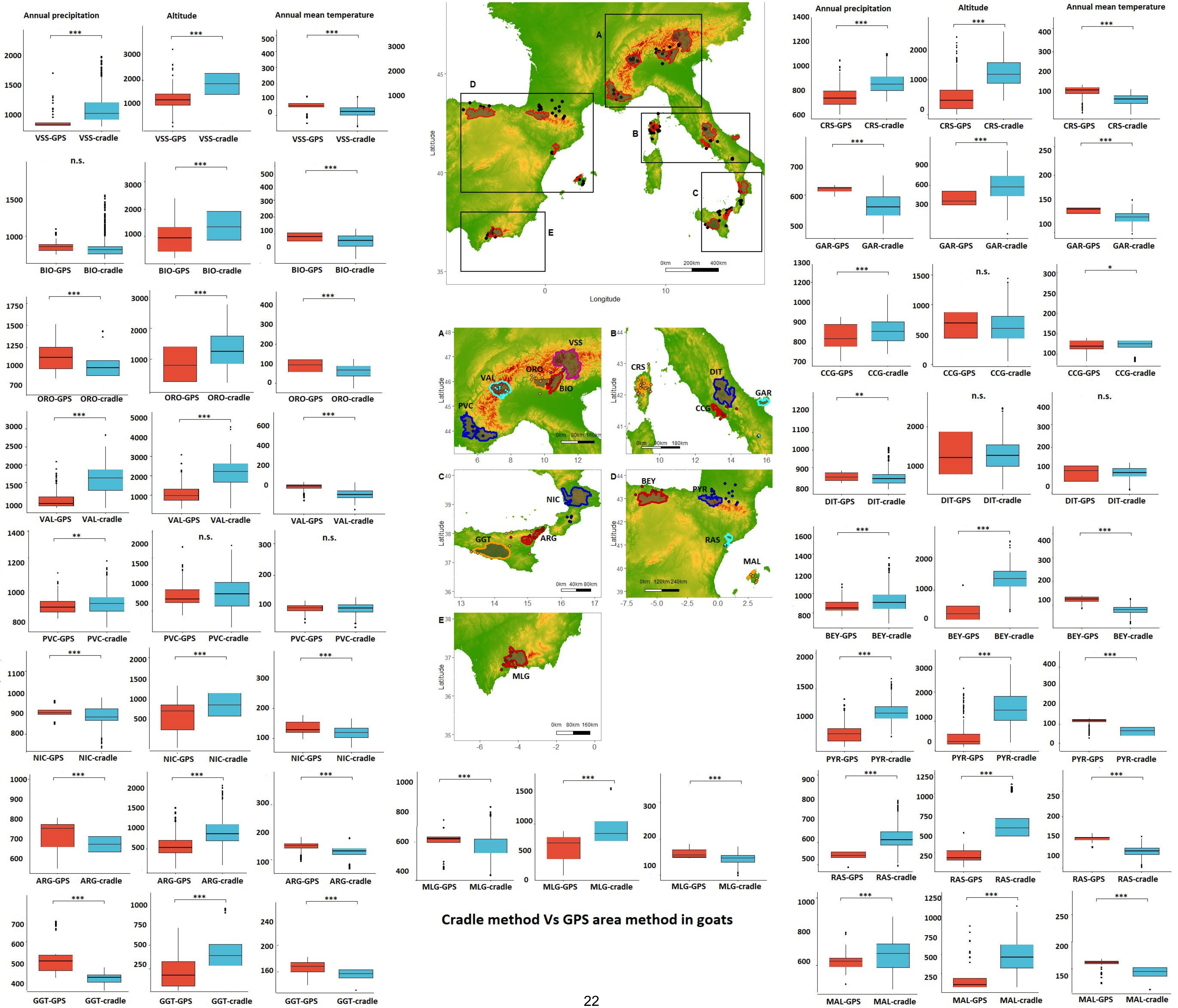

Cradle method Vs GPS area method in goats

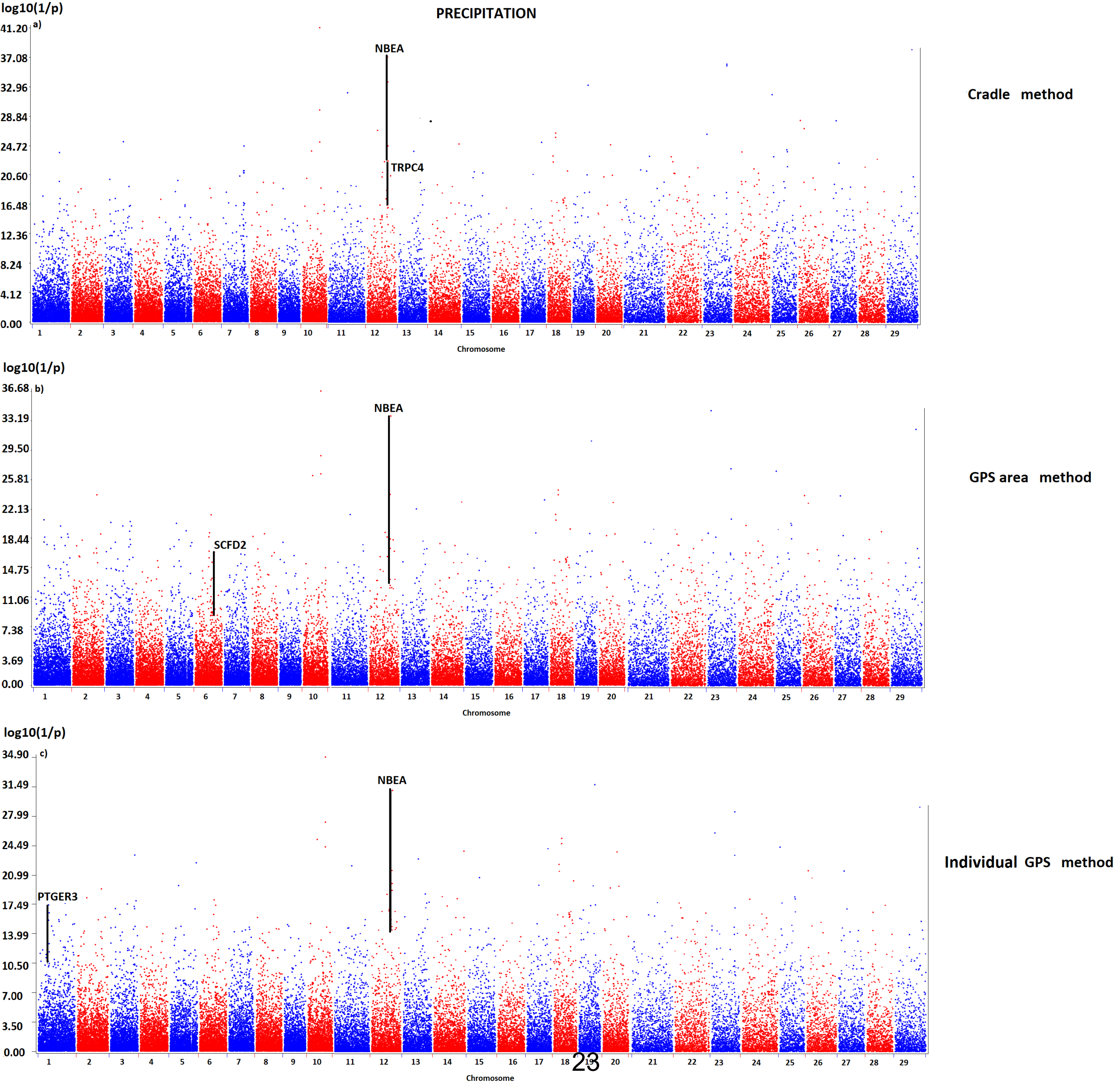
PRECIPITATION
log10(1/p)
a)
NBEA
TRPC4
Cradle method
b)
NBEA
SCFD2
GPS area method
c)
NBEA
PTGER3
Individual GPS method
Chromosome

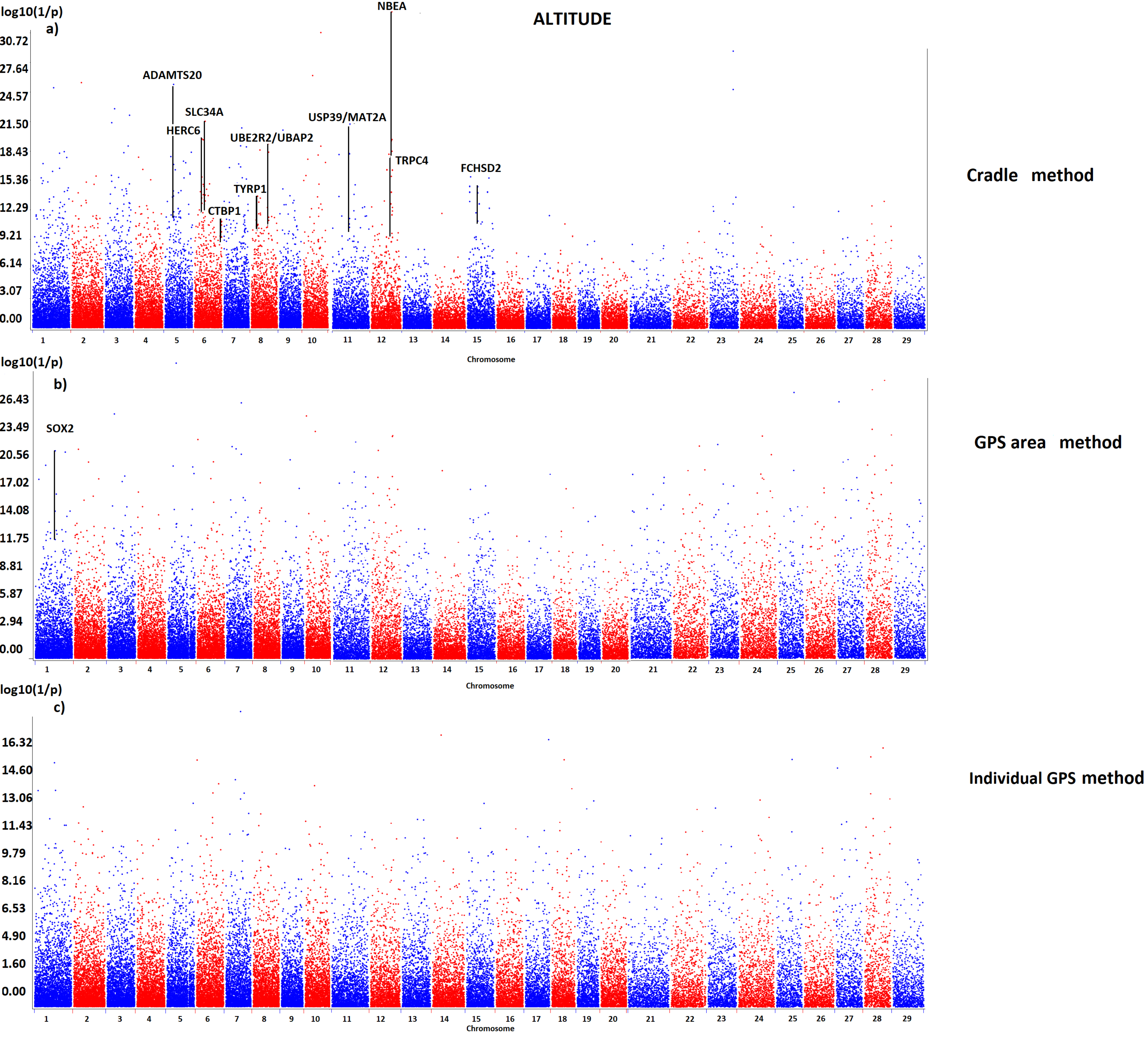

ALTITUDE
a)
log10(1/p)
NBEA
ADAMTS20
SLC34A
HERC6
USP39/MAT2A
UBE2R2/UBAP2
TRPC4
FCHSD2
TYRP1
CTBP1
Cradle method
b)
log10(1/p)
SOX2
GPS area method
c)
log10(1/p)
Individual GPS method
Chromosome

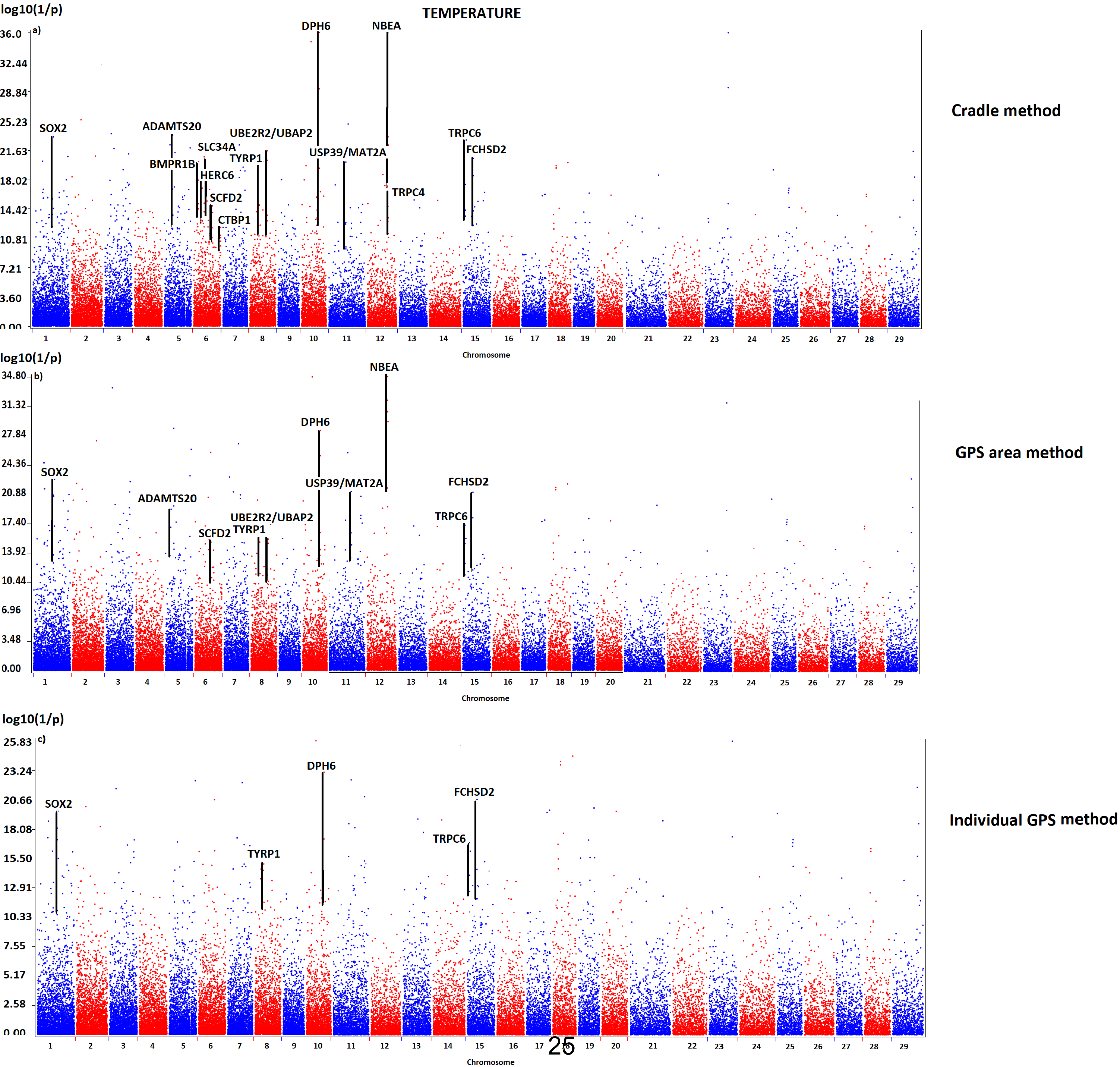
TEMPERATURE
a)
log10(1/p)
Cradle method
SOX2
ADAMTS20
BMPR1B
SLC34A
HERC6
SCFD2
CTBP1
UBE2R2/UBAP2
TYRP1
DPH6
USP39/MAT2A
NBEA
TRPC4
TRPC6
FCHSD2
Chromosome
b)
log10(1/p)
GPS area method
SOX2
ADAMTS20
SCFD2
TYRP1
UBE2R2/UBAP2
DPH6
USP39/MAT2A
NBEA
TRPC6
FCHSD2
Chromosome
c)
log10(1/p)
Individual GPS method
SOX2
TYRP1
DPH6
TRPC6
FCHSD2
Chromosome

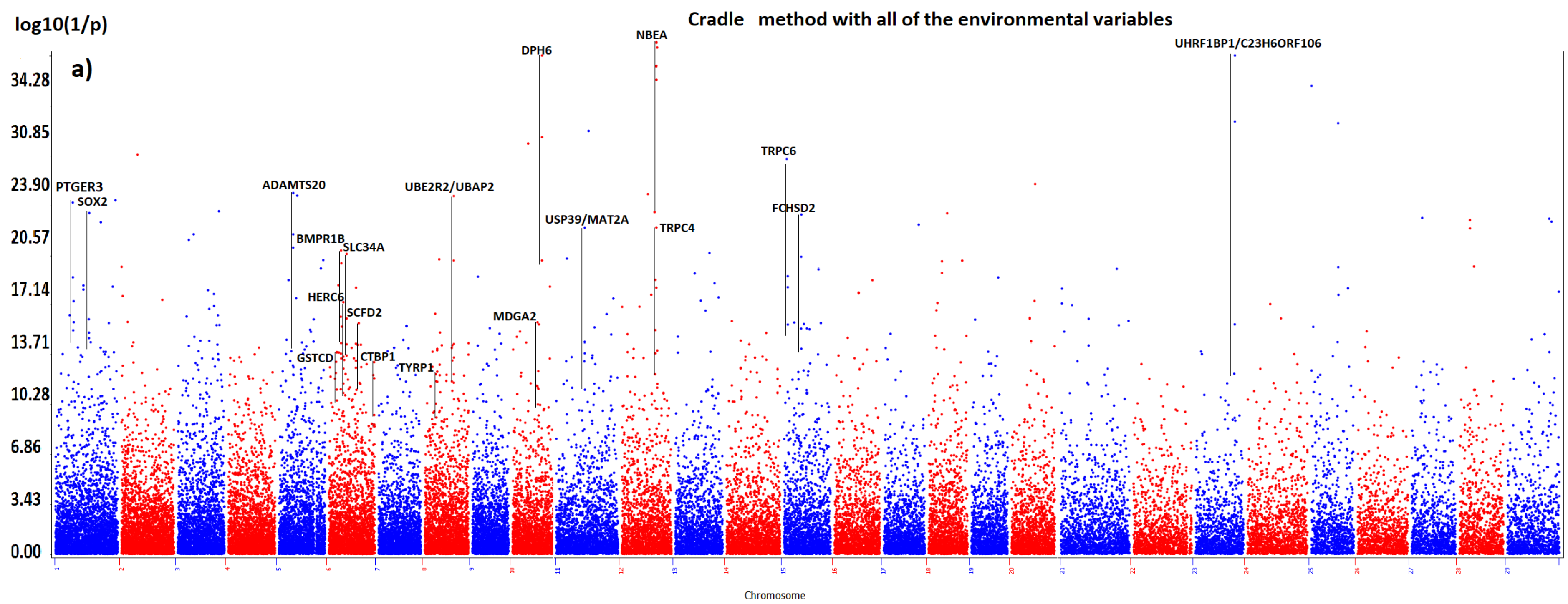
Cradle method with all of the environmental variables
log10(1/p)
a)
34.28
30.85
23.90
20.57
17.14
13.71
10.28
6.86
3.43
0.00
Chromosome
PTGER3
SOX2
ADAMTS20
BMPR1B
SLC34A
HERC6
SCFD2
GSTCD
CTBP1
TYRP1
UBE2R2/UBAP2
MDGA2
DPH6
USP39/MAT2A
NBEA
TRPC4
TRPC6
FCHSD2
UHRF1BP1/C23H6ORF106

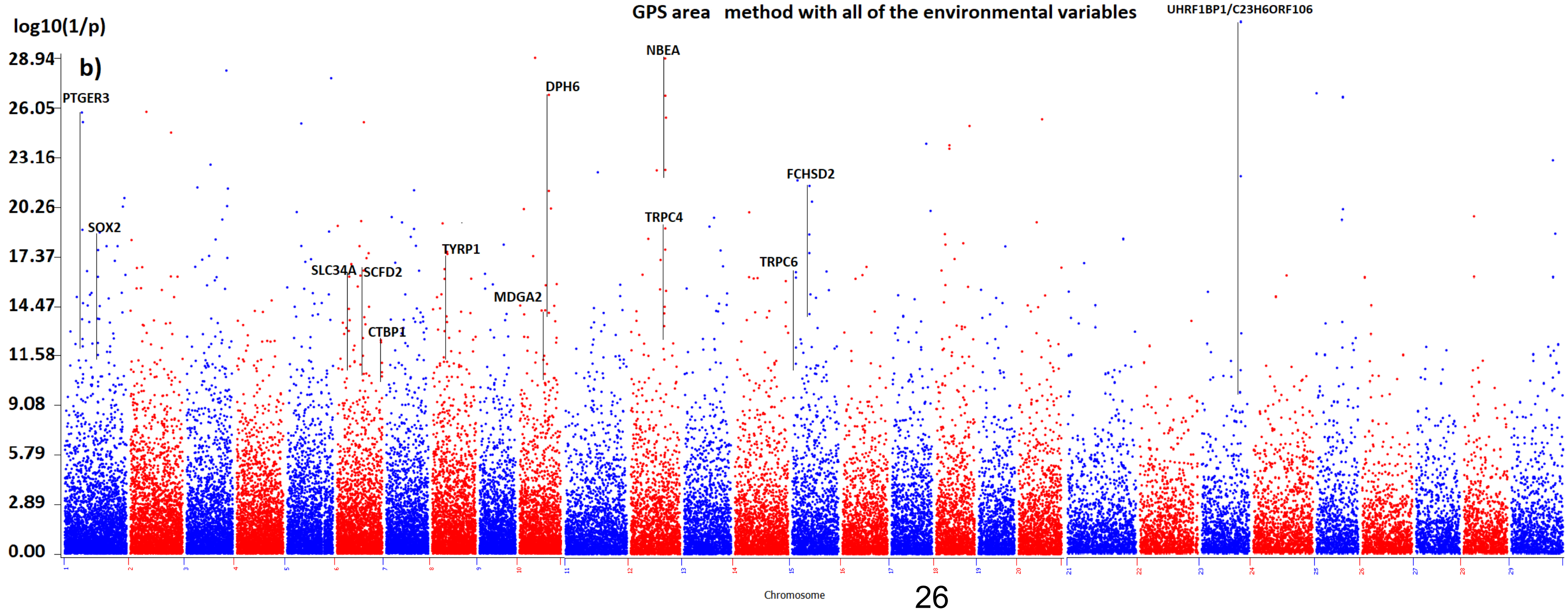
GPS area method with all of the environmental variables
log10(1/p)
b)
28.94
26.05
23.16
20.26
17.37
14.47
11.58
9.08
5.79
2.89
0.00
Chromosome
PTGER3
SOX2
SLC34A
SCFD2
CTBP1
TYRP1
MDGA2
DPH6
NBEA
TRPC4
TRPC6
FCHSD2
UHRF1BP1/C23H6ORF106

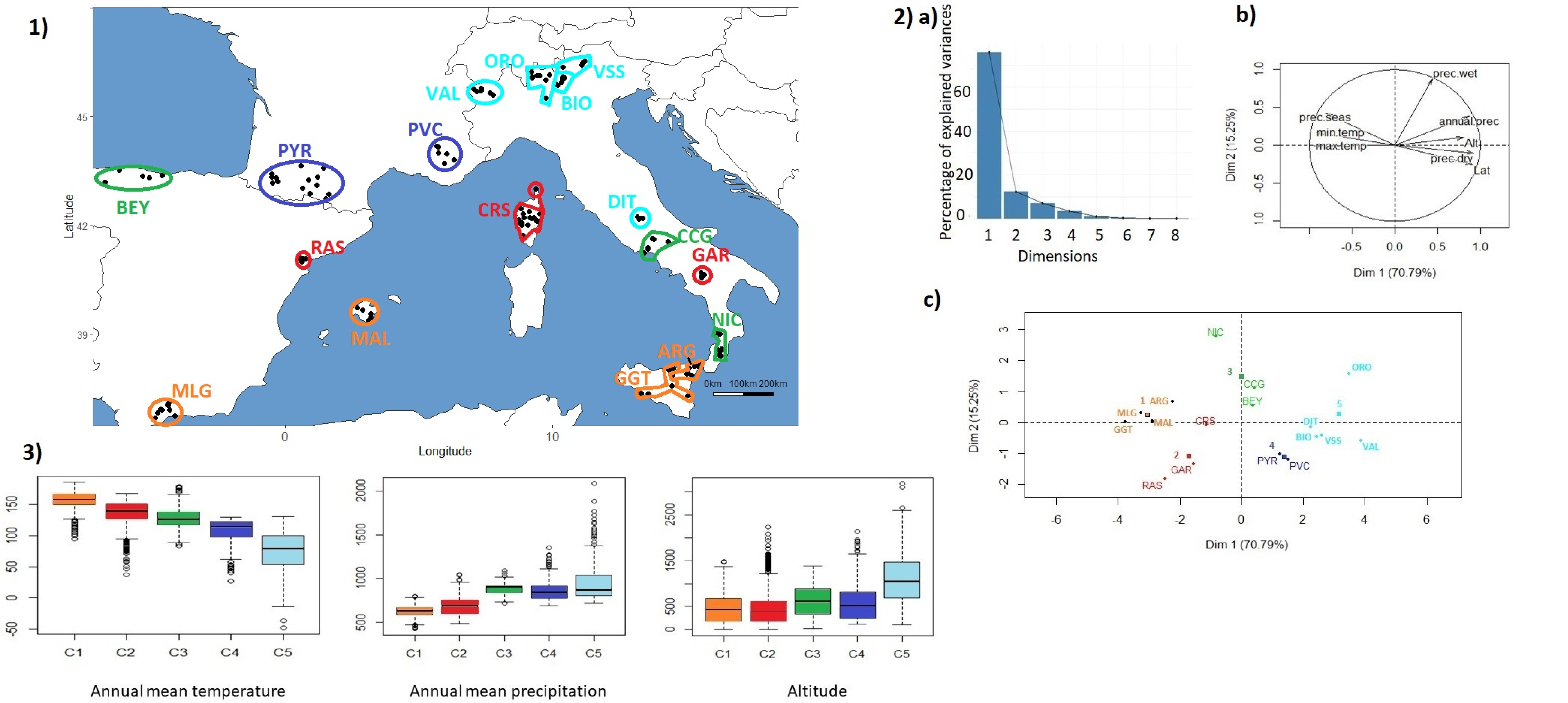

| | C1 | C2 | C3 | C4 | C5 |
|---|---|---|---|---|---|
| mean | 156.9 | 133.1 | 128.5 | 107.6 | 81.9 |
| s.d. | 14.3 | 23.1 | 18.0 | 21.2 | 36.4 |

| | C1 | C2 | C3 | C4 | C5 |
|---|---|---|---|---|---|
| mean | 632.0 | 692.1 | 884.2 | 873.6 | 949.8 |
| s.d. | 84.7 | 102.4 | 62.3 | 121.9 | 180.5 |

| | C1 | C2 | C3 | C4 | C5 |
|---|---|---|---|---|---|
| mean | 444.3 | 485.3 | 614.6 | 578.0 | 1067.5 |
| s.d. | 287.1 | 426.7 | 326.6 | 448.1 | 631.8 |

a)

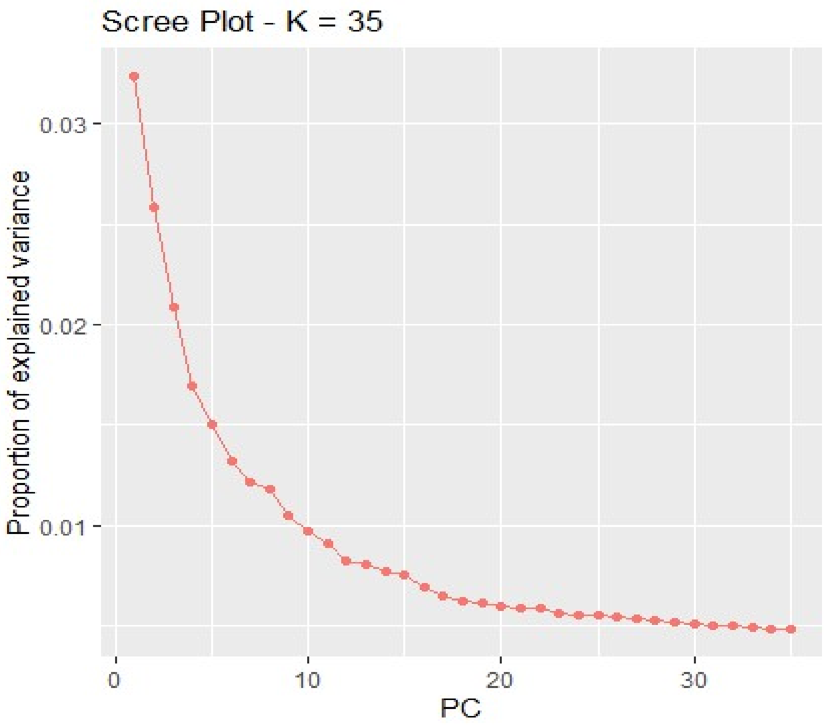
Scree Plot - K = 35
Proportion of explained variance
0.03
0.02
0.01
0
10
20
30
PC

b)

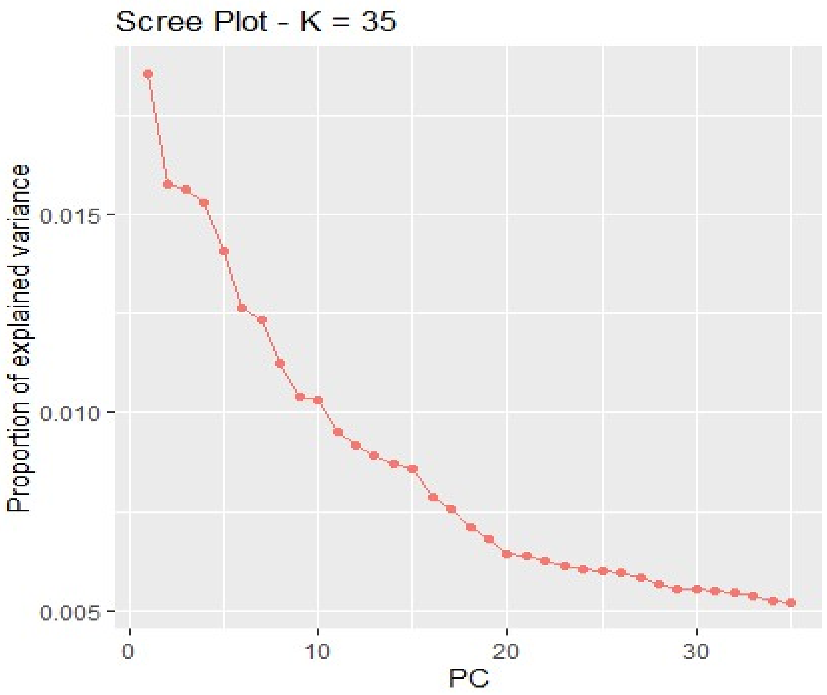
Scree Plot - K = 35
Proportion of explained variance
0.015
0.010
0.005
0
10
20
30
PC

| Chr | Numb. of signatures | Signature range | Identified by PCAdapt | Identified by LFMM | Gene(s) identified (genes around)<br>In bold gene(s) with the highest probability of being under selection | Agronomic trait/function reported in the literature for domestic mammals | Literature on genes identified in relation to environmental adaptation in mammals and birds |
|---|---|---|---|---|---|---|---|
| 1 | 2 | 46047752-46411087 | | | **PTGER3** (ZRANB2) | | |
| | | 84242981-84773966 | | | Intergenic zone between ATP11B and **SOX2** | | **SOX2**: implicated in circadian behavioral rhythms in murine (Cheng et al., 2019), cold adaptation in marmots (Bai et al., 2019) |
| 5 | 1 | 36576428-37074733 | | | **ADAMTS20** | **ADAMTS20:** Coat coloration (Oget, Servin, & Palhière 2019; Bertolini et al., 2018) | **ADAMTS20 :** associated with high altitude condition in Chinese sheep (Yang et al., 2016) |
| 6 | 7 | 19677067-20300302 | | | **GSTCD**/PPA2/**TET2** (ARHGEF38/INTS12/NPNT) | | **GSTCD**: adaptation to hypoxia in yaks (Wang et al., 2019), adaptation to climate variables in South African goats (Mdladla, 2016) |
| | | 30139296-30874402 | | | **BMPR1B/**PDLIM5/HPGDS/SMARCAD1 (UNC5C) | **BMPR1B**: Ovarian function/fecundity (Shokrollahi & Morammazi, 2018; Mulsant et al., 2001; Souza et al., 2001; Wilson et al., 2001), body size in sheep (Cao et al., 2015) | PDLIM5*: high altitude adaptation in Yaks (Qi et al., 2019) |
| | | 36835876-36949805 | | | **HERC6**/PPM1K (HERC5/ABCG2) | **HERC6**/ABCG2**:** milk production (Yurchenko et al., 2019, Do et al., 2017) | **HERC6**: adaptation to climate variables in South African goats (Mdladla, 2016), associated with stress grazing tolerance in goats (Mwacharo et al., 2018); associated with arid adaptation in Chinese sheep (Yang et al., 2016) |
| | | 45415286-45625340 | | | **SLC34A2** (ANAPC4/SEL1L3) | SEL1L3: milk production (Sanchez et al., 2007) | **SLC34A2:** adaptation to high altitude in cattle (Verma et al., 2018)<br>SEL1L3*: high altitude adaptation in Ethiopian sheep (Edea, Dadi, Dessie, & Kim, 2019) |
| | | 69082499-69758377 | | | **SCFD2**/LNX1 (FIP1L1/RASL11B) | **SCFD2:** milk constitution (Yodklaew et al., 2017), involved in lipid metabolism (Glastonbury et al., 2016) | |
| | | 85991683-86155374 | | | **CSN1S1/CSN2** | **CSN1S1/CSN2:** structural genes of ruminant milk (see | |

| | | | | | | | |
|---|---|---|---|---|---|---|---|
| | | | | | | Martin, Szymanowska, Zwierzchowski, & Leroux, 2002) | |
| | | 116501734-117144177 | | | **CTBP1/**FAM193A/TNIP2/NOP14/GRK4/HTT (ADD1/SH3BP2/MFSD10) | **CTBP1**: lipid storage, fat cell regulation, fat tail deposition (Xu et al., 2017) | FAM193A: slope adaptation in Moroccan sheep (Benjelloun, 2015) |
| 7 | 1 | 59624626-60848321 | | | **SIL1/CTNNA1**/UBE2D2/CDC25C/LOC108636363 (TMEM173/ECSCR/DNAJC18/MATR3/SLC23A1/PAIP2/LRRTM2) | **CTNNA1/SIL1:** associated with fecundity (Lai et al., 2016), fat tail deposition in sheep (Ahbara et al., 2019)<br>UBE2D2: fat tail deposition in sheep (Mastrangelo et al., 2019)<br>**CTNNA1:** skeletal muscle (Sadkowski et al., 2008) | **SIL1:** associated with plateau condition in Chinese sheep (Yang et al., 2016)<br>**CTNNA1*:** high altitude adaptation in Yaks (Qi et al., 2019)<br>TMEM173: associated with desert condition in Chinese sheep (Yang et al., 2016) |
| 8 | 2 | 74894754-75321840 | | | **UBE2R2/UBAP2/**NOL6 (AQP7/AQP3/DCAF12) | | **UBE2R2/UBAP2:** associated with hypoxia tolerance in humans (Udpa et al., 2014); associated with high altitude condition in Chinese sheep (Yang et al., 2016)<br>AQP7: high altitude adaptation in Yaks (Qi et al., 2019)<br>AQP7/AQP3*: implicated in thermal adaptation (Wollenberg Valero et al., 2014) |
| | | 31846675-32220312 | | | Intergenic zone close to **TYRP1** | **TYRP1**: coat color (Gratten et al., 2007; Becker et al., 2015) | **TYRP1***: associated with desert condition in Chinese sheep (Yang et al., 2016) |
| 10 | 3 | 49806948-50111884 | | | **TCF12** | **TCF12:** associated with litter size (Tao et al., 2013), lactation (Ozdemir Ozgenturk et al., 2017), muscle lipid composition (Dunner et al., 2013) | **TCF12:** adaptation to climate variables in South African goats (Mdladla, 2016) |
| | | 61579137-63362001 | | | **MDGA2** (RPL10L) | | **MDGA2:** adaptation to climate variables in South African goats (Mdladla, 2016) |
| | | 71545588-71700461 | | | Intergenic between **DPH6** and C10H15ORF41 | | **DPH6**: under selection in northern European cattle (Stronen et al., 2019); under selection in Yaks (Lan et al., 2018); associated with plateau condition in Chinese sheep (Yang et al., 2016) |
| 11 | 3 | 37793580-38186897 | | | **LOC106502619/**PNPT1/CFAP36 | | |

| | | | | | | | |
|---|---|---|---|---|---|---|---|
| | | | | | (SEC4R3B/EFEMP1) | | |
| | | 48845878-49036557 | | | **USP39/MAT2A** | | |
| | | 61247655-61456836 | | | **WDPCP** (EHBP1/OTX1) | **WDPCP**: regulates adipogenesis (Sazzini et al., 2016) | **WDPCP**: high altitude adaptation in Ethiopian sheep (Edea et al., 2019) and in Tibetan chicken (Wang et al., 2015)<br>EHBP1: associated with desert condition in Chinese sheep (Yang et al., 2016) |
| 12 | 4 | 43598089-44471339 | | | Intergenic zone between **KLHL1** and **PCDH9** | **KLHL1**: associated with milk yield and lactation persistence (Yue et al., 2017), related to neuron motion and neuromuscular process (Shin et al., 2014)<br>**KLHL1 and PCDH9:** associated with fat tail deposition (Mastrangelo et al., 2019) | **KLHL1**: associated with high altitude adaptation in Chinese sheep (Yang et al., 2016), under selection in human Tibetan (Wang et al., 2011)<br>**PCDH9:** under selection in goat and sheep of arid environment (Kim et al., 2016) |
| | | 48839256-51252236 | | | **RNF17/ZMYM2/PARP4/XPO4/** PSPC1/CRYL1/IFT88/CENPJ/EEF1AKMT1 (ZMYM5/GJB2/IL17D/MPHOSPH8/ATP12A) | **RNF17**: associated with fatty acid composition (Lemos et al., 2016) and growth traits (Edea et al., 2018, Puig-Oliveras et al., 2014)<br>**PARP4**: influences carcass weight (Edea et al., 2018)<br>MPHOSPH8/PARP4/CENPJ/RNF17: associated with growth rate (Mészáros et al., 2019)<br>MPHOSPH8: up regulated in atretic follicles (Hatzirodos et al., 2014)<br>IFT88: growth and feed intake ( Yurchenko et al., 2019) | **RNF17:** adaptation to climate variables in South African goats (Mdladla, 2016)<br>ATP12A: connected to APT mechanisms and high altitude, adaptation in goats (Wang et al. 2016); under selection in sheep of arid environment (Kim et al., 2016)<br>**ZMYM2*/PARP4/**ZMYM5/PSPC1/CENPJ/GJB2: high altitude adaptation in Ethiopian sheep (Edea et al., 2019)<br>GJB2: under selection in goat of arid environment (Kim et al., 2016), associated with high altitude adaptation in Tibetan sheep (Wei et al., 2016) and environmental adaptation via DNA repair mechanisms (Edea et al., 2014)<br>**XPO4**/CRYL1/CENPJ/**RNF17**/MPHOSPH8/**PARP4**: associated with stress grazing tolerance in goats (Mwacharo et al., 2018)<br>IFT88/XPO4: under selection in Moroccan goat (Benjelloun, 2015) |
| | | 60637258-60950668 | | | **NBEA** (MAB21L1) | **NBEA**: wool trait (Wang et al., 2014) | **NBEA**: high altitude adaptation in Ethiopian sheep (Edea et al., 2019) and in cattle (Zeng, 2017); associated |

| | | | | | | | |
|---|---|---|---|---|---|---|---|
| | | | | | | | with body temperature regulation in cattle (Howard et al., 2013) under selection in Ugandan and Morocan goat (Onzima et al., 2018, Benjelloun, 2015); associated with desert adaptation and high altitude adaptation in Chinese sheep (Yang et al., 2016), high altitude adaptation in Yaks (Qi et al., 2019) |
| | | 62109741-62503627 | | | **TRPC4** (POSTN/SUPT20H) | | **TRPC4**: associated with resistance to *Haemonchus contortus* in goat (Corley et al. 2013), implicated in body temperature regulation in cattle (Howard et al., 2013) |
| 13 | 1 | 46458560-46473488 | | | **PRNP** | | **PRNP***: associated with susceptibility to prion disease in sheep and goat (Greenlee, 2019 for a review) |
| 14 | 1 | 16942897-17652266 | | | **VPS13B** | **VPS13B**: milk-related trait (Liu et al., 2018); associated with leg morphology, related with fertility and milk production (Capitan et al., 2014) | **VPS13B**: under selection in Morocan goat (Benjelloun, 2015); associated with arid adaptation in Chinese sheep (Yang et al., 2016), high altitude adaptation in Yaks (Qi et al., 2019) |
| 15 | 2 | 6473101-6606528 | | | **TRPC6/** ANGPTL5 (CEP126) | | **TRPC6*:** high altitude adaptation in Tibetan highlanders (Deng et al., 2019) |
| | | 30213116-30556088 | | | **FCHSD2/**ARHGEF17 (P2RY6/P2RY2) | | |
| 18 | 2 | 36054355-37092141 | | | **NFATC3**/NUFT2/SMPD3/PRMT7/SLC7A6/PSKH1/CENPT (SLC12A4) | | |
| | | 39636636-39980690 | | | **LOC108637979** (ZFHX3/PMFB1) | | |
| 19 | 1 | 32969117-33274858 | | | **PIGL/**LOC102188626**/NCOR1**/CENPV (ADORA2B) | | **PIGL**/CENPV: associated with high altitude in Chinese sheep (Yang et al., 2016)<br>**NCOR1** belong to clock circadian gene network in cattle (Wang et al., 2015)<br>ADORA2B*: implicated in thermal adaptation (Wollenberg Valero et al., 2014) |
| 20 | 1 | 13992024-14204949 | | | **ADAMTS6/**CENPK (PPWD1) | **ADAMTS6:** involved in growth traits in pig (Wu et al. 2019) | **ADAMTS6**/CENPK: associated with high altitude adaptation in Chinese sheep (Yang et al., 2016) |
| 22 | 2 | 28867395-29079702 | | | **SHQ1** (RYBP/PPP4R2/GXYLT2) | **SHQ1**/PPP4R2**:** associated with milk production | **SHQ1**: associated with plateau and desert condition in Chinese sheep (Yang et al., 2016) |

| | | | | | | | |
|---|---|---|---|---|---|---|---|
| | | | | | | (Pasandideh et al., 2018; Stella et al., 2010) | |
| | | 33594470-34062919 | | | **SUCLG2** | **SUCLG2 :** associated with milk production (Di Gerlando et al., 2019), associated with fat deposition in cattle (Silva-Vignato et al., 2019), associated with growth in pig (Yang et al., 2012) | **SUCLG2* :** under selection in sheep of arid environment (Kim et al. 2016); associated with desert condition in Chinese sheep (Yang et al., 2016), high altitude adaptation in Yaks (Qi et al., 2019) |
| 23 | 2 | 36461204-36962562 | | | **BTBD9**/ZFAND3 | | **BTBD9**: adaptation to climate variables in South African goats (Mdladla, 2016)<br>ZFAND3: associated with high altitude condition in Chinese sheep (Yang et al., 2016) |
| | | 39855572-40059589 | | | **UHRF1BP1/C23H6ORF106 (**SNRPC) | | **UHRF1BP1** high altitude adaptation in Yaks ( Qi et al., 2019) |
| 25 | 1 | 30481819-30744262 | | | **AUTS2** | | |
| 26 | 1 | 28596734-29498263 | | | **FBXW4/BTRC/GBF1/**PSD/C26H10ORF76/DPCD (MFSD13A/ACTR1A/NFKB2/PITX3) | **BTRC:** associated with milk fatty acid composition (Palombo et al., 2018); associated with milk production (Paten, 2014); genes associated with fatty acid composition of pork (Viterbo et al., 2018) | **FBXW4/BTRC:** associated with desert in Chinese sheep (Yang et al., 2016)<br>**BTRC:** adaptation to climate variables in goats (Mdladla, 2016), associated with high altitude condition in Chinese sheep (Yang et al., 2016) |

*belong to the hypoxia regulated proteins database (http://www.hypoxiadb.com)

| Chr | Numb. of signatures | Signature range | Identified PCAdapt | Identified LFMM | Gene(s) identified<br>In bold gene(s) with the highest probability of being under selection | Agronomic trait/function reported in the literature for domestic mammals | Literature on genes identified in relation to environmental adaptation in mammals and birds |
|---|---|---|---|---|---|---|---|
| 1 | 3 | 117132460-117619369 | | | CD247/MAEL/**POU2F1**/ LOC105612997/RCSD1 | **POU2F1**: associated with lactase persistence ( Lewinsky et al., 2005); involved in the expression of beta-casein in mammary gland (Zhao et al., 2002), involved in growth and fat deposition in pig ( Pérez-Montarelo et al., 2014) | **POU2F1**: participate in the cellular response to hypoxia in human (Choi et al., 2005; Yan et al., 1999)<br>RCSD1: association with endoparasite phenotypes in cattle (Twomey et al., 2019) |
| | | 170496673-170755505 | | | **BBX** | **BBX:** would be associated with supernumerary nipple in sheep (Peng et al., 2017) | |
| | | 198199858-198342295 | | | **MASP1**/RTP1 (LOC105602729) | | |
| 2 | 2 | 74396276-74749751 | | | **KDM4C** | **KDM4C:** affect reproduction (Guan et al., 2016) | |
| | | 203734535-204610983 | | | **NBEAL1/**ICA1L/ FAM117B (BMPR2) | **NBEA**: wool trait (Wang et al. 2014)<br>ICA1L/FAM117: associated with the pygmy phenotype (Pemberton et al., 2017)<br>BMPR2: involved in ovulation rate and litter size in sheep (Jansson, 2014) | **NBEA**: high altitude adaptation in Ethiopian sheep (Edea et al., 2019) and in cattle (Zeng, 2017); associated with body temperature regulation in cattle (Howard et al., 2013) under selection in Ugandan and Morocan goat (Onzima et al., 2018, Benjelloun, 2015); associated with desert adaptation and high altitude adaptation in Chinese sheep (Yang et al., 2016), high altitude adaptation in Yaks (Qi et al., 2019)<br><br>BMPR2*: Associated with high Altitude Pulmonary Hypertension in Kyrgyz Highlanders (Iranmehr et al., 2019) and in cattle (Newman et al., 2011), associated with desert adaptation and plateau adaptation in Chinese sheep (Yang et al., 2016) |

| | | | | | | | |
|---|---|---|---|---|---|---|---|
| 3 | 1 | 153576602-154585886 | | | **MSRB3**/LOC105609946/WIF1 | **MSRB3:** associated with hear phenotype (Wei et al., 2015, Chen et al., 2018, Webster et al., 2015), associated with fat deposition (Mastrangelo et al., 2019) | **MSRB3:** high altitude adaptation in Ethiopian sheep (Edea et al., 2019) and in dogs (Gou et al., 2014); identified as being under selection in Tibetan sheep (Wei et al. 2016), high altitude adaptation in Yaks (Qi et al., 2019) |
| 4 | 3 | 24727308-25337674 | | | **ISPD**/TSPAN13 | | |
| | | 43272227-43454930 | | | **MAGI2** | | **MAGI2*:** high altitude adaptation in Ethiopian sheep (Edea et al., 2019); associated with desert and arid adaptation in Chinese sheep (Yang et al., 2016), high altitude adaptation in Yaks (Qi et al., 2019) |
| | | 47614996-48783775 | | | COG5/DUS4L/**BCAP29**/**SLC26A4**/CBLL1 (PIK3CG/HBP1/PRKAR2B/GPR22/SLC26A3) | SLC26A4: associated with hearing (Tsukamoto et al., 2003, Park et al., 2003) | **BCAP29/ SLC26A4**: associated with arid adaptation in sheep (Yang et al., 2016)<br>PRKAR2B: associated with plateau adaptation in Chinese sheep (Yang et al., 2016) |
| 5 | 1 | 51252017-51859391 | | | **ARHGAP26**/NR3C1 (TAX1BP1) | **ARHGAP26/**NR3C1*:* affect growth and intra-muscular fatty acid in pig (Edea et al., 2017) | |
| 6 | 1 | 36121839-37756801 (PCAdapt) 37536930-38567860 (LFMM) | | | **LCORL/**HERC3/HERC5/LOC105615454/NCAPG/DCAF16 (FAM184B/HERC6/SPP1/PKD2/LAP3/MED28/PPM1K/ABCG2/MEPE/IBSP) | **LCORL/** LAP3/ FAM184B: growth traits (Al-Mamun et al. 2015, Lindholm-Perry et al., 2001; Bongiorni et al., 2012; Eberlein et al., 2009)<br>ABCG2/SPP1/LAP3/NCAPG/MEPE: milk production (see Ruiz-Larrañaga et al., 2018) | HERC6/SPP1*: associated with arid adaptation in Chinese sheep (Yang et al., 2016)<br>HERC6: adaptation to climate variables in South African goats (Mdladla, 2016), associated with stress grazing tolerance in goats (Mwacharo et al., 2018 );<br>LAP3: under selection in goat of arid environment (Kim et al., 2016) |
| 8 | 1 | 62665338-63271621 | | | **ARFGEF3**/PERP (TNFAIP3) | | **ARFGEF3:** associated with arid adaptation in Chinese sheep (Yang et al., 2016)<br>TNFAIP3*: associated with parasite resistance in cattle (Tay et al., 2018) |

| | | | | | | | |
|---|---|---|---|---|---|---|---|
| 9 | 1 | 34730111-35192293 | | | LOC10114620/**ATP6V1H**/RGS20 (TCEA1/LYPLA1/SOX17) | | **ATP6V1H**: under selection in goat of arid environment (Kim et al., 2016)<br>**ATP6V1H**/SOX17/RGS20/LYPLA1*: associated with humidity condition in goat (Bertolini et al., 2018) |
| 10 | 3 | 29413536-29776019 (PCAdapt) 29479711-29940445 (LFMM) | | | **RXFP2/** BLGALTL | **RXFP2:** involved in horn bud differentiation (Johnston et al., 2011; Allais-Bonnet et al., 2013) | **RXFP2**: associated with latitude, precipitation and temperature in free-ranging bighorn sheep (Roffler et al., 2016); identified as being under selection in Tibetan sheep (Wei et al., 2016). |
| | | 38196759-39403332 (PCAdapt) 37941892-39043416 (LFMM) | | | Intergenic zone before **PCDH9** | **KLHL1**: associated with milk yield and lactation persistence (Yue et al., 2017), related to neuron motion and neuromuscular process (Shin et al., 2014)<br>**KLHL1 and PCDH9:** associated with fat tail deposition (Mastrangelo et al., 2019) | **KLHL1**: associated with high altitude adaptation in sheep (Yang et al., 2016); under selection in Tibetan (Wang et al., 2011)<br>**PCDH9:** under selection in goat and sheep of arid environment (Kim et al., 2016) |
| | | 42194236-43935671 | | | Intergenic zone between **PCDH9** and **KLHL1** | | |
| 12 | 1 | 38573067-39683281 | | | **VPS13D**/TNFSF18 (TNFSF4/TNFRSF1B/DHRS3) | | DHRS3*/TNFRSF1B: associated with arid adaptation in sheep (Yang et al., 2016) |
| 13 | 3 | 48552093-49208171 | | | Intergenic close to **BMP2** | **BMP2**: fat tail deposition (Zhu et al., 2019 ; Yuan et al., 2017 ; Mastrangelo et al., 2019) | **BMP2**: associated with arid condition in Chinese sheep (Yang et al., 2016); under selection in sheep of arid environment (Kim et al., 2016); identified as being under selection in Tibetan sheep (Wei et al., 2016); high altitude adaptation in Yaks (Qi et al., 2019) |
| | | 55025081-56301025 | | | **CDH26**/FAM217B/PPP1R3D (SYCP2/CDH4/EDN3) | EDN3: associated with coat color (Gautier et al. 2015, Kaelin et al., 2012) | EDN3: associated with plateau adaptation in sheep (Yang et al., 2016), slope in Moroccan sheep (Benjelloun 2015); hypoxic adaptation in cetaceans (Tian et al., 2016) |

| | | | | | | | |
|---|---|---|---|---|---|---|---|
| | | | | | | CDH26: associated with fecundity (Lai et al., 2016) | |
| | | 57302923-57809890 | | | **RAB22A**/PMEPA1 | **RAB22A:** implicated in fecundity (Lai et al., 2016) | **RAB22A:** implicated in hypoxia (Wang et al., 2014) (PMEPA1 belong to the hypoxia up-regulated proteins database) |
| 15 | 1 | 59813250-60219075 | | | **DCDC5** (DNAJC24/DCDC1) | **DCDC5**: associated with body conformation (Wu et al., 2013) | DNAJC24: associated with high altitude adaptation in sheep; Heat Shock protein (Yang et al., 2016); under selection in Uganda goat (Onzima et al., 2018) |
| 17 | 1 | 55110665-55329873 | | | **CIT**/TMEM233 (PRKAB1) | | PRKAB1: under selection in goat of arid environment (Kim et al., 2015) |
| 18 | 1 | 66257896-68025345 | | | **TRAF3**/**CDC42BPB**/PPP1R13B (CKB/KLC1/TDRD9/AKT1/TMEM179/SIVA1/ADSSL1/INF2/C18H14orf180) | | TMEM179B: associated with arid condition in sheep (Yang et al., 2016); implicated in hypoxia (Gao et al., 2017) |
| 19 | 1 | 28133753-28962852 | | | **PDZRN3/**GXYLT2 (PPP4R2/SHQ1) | PPP4R2/SHQ1: associated with growth rate (Pasandideh et al., 2018) | SHQ1 associated with plateau and desert condition in sheep (Yang et al., 2016) |
| 21 | 1 | 17885334-18117515 | | | **PAK1** (LOC101117288/AQP11) | | PAK1 associated with plateau condition in sheep (Yang et al., 2016); under selection in sheep of arid environment (Kim et al., 2016) |
| 25 | 1 | 19073058-19380300 | | | **JMJD1C/**NRBF2 (REEP3) | | REEP3 associated with plateau condition in sheep (Yang et al., 2016)<br>**JMJD1C*/**NRBF2/REEP3:associated with high-altitude adaptation in pig (Dong et al., 2014) |

*belong to the hypoxia regulated proteins database (http://www.hypoxiadb.com)

| Chr. | GOAT<br>PCAdapt Signatures | GOAT<br>LFMM Signatures | SHEEP<br>LFMM Signatures | SHEEP<br>PCAdapt Signatures |
|---|---|---|---|---|
| Chr 1 | | **46047752-46411087**<br><br>**snp45739-scaffold627-5315341 46047752***<br>**snp45746-scaffold627-5616708 46348421**<br>**snp45747-scaffold627-5650100 46381611**<br>**snp12636-scaffold1480-1163799 46411087**<br><br>**84242981-84773966**<br><br>**snp32731-scaffold377-2079945 84242981**<br>snp32730-scaffold377-2049881 84272916<br>**snp32729-scaffold377-2002407 84320917**<br>**snp32728-scaffold377-1966001 84357680***<br>snp32723-scaffold377-1745669 84573836<br>snp32721-scaffold377-1639477 84680820<br>**snp32720-scaffold377-1606429 84713972**<br>**snp32719-scaffold377-1576455 84743986**<br>**snp32718-scaffold377-1546443 84773966** | **117132460-117619369**<br><br>**s74762.1 117132460**<br>**OAR1_127060133.1 117619369**<br>**s71952.1 117697523**<br>s49384.1 117771574<br>**OAR1_127291951.1 117853500***<br>**s73007.1 117916288***<br><br>**170496673-170755505**<br><br>**OAR1_183962708.1 170496673**<br>**OAR1_184162321.1 170690492**<br>**s12579.1 170725424**<br>**OAR1_184227318.1 170755505***<br><br>**198199858-198342295**<br><br>**OAR1_214098599.1 198199858**<br>**OAR1_214136810.1 198227861**<br>**OAR1_214203546.1 198286740**<br>**OAR1_214229899.1 198312258**<br>**OAR1_214247090.1 198329788**<br>**OAR1_214257185.1 198342295*** | |
| Chr2 | | | **74396276-74749751**<br><br>**OAR2_79262293_X.1 74396276**<br>**OAR2_79499240.1 74613800**<br>**OAR2_79550682.1 74669837***<br>**OAR2_79594745.1 74714666*** | |

| | | | | |
|---|---|---|---|---|
| | | | s62643.1 74749751*<br><br><u>203734535-204610983</u><br><br>s34105.1 203717963<br>OAR2_215477291.1 203734535<br>OAR2_215596955.1 203761367<br>OAR2_215636399.1 203797311*<br>OAR2_215711808.1 203869407<br>OAR2_216615372.1 204610983 | |
| Chr 3 | | | <u>153576602-154585886</u><br><br>OAR3_164422954.1 153576602<br>s26177.1 153826281*<br>s69653.1 153976304*<br>s69686.1 153996225<br>OAR3_165009241.1 154033734<br>OAR3_165050963.1 154072493<br>OAR3_165200988.1* 154223123<br>s75917.1 154318689*<br>s61003.1 154585886 | <u>153889169-154318689</u><br><br>OAR3_164788310.1 153889169<br>s16949.1 153927239<br>s69653.1 153976304<br>s69686.1 153996225<br>OAR3_165009241.1 154033734<br>OAR3_165050963.1 154072493*<br>s75917.1 154318689 |
| Chr 4 | | | <u>24727308-25337674</u><br><br>OAR4_26039959.1 24727308*<br>OAR4_26021918.1 24745112*<br>OAR4_26202538_X.1 24891467*<br>OAR4_26645098.1 25337674<br><br><u>43272227-43454930</u><br><br>OAR4_45683707.1 43272227<br>OAR4_45746457.1 43333090* | |

| | | | | |
|---|---|---|---|---|
| | | | **s36928.1 43433914**<br>**OAR4_45871691.1 43454930***<br><br>47614996-48783775<br><br>**OAR4_50346558.1 47614996**<br>s34975.1 47852019<br>OAR4_50737323.1 47928357<br>**s24500.1 47960510**<br>**s39886.1 48085823**<br>**s32219.1 48284042**<br>**OAR4_51241289.1 48402666**<br>**OAR4_51315739.1 48480244**<br>**OAR4_51346813.1 48510936**<br>**OAR4_51441757.1 48607563**<br>**OAR4_51489408.1 48649775***<br>**s46022.1 48737779***<br>**OAR4_51625352.1 48783775** | |
| Chr 5 | | 36576428-37074733<br><br>**snp9058-scaffold1329-759519 36576428**<br>**snp9059-scaffold1329-790608 36606034**<br>**snp9062-scaffold1329-936768 36750654**<br>**snp9063-scaffold1329-976103 36790914***<br>snp9066-scaffold1329-1128684 36939084<br>**snp12728-scaffold149-35530 37074733** | 51252017-51859391<br><br>**s44340.1 51252017***<br>**OAR5_55651844.1 51284980**<br>**OAR5_55827200.1 51367486**<br>**OAR5_55949772.1 51459909***<br>OAR5_56305688.1 51818687<br>**OAR5_56351796.1 51859391*** | |
| Chr 6 | 36835876-36949805<br><br>**snp26737-scaffold281-187160 36835876**<br>**snp26738-scaffold281-221127 36870090** | 19677067-20300302<br><br>snp49755-scaffold710-1594826 19677067<br>**snp49753-scaffold710-1514064 19757830** | 37536930-38567860<br><br>**OAR6_41850329.1 37536930**<br>OAR6_42094768.1 37756801 | 36121839-37756801<br><br>OAR6_40409402.1 36121839<br>**OAR6_40496376.1 36210335** |

| | | | | |
|---|---|---|---|---|
| | **snp26739-scaffold281-267352 36916955***<br>**snp26740-scaffold281-299943 36949805**<br><br>**85991683-86155374**<br><br>**snp59426-scaffold980-307288 85991683**<br>**snp60000-CSN1S1-ex17 85993405**<br>**snp59427-scaffold980-309132 85994156**<br>**snp59428-scaffold980-310410 85995436**<br>**snp59429-scaffold980-310523 85995549**<br>**snp59430-scaffold980-311053 85996079**<br>**snp59432-scaffold980-311547 85996571**<br>**snp59434-scaffold980-322977 86007956***<br>**snp59438-scaffold980-323773 86008752***<br>**snp59439-scaffold980-324311 86009290***<br>**snp59440-scaffold980-324828 86009807***<br>**snp59441-scaffold980-325353 86010332***<br>snp59450-scaffold980-400667 86085897<br>**snp59451-scaffold980-402598 86087828**<br>**snp59452-scaffold980-402711 86087941**<br>**snp59454-scaffold980-403305 86088536**<br>**snp59455-scaffold980-407817 86093124**<br>snp59458-scaffold980-434555 86118732<br>snp59459-scaffold980-469488 86155374 | **snp49748-scaffold710-1271522 20001811**<br>**snp49747-scaffold710-1236741 20036645***<br>snp49744-scaffold710-1107424 20167200<br>**snp49743-scaffold710-1073847 20200941***<br>**snp49742-scaffold710-1035063 20239512**<br>snp49741-scaffold710-1002906 20271740<br>**snp49740-scaffold710-974383 20300302**<br><br>**30139296-30874402**<br><br>**snp59004-scaffold968-1784874 30139296**<br>**snp59005-scaffold968-1816117 30170190***<br>snp59007-scaffold968-1927374 30282214<br>**snp59013-scaffold968-2181677 30541674**<br>**snp59020-scaffold968-2472589 30831824***<br>**snp59021-scaffold968-2515154 30874402***<br><br>**36835876-36949805**<br><br>**snp26737-scaffold281-187160 36835876**<br>**snp26738-scaffold281-221127 36870090**<br>**snp26739-scaffold281-267352 36916955 ***<br>**snp26740-scaffold281-299943 36949805***<br>**snp26741-scaffold281-332466 36982221**<br><br>**45415286-45625340**<br><br>**snp12879-scaffold1498-8679 45415286**<br>**snp12880-scaffold1498-47642 45452714***<br>snp12881-scaffold1498-89673 45494742<br>**snp12882-scaffold1498-142431 45547830**<br>snp12883-scaffold1498-190653 45596401<br>**snp12884-scaffold1498-220976 45625340**<br><br>**69082499-69758377** | OAR6_42208195.1 37865761<br>**OAR6_42247197.1 37911182**<br>**OAR6_42557643.1 38214635***<br>**OAR6_42743614.1 38345613**<br>**OAR6_42930840.1 38480285**<br>**OAR6_43034224.1 38567860*** | **OAR6_41003295.1 36747507***<br>**DU178311_404.1 36768153**<br>OAR6_41424992.1 37126564<br>**OAR6_41476497.1 37177538**<br>**OAR6_41494878.1 37198398***<br>**OAR6_41558126.1 37257076**<br>**OAR6_41583796.1 37282110**<br>**OAR6_41709987.1 37400993**<br>OAR6_41768532.1 37455134<br>**OAR6_41936490.1 37619726**<br>OAR6_42094768.1 37756801 |

| | | |
|---|---|---|
| | | snp58054-scaffold94-3761495 69082499<br>**snp58060-scaffold94-3954937 69276841**<br>**snp58062-scaffold94-4076449 69398715**<br>**snp58063-scaffold94-4115460 69437556**<br>snp58065-scaffold94-4243148 69566293<br>**snp58069-scaffold94-4434969 69758377***<br><br>**<u>116501734-117144177</u>**<br><br>**snp16110-scaffold1698-5528 116501734**<br>snp16112-scaffold1698-77702 116572175<br>snp16113-scaffold1698-116152 116608825<br>snp16116-scaffold1698-266267 116756644<br>**snp16117-scaffold1698-297420 116787929***<br>snp16120-scaffold1698-458655 116929275<br>**snp29345-scaffold3166-28105 117121013**<br>**snp29346-scaffold3166-62970 117131241**<br>**snp38582-scaffold4894-9087 117144177** |
| Chr 7 | **<u>59624626-60848321</u>**<br><br>**snp30691-scaffold339-5649390 59624626**<br>**snp30688-scaffold339-5531611 59742326**<br>**snp30687-scaffold339-5490945 59782859**<br>snp30685-scaffold339-5400884 59873465<br>**snp30682-scaffold339-5258982 60015475**<br>**snp30681-scaffold339-5230221 60044210**<br>**snp30680-scaffold339-5200334 60074382**<br>**snp30679-scaffold339-5167374 60107657***<br>**snp30678-scaffold339-5138454 60136996**<br>snp30677-scaffold339-5071691 60205663<br>**snp30675-scaffold339-4996663 60280996**<br>**snp30674-scaffold339-4962908 60315027**<br>**snp30673-scaffold339-4924262 60353593**<br>**snp30672-scaffold339-4888154 60388662** | |

| | | | |
|---|---|---|---|
| | **snp30670-scaffold339-4796176 60480852**<br>**snp30669-scaffold339-4761615 60515404**<br>**snp30667-scaffold339-4673976 60603707**<br>**snp30663-scaffold339-4429271 60848321** | | |
| Chr 8 | **74894754-75321840**<br><br>snp10557-scaffold1376-1305714 74894754<br>**snp10556-scaffold1376-1245744 74954737**<br>**snp10555-scaffold1376-1193823 75006929**<br>**snp10554-scaffold1376-1159382 75041455***<br>**snp10553-scaffold1376-1118888 75082828**<br>snp10552-scaffold1376-1089536 75112213<br>**snp10551-scaffold1376-1054710 75147070**<br>**snp10550-scaffold1376-1023594 75178621**<br>**snp10549-scaffold1376-984757 75217436**<br>**snp10548-scaffold1376-951770 75250756***<br>**snp10546-scaffold1376-880604 75321840** | **74894754-75321840**<br><br>**snp10557-scaffold1376-1305714 74894754**<br>**snp10556-scaffold1376-1245744 74954737***<br>**snp10554-scaffold1376-1159382 75041455**<br>snp10553-scaffold1376-1118888 75082828<br>snp10551-scaffold1376-1054710 75147070<br>**snp10548-scaffold1376-951770 75250756**<br>snp10546-scaffold1376-880604 75321840<br><br>**31846675-32220312**<br><br>**snp3828-scaffold1121-278677 31846675***<br>**snp3825-scaffold1121-173930 31952477***<br>**snp3824-scaffold1121-138568 31987692**<br>snp3823-scaffold1121-95838 32030671<br>**snp3822-scaffold1121-66026 32061108**<br>snp3821-scaffold1121-32831 32094669<br>snp28582-scaffold3060-91473 32220312 | **62665338-63271621**<br><br>**OAR8_67529714.1 62665338***<br>**s54005.1 62712894**<br>**OAR8_67633238.1 62766989**<br>**s21394.1 62941437***<br>**OAR8_68132928.1 63271621** |
| Chr 9 | | | **34730111-35192293**<br><br>**OAR9_36726320.1 34730111***<br>**s73611.1 34779773***<br>**OAR9_36817104.1 34825201**<br>**OAR9_37077743.1 35066451**<br>s08861.1 35166051<br>**OAR9_37208898.1 35192293** |

| | | | | |
|---|---|---|---|---|
| Chr 10 | **49806948-50111884**<br><br>snp48056-scaffold68-2052280 49806948<br>**snp48057-scaffold68-2081562 49836347***<br>**snp48058-scaffold68-2125826 49880369***<br>**snp48059-scaffold68-2159758 49914964**<br>snp48061-scaffold68-2277132 50032968<br>snp48062-scaffold68-2321770 50077763<br>**snp48063-scaffold68-2355884 50111884** | **61579137-63362001**<br><br>**snp33144-scaffold388-638006 61579137**<br>**snp33150-scaffold388-885658 61834915**<br>snp33163-scaffold388-1462914 62410197<br>**snp33166-scaffold388-1627736 62575526 ***<br>snp33168-scaffold388-1694651 62643336<br>**snp11201-scaffold1400-366987 62900522**<br>**snp11198-scaffold1400-225713 63042347**<br>**snp24945-scaffold2554-7579 63321283**<br>snp33726-scaffold397-1169 63330652<br>**snp33727-scaffold397-32542 63362001**<br><br>**71545588-71700461**<br><br>**snp31014-scaffold343-340685 71545588**<br>**snp31015-scaffold343-388340 71594906**<br>**snp31016-scaffold343-426662 71633591***<br>**snp31017-scaffold343-492910 71700461** | **29479711-29940445**<br><br>**OAR10_29538398.1 29479711***<br>**OAR10_29722772.1 29660838**<br>**OAR10_29737372.1 29685665**<br>**OAR10_29793750.1 29742016**<br>**s18834.1 2977601***<br>**OAR10_29867192.1 29812108**<br>**OAR10_30003563.1 29940445**<br>s47229.1 30099193<br>s73573.1 30201353<br><br>**37941892-39043416**<br><br>**OAR10_38772235.1 37941892**<br>**OAR10_38966500.1 38196759**<br>**OAR10_39356591.1 38481236***<br>**OAR10_39377612.1 38504099**<br>**OAR10_39484055.1 38597569***<br>**OAR10_39526423_X.1 38639728***<br>OAR10_39573000.1 38685897<br>s32389.1 38717908<br>**OAR10_39630287.1 38741781**<br>**OAR10_39841998.1 39015113**<br>**OAR10_39875924.1 39043416** | **29413536-29776019**<br><br>**OAR10_29469450.1 29413536**<br>**OAR10_29511510.1 29453722**<br>**OAR10_29538398.1 29479711***<br>**OAR10_29546872.1 29489616**<br><br>**38196759-39403332**<br><br>OAR10_38772235.1 37941892<br>**OAR10_38966500.1 38196759**<br>**OAR10_39000561.1 38216203**<br>**OAR10_39356591.1 38481236**<br>**OAR10_39377612.1 38504099***<br>**OAR10_39484055.1 38597569**<br>**OAR10_39526423_X.1 38639728***<br>**OAR10_39573000.1 38685897***<br>**OAR10_39630287.1 38741781**<br>**OAR10_39756558.1 38938375**<br>OAR10_39841998.1 39015113<br>**OAR10_39875924.1 39043416**<br>**OAR10_39900534.1 39061704**<br>**OAR10_40027965.1 39201371**<br>OAR10_40036146.1 39202994<br>OAR10_40078675.1 39246697<br>**OAR10_40150757.1 39312880**<br>**OAR10_40237265.1 39403332**<br><br>**42194236-43935671**<br><br>**OAR10_43095152.1 42194236**<br>**OAR10_43161187.1 42263285**<br>OAR10_43195785.1 42297892<br>**OAR10_43246818.1 42358254***<br>**OAR10_43391432.1 42454938** |

| | | | | |
|---|---|---|---|---|
| | | | | **OAR10_43407475.1 42468525**<br>**OAR10_43441368.1 42512611**<br>**OAR10_43550150.1 42650662**<br>**OAR10_43601954.1 42713204**<br>**OAR10_43626984_X.1 42735479**<br>**OAR10_44407983.1 43782188**<br>**OAR10_44470993.1 43845305**<br>**OAR10_44569028.1 43935671** |
| Chr 11 | **37793580-38186897**<br><br>**snp24199-scaffold247-290385 37793580**<br>**snp24200-scaffold247-348793 37852358***<br>**snp24201-scaffold247-416001 37920009***<br>**snp24202-scaffold247-451263 37955407**<br>**snp24203-scaffold247-491893 37996657***<br>**snp24204-scaffold247-523077 38027819***<br>**snp24205-scaffold247-567391 38072888***<br>**snp24206-scaffold247-602548 38108083**<br>**snp24208-scaffold247-681263 38186897**<br><br>**61247655-61456836**<br><br>**snp18100-scaffold185-12622205 61247655***<br>**snp18099-scaffold185-12591335 61278930***<br>**snp18098-scaffold185-12562590 61308199**<br>**snp18097-scaffold185-12494429 61379162**<br>snp18096-scaffold185-12445655 61427940<br>snp18095-scaffold185-12416881 61456836 | **48845878-49036557**<br><br>**snp3591-scaffold1111-454455 48845878**<br>**snp3592-scaffold1111-496847 48888368**<br>**snp3594-scaffold1111-599861 48991424***<br>**snp3595-scaffold1111-644798 49036557** | | |
| Chr 12 | **43598089-44471339** | **60637258-60950668** | **38573067-39683281** | |

| | | | |
|---|---|---|---|
| | snp55336-scaffold855-17875 43598089<br>**snp53668-scaffold817-874431 43916255**<br>**snp53666-scaffold817-776660 44012257**<br>**snp53665-scaffold817-739382 44050377**<br>**snp53664-scaffold817-708766 44081824***<br>**snp53663-scaffold817-677742 44113302**<br>**snp53662-scaffold817-625200 44165707***<br>**snp53661-scaffold817-570565 44221034**<br>snp53660-scaffold817-516359 44275453<br>**snp53659-scaffold817-464982 44327480**<br>**snp53657-scaffold817-364334 44429883**<br>**snp53656-scaffold817-324137 44471339**<br><br>**<u>48839256-51252236</u>**<br><br>**snp11147-scaffold140-984058 48839256**<br>**snp11146-scaffold140-931318 48893166**<br>snp11142-scaffold140-760668 49066547<br>snp11141-scaffold140-711242 49116716<br>**snp11135-scaffold140-435808 49396564**<br>snp11128-scaffold140-65176 49775064<br>**snp11127-scaffold140-35215 49805838**<br>**snp11126-scaffold140-2873 49837948**<br>**snp30388-scaffold335-708 49846240**<br>**snp30389-scaffold335-38638 49884682**<br>**snp30390-scaffold335-79561 49926025**<br>snp30391-scaffold335-171300 50018490<br>**snp30392-scaffold335-200469 50047466**<br>**snp30393-scaffold335-255434 50102642**<br>**snp30394-scaffold335-315825 50152165**<br>**snp30395-scaffold335-354367 50190714**<br>**snp30396-scaffold335-385244 50221592**<br>**snp30397-scaffold335-418126 50253401**<br>**snp30398-scaffold335-469650 50304848**<br>**snp30399-scaffold335-501928 50327948***<br>**snp30400-scaffold335-530813 50356882**<br>**snp30402-scaffold335-625380 50451600*** | **snp50177-scaffold717-4519853 60637258***<br>**snp50175-scaffold717-4447356 60710546**<br>**snp50172-scaffold717-4328533 60829727**<br>snp50171-scaffold717-4286071 60872237<br>**snp50170-scaffold717-4247907 60910543***<br>**snp50169-scaffold717-4207960 60950668***<br><br>**<u>62109741-62503627</u>**<br><br>**snp50143-scaffold717-3039794 62109741***<br>**snp50140-scaffold717-2929052 62221458**<br>**snp50138-scaffold717-2864241 62286750**<br>**snp50133-scaffold717-2645969 62503627** | **OAR12_43059012.1 38573067**<br>**OAR12_43069135.1 38584294**<br>**OAR12_43331187.1 38845652***<br>**OAR12_43408733.1 38925278**<br>OAR12_43836490.1 39266000<br>**OAR12_44013347.1 39430517**<br>**OAR12_44173454.1 39590218**<br>**s01697.1 39683281** |

| | | | | |
|---|---|---|---|---|
| | **snp30403-scaffold335-692736 50515766***<br>**snp30404-scaffold335-730262 50553986**<br>**snp30405-scaffold335-772607 50596847**<br>**snp30406-scaffold335-807385 50631622**<br>**snp30407-scaffold335-852300 50676638**<br>**snp30410-scaffold335-988243 50813753**<br>**snp30411-scaffold335-1024805 50849395**<br>**snp30412-scaffold335-1063925 50888523**<br>**snp30413-scaffold335-1113038 50937829**<br>snp30414-scaffold335-1149164 50971205<br>**snp30415-scaffold335-1248178 51070302**<br>snp30416-scaffold335-1279037 51101199<br>snp30419-scaffold335-1430456 51252236<br><br>**60391688-60950668**<br><br>snp50183-scaffold717-4762486 60391688<br>snp50182-scaffold717-4723901 60430663<br>snp50181-scaffold717-4685676 60468859<br>snp50179-scaffold717-4583525 60571662<br>snp50178-scaffold717-4551817 60605463<br>**snp50175-scaffold717-4447356 60710546**<br>**snp50172-scaffold717-4328533 60829727**<br>snp50171-scaffold717-4286071 60872237<br>**snp50170-scaffold717-4247907 60910543***<br>**snp50169-scaffold717-4207960 60950668*** | | | |
| Chr 13 | **46458560-46473488**<br><br>**snp56302-scaffold881-1647723 46458560**<br>**snp56304-scaffold881-1648328 46459167**<br>**snp56308-scaffold881-1657525 46468393**<br>**snp56310-scaffold881-1658893 46469761**<br>**snp56312-scaffold881-1659181 46470049*** | | **48552093-49208171**<br><br>**OAR13_51852034.1 48552093**<br>**OAR13_51886803.1 48585294**<br>**OAR13_52092653.1 48761402**<br>**OAR13_52482285.1 48935908**<br>**OAR13_52630089.1 49070447*** | **48552093-49070447**<br><br>**OAR13_51852034.1 48552093***<br>OAR13_51886803.1 48585294<br>OAR13_52092653.1 48761402<br>**s27419.1 48897111**<br>**OAR13_52482285.1 48935908** |

| | | | | |
|---|---|---|---|---|
| | **snp56315-scaffold881-1659399 46470267**<br>**snp56317-scaffold881-1659485 46470353**<br>**snp56318-scaffold881-1660022 46470890**<br>**snp56319-scaffold881-1662266 46473134**<br>**snp56320-scaffold881-1662295 46473163**<br>**snp56322-scaffold881-1662620 46473488** | | **OAR13_52594105.1 49082829**<br>**OAR13_52640992.1 49137513***<br>**OAR13_52723941.1 49208171**<br><br>**<u>55025081-56301025</u>**<br><br>**s73781.1 55025081**<br>**s44347.1 55413774**<br>**OAR13_60486404.1 55415559**<br>**OAR13_60759835.1 55773534**<br>**s47781.1 55823232**<br>**OAR13_60855392.1 55877764**<br>**OAR13_60893851.1 55918326**<br>**s13640.1 56301025***<br><br>**<u>57302923-57809890</u>**<br><br>**s35407.1 57302923**<br>**s18305.1 57332790**<br>**OAR13_62551597.1 57339112**<br>**s32799.1 57571809**<br>**s63169.1 57809890*** | **OAR13_52630089.1 49070447** |
| Chr 14 | **<u>16942897-17652266</u>**<br><br>snp42629-scaffold566-2157875 16942897<br>**snp42627-scaffold566-2087522 17014833**<br>**snp42626-scaffold566-2037573 17064867***<br>**snp42624-scaffold566-1939716 17163042***<br>**snp42622-scaffold566-1844989 17258805**<br>snp42620-scaffold566-1767368 17334795<br>snp42618-scaffold566-1682208 17420173<br>**snp42617-scaffold566-1618985 17483834**<br>**snp42616-scaffold566-1587537 17515692**<br>**snp42615-scaffold566-1545022 17558651** | | | |

| | | | |
|---|---|---|---|
| | snp42614-scaffold566-1452334 17652266 | | |
| Chr 15 | | <u>**6473101-6606528**</u><br><br>**snp40417-scaffold516-614362 6473101**<br>**snp40416-scaffold516-558833 6528574***<br>**snp40415-scaffold516-526881 6560526***<br>**snp40414-scaffold516-480259 6606528**<br><br><u>**30213116-30556088**</u><br><br>**snp1696-scaffold1048-61993 30213116**<br>**snp41709-scaffold542-3016217 30310244***<br>snp41708-scaffold542-2933956 30393252<br>snp41707-scaffold542-2884418 30441621<br>snp41706-scaffold542-2845412 30481062<br>**snp41705-scaffold542-2802565 30524299**<br>**snp41704-scaffold542-2770868 30556088*** | <u>**59813250-60219075**</u><br><br>**OAR15_65397148.1 59813250**<br>**OAR15_65467094_X.1 59869594**<br>**OAR15_65536256.1 59929380**<br>**OAR15_65584079.1 59981575***<br>**s56927.1 60219075** |
| Chr 17 | | | <u>**55110665-55329873**</u><br><br>**OAR17_60138319.1 55110665***<br>**OAR17_60143832.1 55116388***<br>**OAR17_60225378.1 55179421**<br>**s00902.1 55329873** |
| Chr 18 | <u>**36054355-37092141**</u><br><br>snp22639-scaffold225-669244 36054355<br>snp22636-scaffold225-501605 36221683<br>snp22635-scaffold225-468855 36254528<br>**snp22633-scaffold225-401770 36321627**<br>**snp22630-scaffold225-275173 36448079** | | <u>**66257896-68025345**</u><br><br>**s10898.1 66257896**<br>**s43273.1 66283067**<br>**OAR18_70439577_X.1 66321715***<br>**s59254.1 66841063**<br>DU396708_413.1 66925898 |

| | | | |
|---|---|---|---|
| | **snp22627-scaffold225-125543 36598178***<br>**snp22626-scaffold225-94485 36629376***<br>**snp22625-scaffold225-48106 36674199**<br>**snp7213-scaffold1266-2438708 36745379**<br>**snp7212-scaffold1266-2397446 36786664***<br>**snp7211-scaffold1266-2368797 36815394***<br>**snp7210-scaffold1266-2322163 36862059***<br>**snp7209-scaffold1266-2252567 36931726***<br>**snp7208-scaffold1266-2217995 36966108***<br>**snp7207-scaffold1266-2177748 37006519**<br><br>**39636636-39980690**<br><br>snp9593-scaffold1344-443953 39636636<br>snp9594-scaffold1344-484533 39665961<br>**snp9595-scaffold1344-523130 39704456***<br>**snp9596-scaffold1344-574879 39758374***<br>**snp9597-scaffold1344-624191 39807767***<br>**snp9598-scaffold1344-658729 39842792***<br>**snp9599-scaffold1344-690205 39874628**<br>**snp9600-scaffold1344-741295 39926740**<br>**snp9601-scaffold1344-794509 39980690** | | **s48176.1 67190301**<br>**OAR18_71769962.1 67554481**<br>**s34796.1 67809480***<br>**s03219.1 68025345** |
| Chr 19 | **32969117-33274858**<br><br>snp28431-scaffold303-2935885 32969117<br>**snp28432-scaffold303-2995232 33028814**<br>**snp28433-scaffold303-3044995 33078622**<br>**snp28436-scaffold303-3170123 33202721***<br>**snp28437-scaffold303-3243981 33274858** | | **28133753-28962852**<br><br>**OAR19_29758110.1 28133753***<br>OAR19_29776311.1 28153652<br>**s37139.1 28594822**<br>**s10229.1 28632951**<br>**OAR19_30435043.1 28786157**<br>**OAR19_30535417.1 28962852** |
| Chr 20 | **13992024-14204949**<br><br>**snp43213-scaffold575-1038863 13992024*** | | |

| | | | | |
|---|---|---|---|---|
| | **snp43214-scaffold575-1067749 14021075**<br>**snp43215-scaffold575-1103515 14056838***<br>**snp43216-scaffold575-1149259 14103234***<br>**snp43217-scaffold575-1178652 14132568***<br>**snp43218-scaffold575-1250834 14204949** | | | |
| Chr 21 | | | | **17885334-18117515**<br><br>**OAR21_20255140_X.1 17885334**<br>**s23216.1 17896095***<br>**OAR21_20293826.1 17924936**<br>**OAR21_20301210.1 17931634***<br>**OAR21_20371526.1 18002035**<br>**OAR21_20401829.1 18030662**<br>**s62605.1 18117515** |
| Chr 22 | **28867395-29079702**<br><br>**snp3022-scaffold1091-866051 28867395**<br>**snp3023-scaffold1091-919725 28921320***<br>**snp3024-scaffold1091-953204 28955150***<br>**snp3025-scaffold1091-1001799 29004527***<br>**snp3026-scaffold1091-1047549 29050298**<br>**snp3027-scaffold1091-1076915 29079702**<br><br>**33594470-34062919**<br><br>**snp13803-scaffold154-526016 33594470**<br>snp13800-scaffold154-407557 33709623<br>**snp13799-scaffold154-347555 33769647**<br>**snp13798-scaffold154-300085 33818637**<br>**snp13797-scaffold154-243475 33876201**<br>**snp13796-scaffold154-213942 33905179**<br>**snp13795-scaffold154-182886 33936550**<br>snp13794-scaffold154-139936 33979489 | | | |

| | | | | |
|---|---|---|---|---|
| | **snp13793-scaffold154-98284 34020913***<br>**snp13792-scaffold154-56173 34062919** | | | |
| Chr 23 | <u>36461204-36962562</u><br><br>**snp59312-scaffold975-869095 36461204**<br>**snp59311-scaffold975-838707 36491546***<br>**snp59310-scaffold975-794737 36534387**<br>snp59309-scaffold975-765057 36564031<br>**snp59308-scaffold975-735328 36594106**<br>snp59307-scaffold975-682257 36642064<br>snp59299-scaffold975-362386 36962562 | <u>39855572-40059589</u><br><br>**snp23559-scaffold2376-180998 39855572**<br>**snp23560-scaffold2376-217634 39892708**<br>snp23561-scaffold2376-253763 39928817<br>**snp23562-scaffold2376-312530 39987612***<br>**snp23563-scaffold2376-384575 40059589*** | | |
| Chr 25 | <u>30481819-30744262</u><br><br>**snp9550-scaffold1343-2237381 30481819**<br>**snp9551-scaffold1343-2270352 30514941***<br>snp9553-scaffold1343-2330570 30576113<br>**snp9554-scaffold1343-2381252 30626531**<br>**snp9556-scaffold1343-2459116 30704234**<br>snp9557-scaffold1343-2499155 30744262 | | | <u>19073058-19380300</u><br><br>**OAR25_19823535.1 19073058**<br>**OAR25_19926183.1 19174899**<br>**OAR25_19941944.1 19190707**<br>**OAR25_20026839.1 19273755***<br>**OAR25_20041897.1 19290987***<br>**OAR25_20106030.1 19342453***<br>OAR25_20142347.1 19380300 |
| Chr 26 | <u>28596734-29498263</u><br><br>snp41096-scaffold532-288303 28596734<br>**snp41097-scaffold532-332094 28640088**<br>**snp41098-scaffold532-368969 28677110**<br>**snp41099-scaffold532-409992 28717972**<br>snp41100-scaffold532-448160 28756401<br>snp41101-scaffold532-481203 28788742<br>snp41104-scaffold532-649324 28957011<br>snp41105-scaffold532-692721 29001092<br>snp41109-scaffold532-897192 29205431<br>**snp41110-scaffold532-933054 29241421***<br>**snp41111-scaffold532-965874 29274605*** | | | |

| | |
|---|---|
| | **snp41112-scaffold532-1001039 29309963**<br>snp41113-scaffold532-1051457 29360578<br>**snp41114-scaffold532-1082302 29391742**<br>**snp41117-scaffold532-118800329498263** |

The beginning and end of each signature is indicated in base pairs and underlined; SNPs showing p-value<$10^{-6}$ are indicated for each signature; in bold SNPs showing p-value<$1.10^{-9}$; *SNPs showing highest p-values of the signatures.

**GOAT**

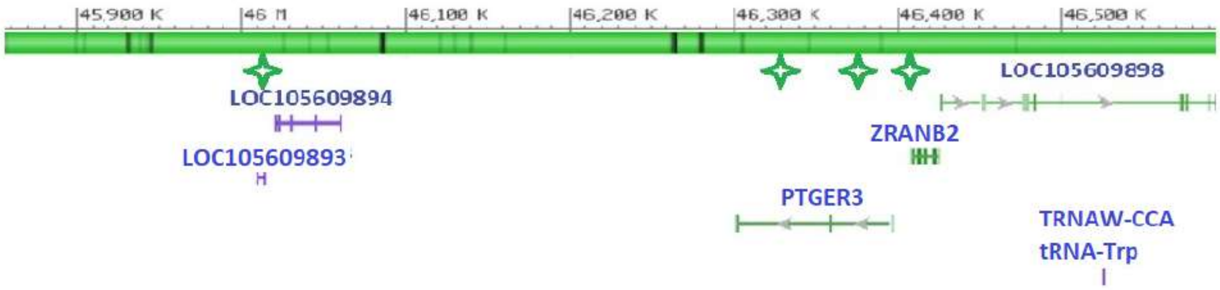
45,900 K
46 M
46,100 K
46,200 K
46,300 K
46,400 K
46,500 K
LOC105609894
LOC105609893
LOC105609898
ZRANB2
PTGER3
TRNAW-CCA
tRNA-Trp

*Chr. 1*

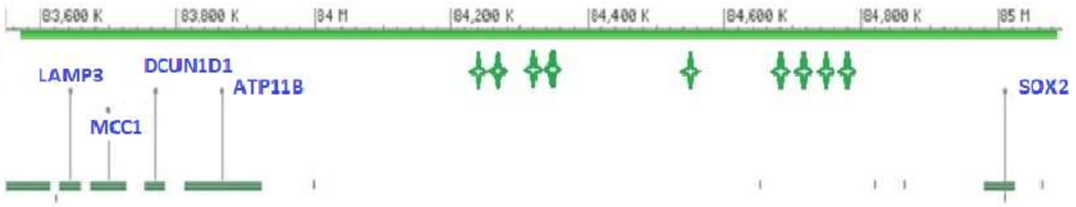
83,600 K
83,800 K
84 M
84,200 K
84,400 K
84,600 K
84,800 K
85 M
LAMP3
DCUN1D1
ATP11B
MCC1
SOX2

**GOAT**

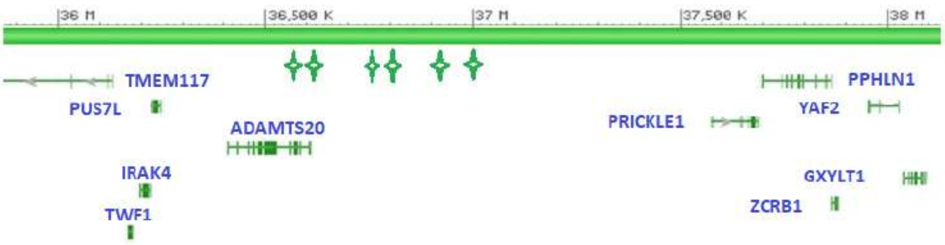

*Chr. 5*

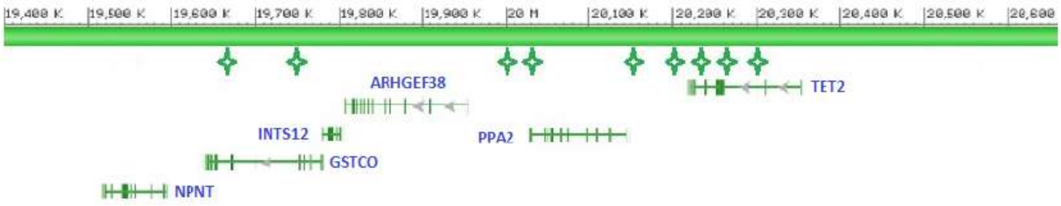

*Chr. 6*

**GOAT**

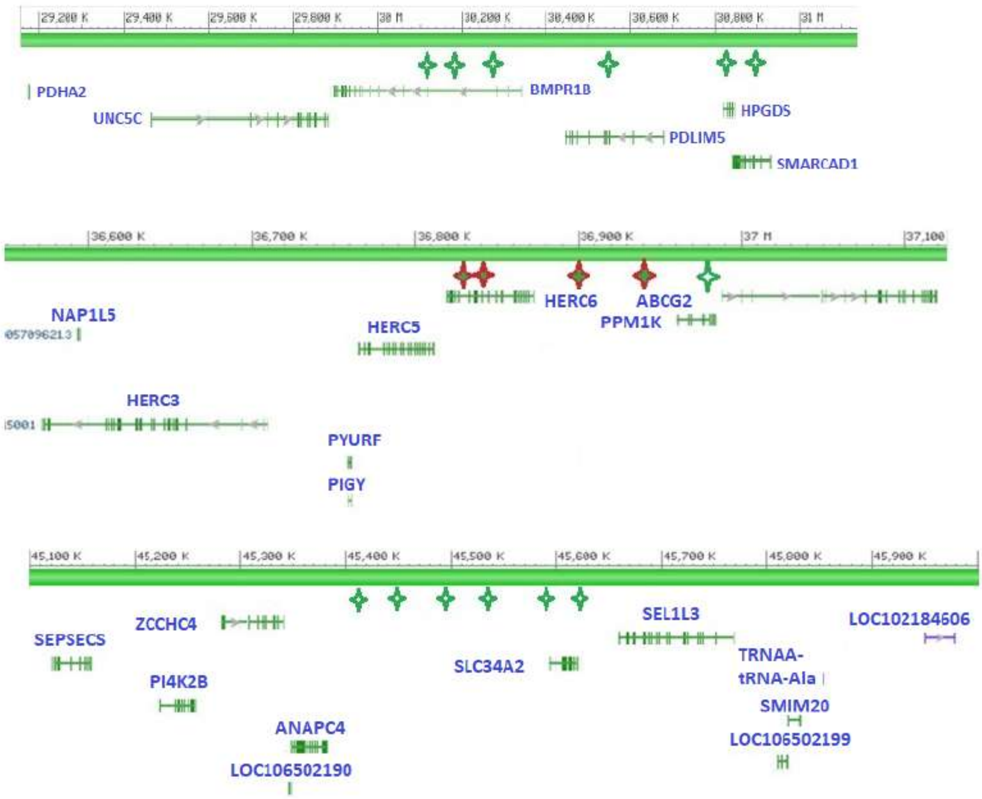

*Chr. 6*

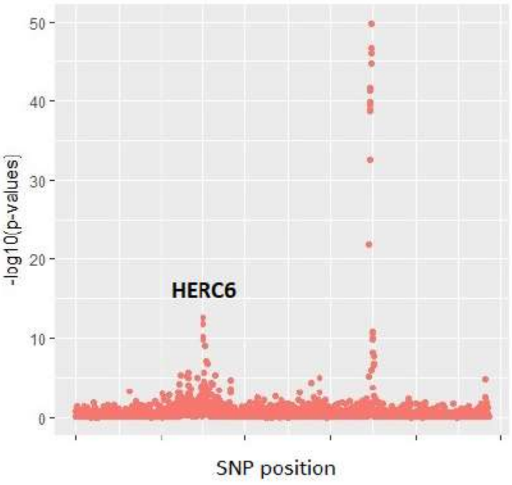

*Chr. 6*

**GOAT**

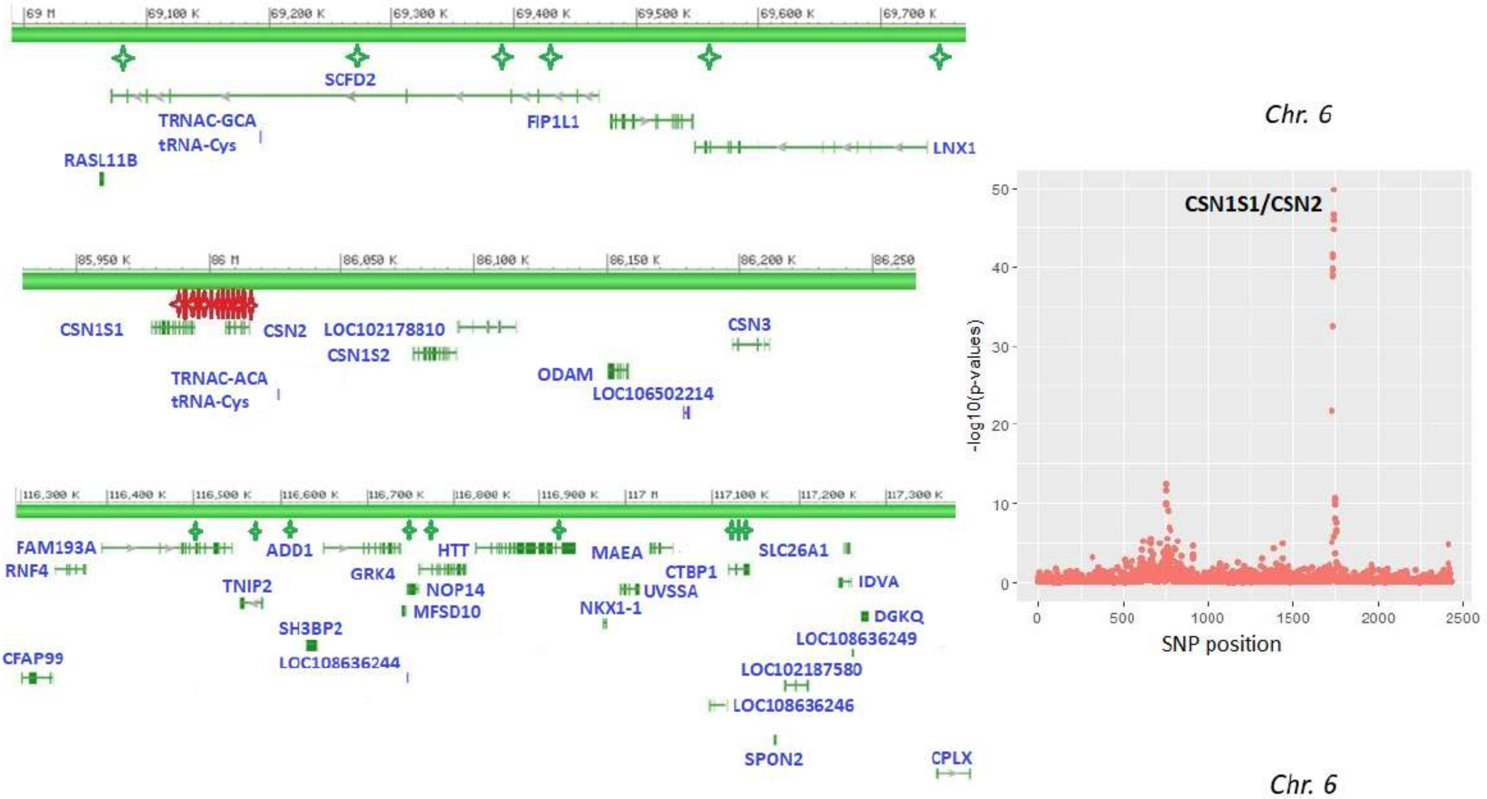
Chr. 6
CSN1S1/CSN2
-log10(p-values)
SNP position
Chr. 6
SCFD2
TRNAC-GCA
tRNA-Cys
FIP1L1
LNX1
RASL11B
CSN1S1
CSN2
LOC102178810
CSN1S2
CSN3
ODAM
LOC106502214
TRNAC-ACA
tRNA-Cys
FAM193A
RNF4
ADD1
GRK4
HTT
MAEA
SLC26A1
TNIP2
NOP14
MFSD10
CTBP1
UVSSA
IDVA
NKX1-1
DGKQ
SH3BP2
LOC108636249
LOC108636244
CFAP99
LOC102187580
LOC108636246
SPON2
CPLX

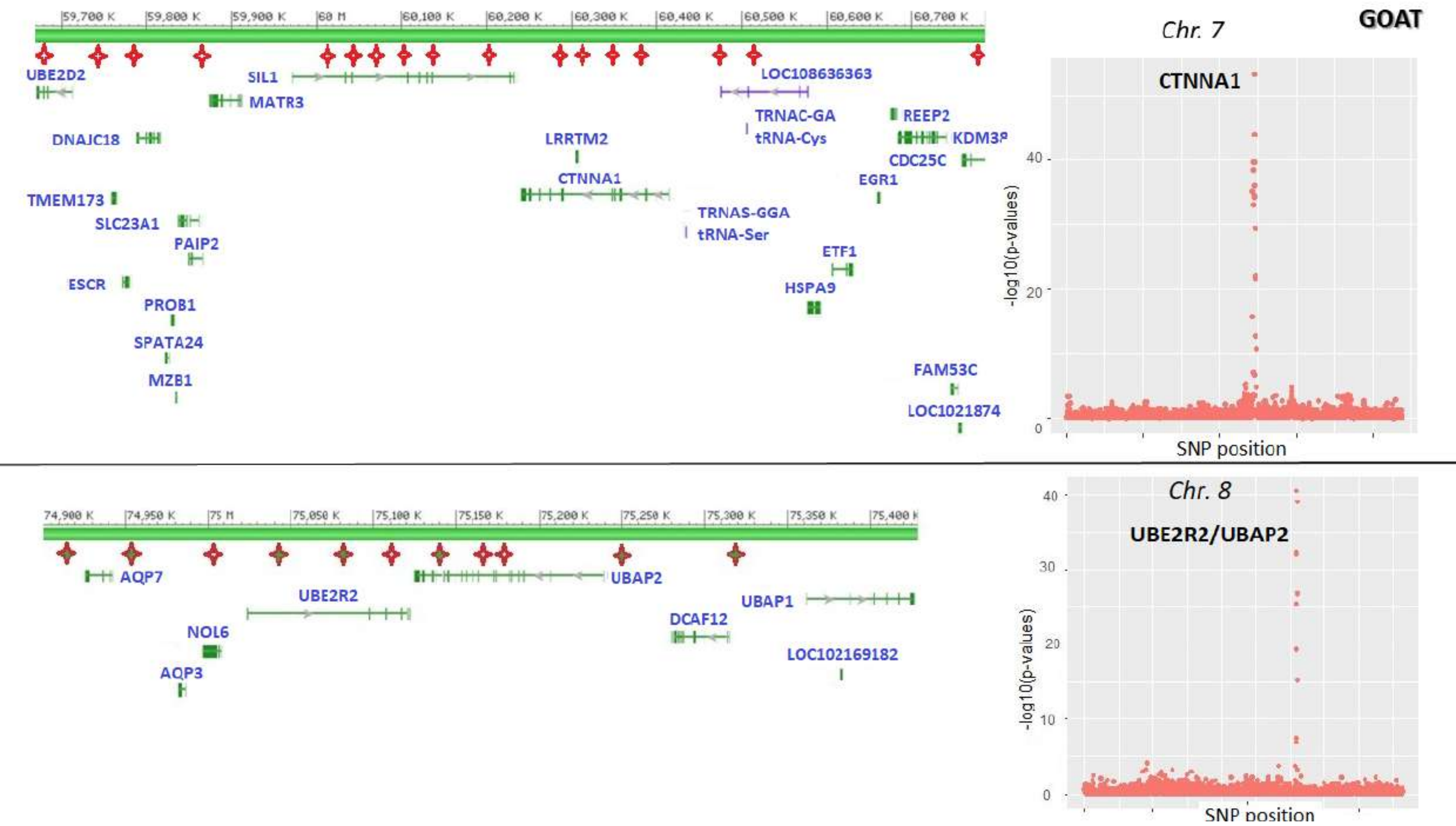
59,700 K
59,800 K
59,900 K
60 M
60,100 K
60,200 K
60,300 K
60,400 K
60,500 K
60,600 K
60,700 K
UBE2D2
SIL1
MATR3
LOC108636363
DNAJC18
TRNAC-GA
tRNA-Cys
REEP2
LRRTM2
CDC25C
CTNNA1
EGR1
TMEM173
TRNAS-GGA
SLC23A1
tRNA-Ser
PAIP2
ETF1
ESCR
HSPA9
PROB1
SPATA24
MZB1
FAM53C
LOC1021874
Chr. 7
GOAT
CTNNA1
-log10(p-values)
40
20
0
SNP position
74,900 K
74,950 K
75 M
75,050 K
75,100 K
75,150 K
75,200 K
75,250 K
75,300 K
75,350 K
75,400 K
AQP7
UBE2R2
UBAP2
UBAP1
NOL6
DCAF12
AQP3
LOC102169182
Chr. 8
UBE2R2/UBAP2
40
30
20
10
0
-log10(p-values)
SNP position

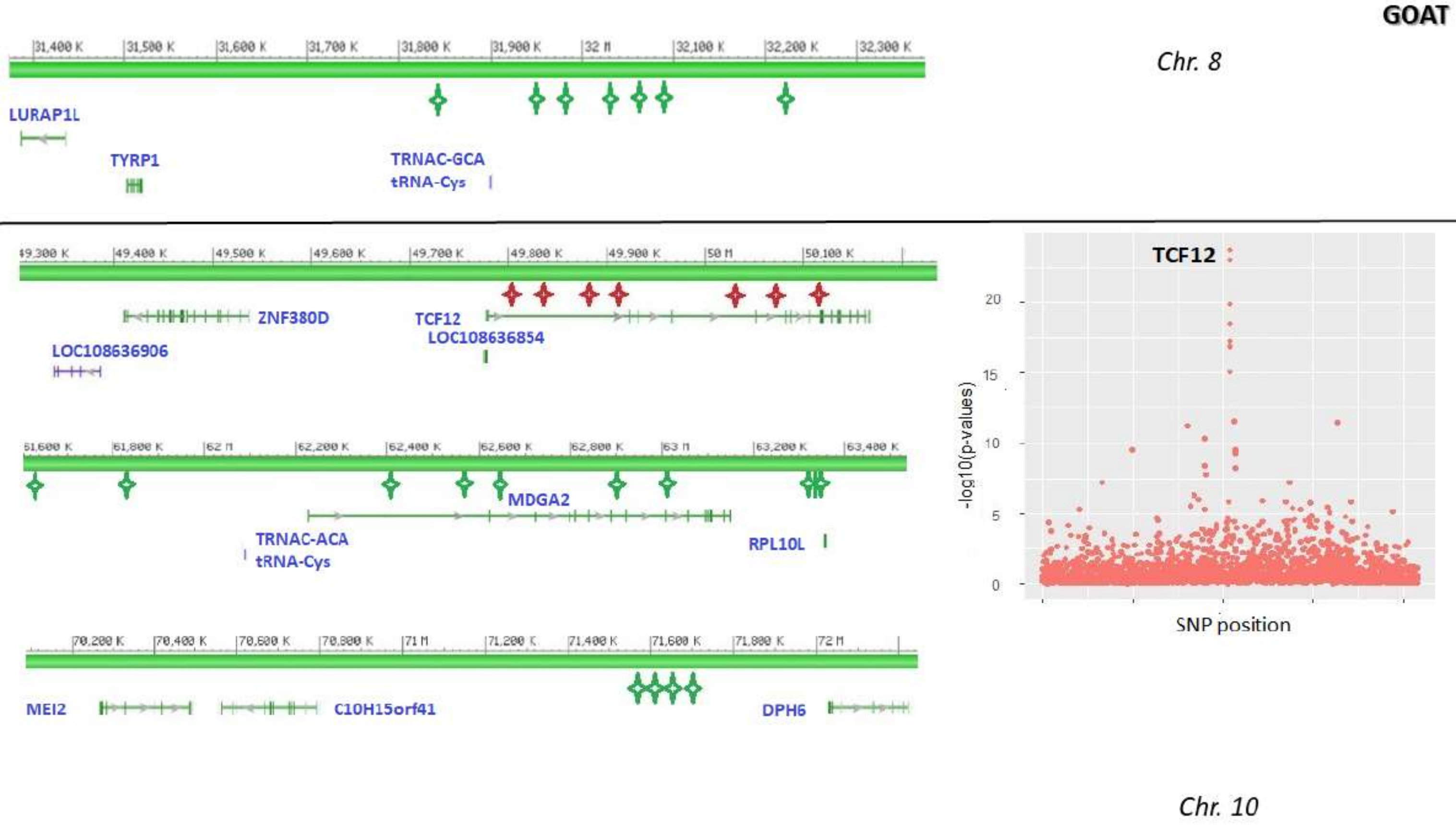

GOAT
Chr. 8
LURAP1L
TYRP1
TRNAC-GCA
tRNA-Cys
ZNF380D
TCF12
LOC108636854
LOC108636906
MDGA2
TRNAC-ACA
tRNA-Cys
RPL10L
MEI2
C10H15orf41
DPH6
TCF12
-log10(p-values)
SNP position
Chr. 10

**GOAT**

*Chr. 11*

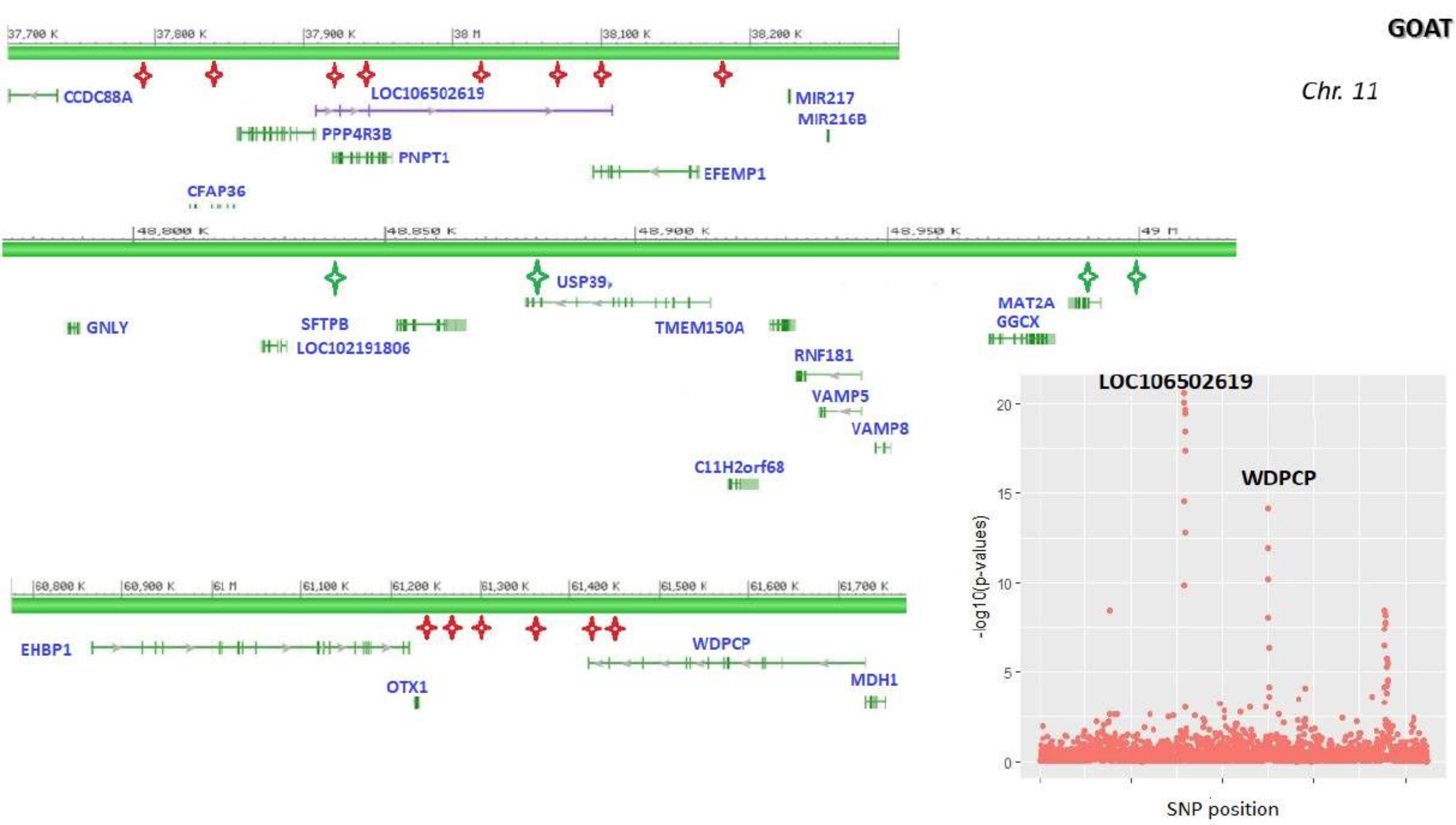
CCDC88A
LOC106502619
PPP4R3B
PNPT1
CFAP36
EFEMP1
MIR217
MIR216B
USP39
GNLY
SFTPB
LOC102191806
TMEM150A
MAT2A
GGCX
RNF181
VAMP5
VAMP8
C11H2orf68
EHBP1
OTX1
WDPCP
MDH1
LOC106502619
WDPCP
-log10(p-values)
SNP position

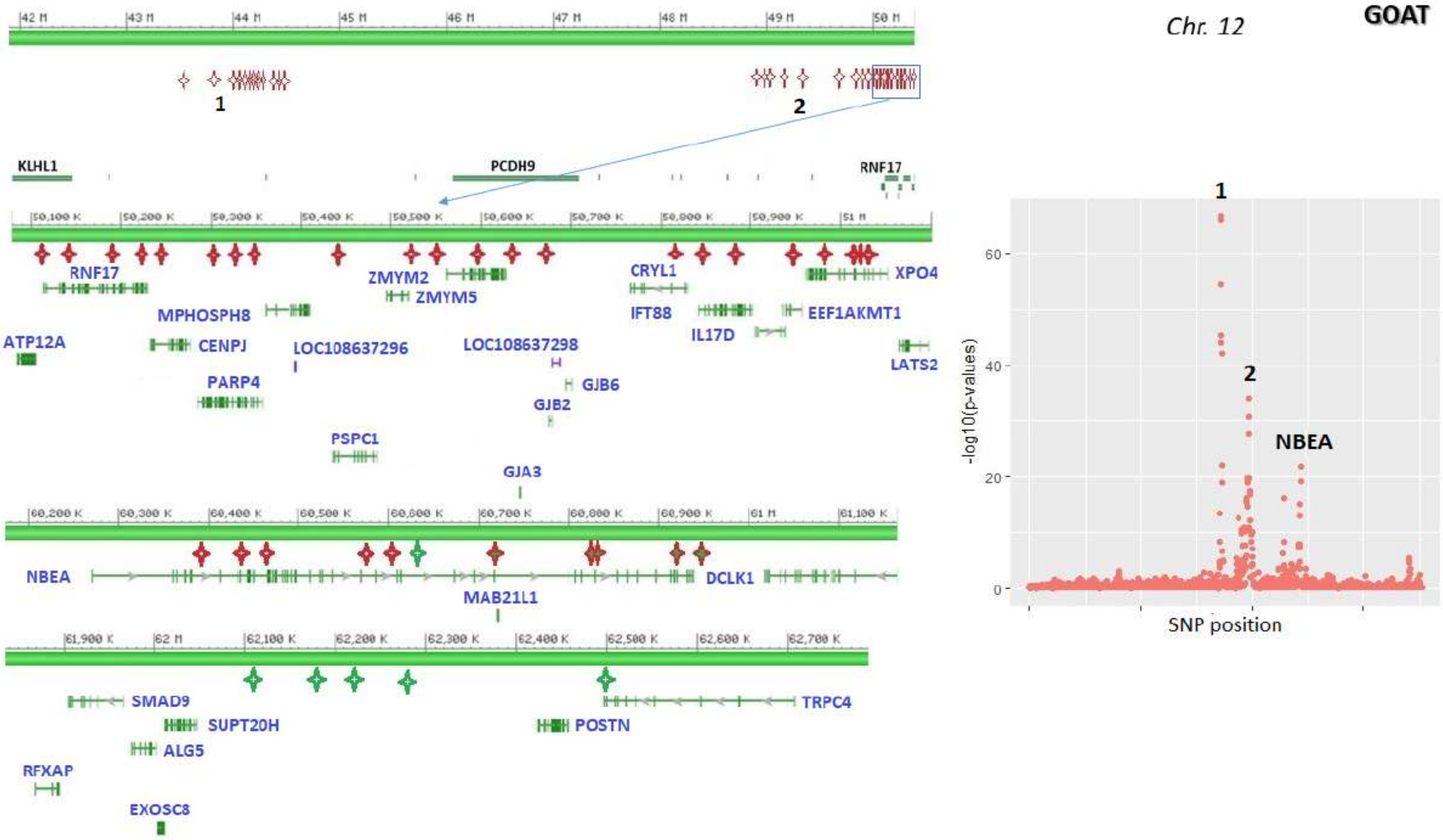
Chr. 12
GOAT
42 M
43 M
44 M
45 M
46 M
47 M
48 M
49 M
50 M
1
2
KLHL1
PCDH9
RNF17
RNF17
ZMYM2
ZMYM5
CRYL1
XPO4
MPHOSPH8
IFT88
IL17D
EEF1AKMT1
ATP12A
CENPJ
LOC108637296
LOC108637298
LATS2
PARP4
GJB6
GJB2
PSPC1
GJA3
NBEA
DCLK1
MAB21L1
SMAD9
TRPC4
SUPT20H
POSTN
ALG5
RFXAP
EXOSC8
-log10(p-values)
SNP position
NBEA
60
40
20
0

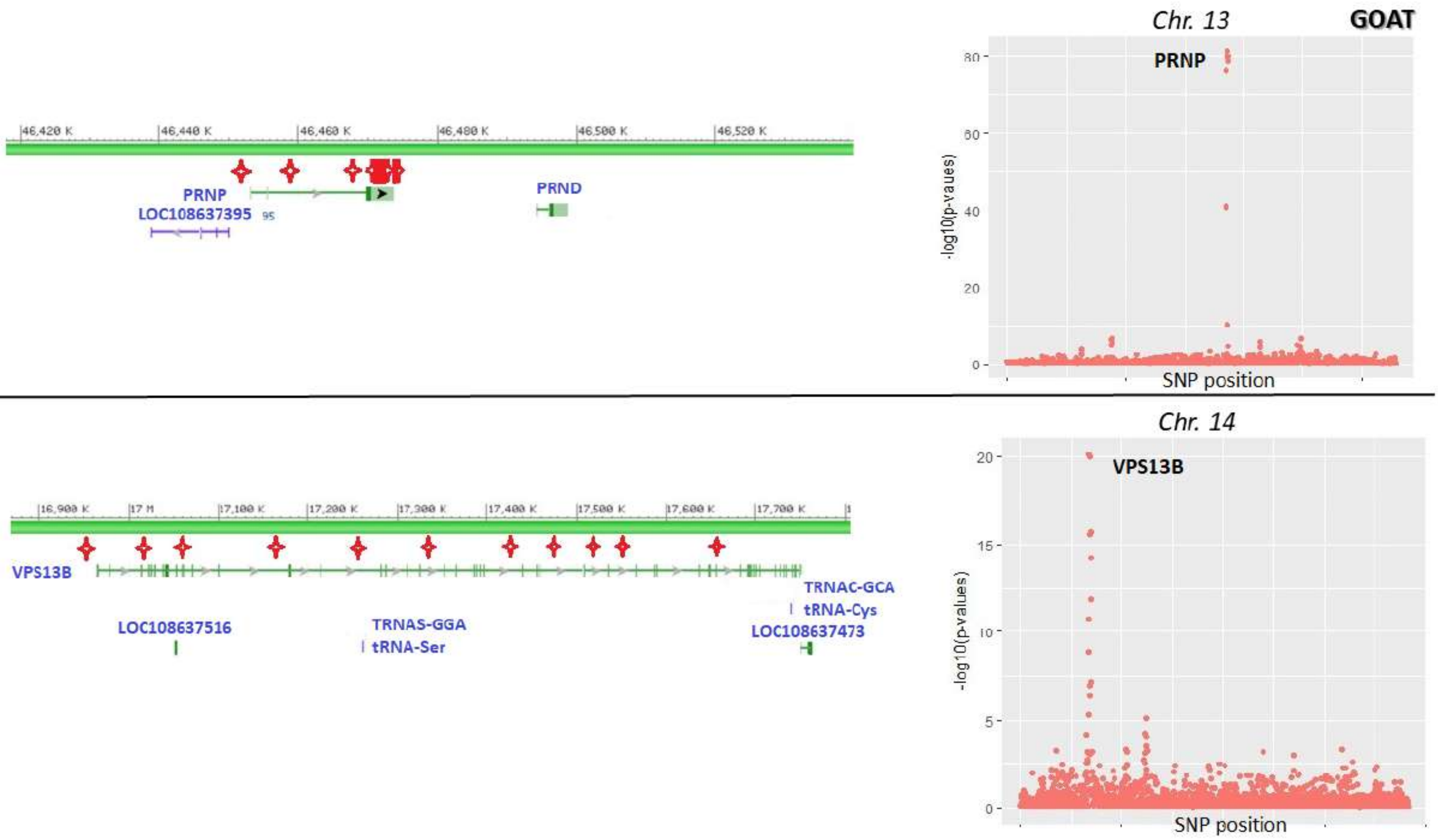

Chr. 13
GOAT
PRNP
-log10(p-vaues)
SNP position
46,420 K
46,440 K
46,460 K
46,480 K
46,500 K
46,520 K
PRNP
LOC108637395
PRND
Chr. 14
VPS13B
-log10(p-values)
SNP position
16,900 K
17 M
17,100 K
17,200 K
17,300 K
17,400 K
17,500 K
17,600 K
17,700 K
VPS13B
LOC108637516
TRNAS-GGA
tRNA-Ser
TRNAC-GCA
tRNA-Cys
LOC108637473

**GOAT**

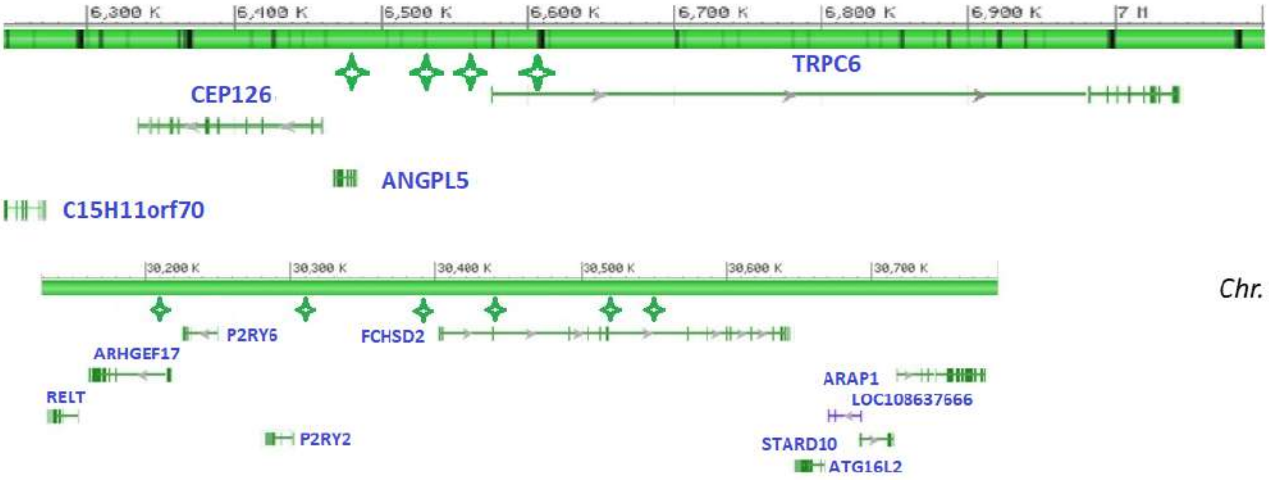

6,300 K
6,400 K
6,500 K
6,600 K
6,700 K
6,800 K
6,900 K
7 M
TRPC6
CEP126
ANGPL5
C15H11orf70
30,200 K
30,300 K
30,400 K
30,500 K
30,600 K
30,700 K
P2RY6
FCHSD2
ARHGEF17
RELT
P2RY2
ARAP1
LOC108637666
STARD10
ATG16L2

*Chr. 15*

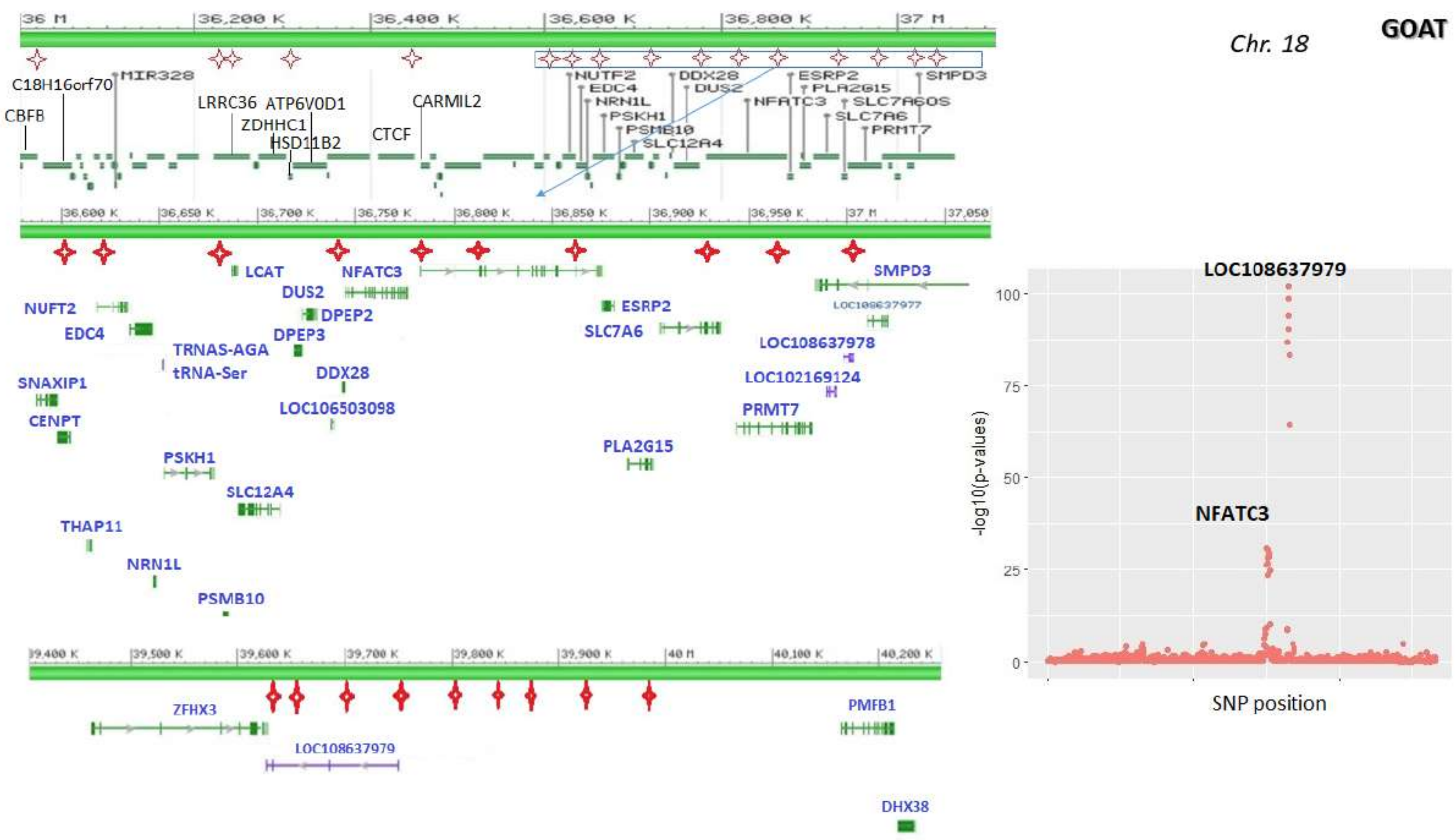
Chr. 18
GOAT
36 M
36,200 K
36,400 K
36,600 K
36,800 K
37 M
C18H16orf70
MIR328
CBFB
LRRC36
ATP6V0D1
ZDHHC1
HSD11B2
CARMIL2
CTCF
NUTF2
EDC4
NRN1L
PSKH1
PSMB10
SLC12A4
DDX28
DUS2
NFATC3
ESRP2
PLA2G15
SLC7A6OS
SLC7A6
PRMT7
SMPD3
LCAT
DPEP2
DPEP3
NUFT2
TRNAS-AGA
tRNA-Ser
SNAXIP1
CENPT
LOC106503098
LOC108637977
LOC108637978
LOC102169124
THAP11
ZFHX3
LOC108637979
PMFB1
DHX38
-log10(p-values)
SNP position
100
75
50
25
0

**GOAT**

*Chr. 19*

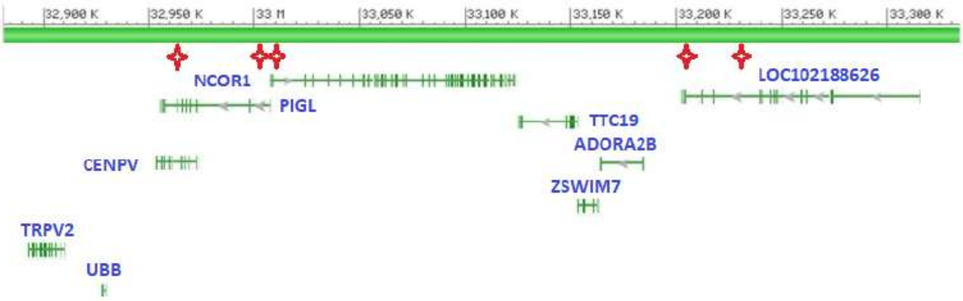

32,900 K
32,950 K
33 M
33,050 K
33,100 K
33,150 K
33,200 K
33,250 K
33,300 K
NCOR1
PIGL
LOC102188626
TTC19
ADORA2B
CENPV
ZSWIM7
TRPV2
UBB

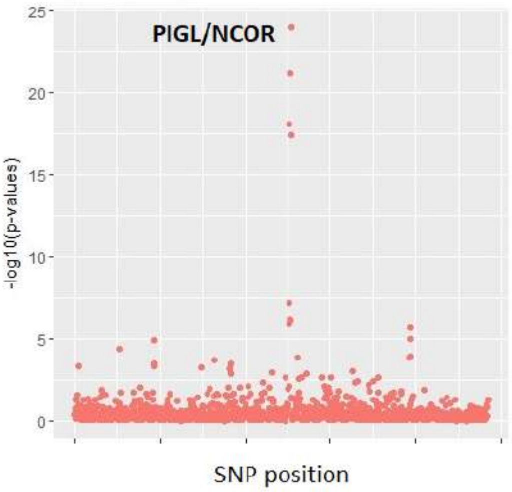

PIGL/NCOR
-log10(p-values)
25
20
15
10
5
0
SNP position

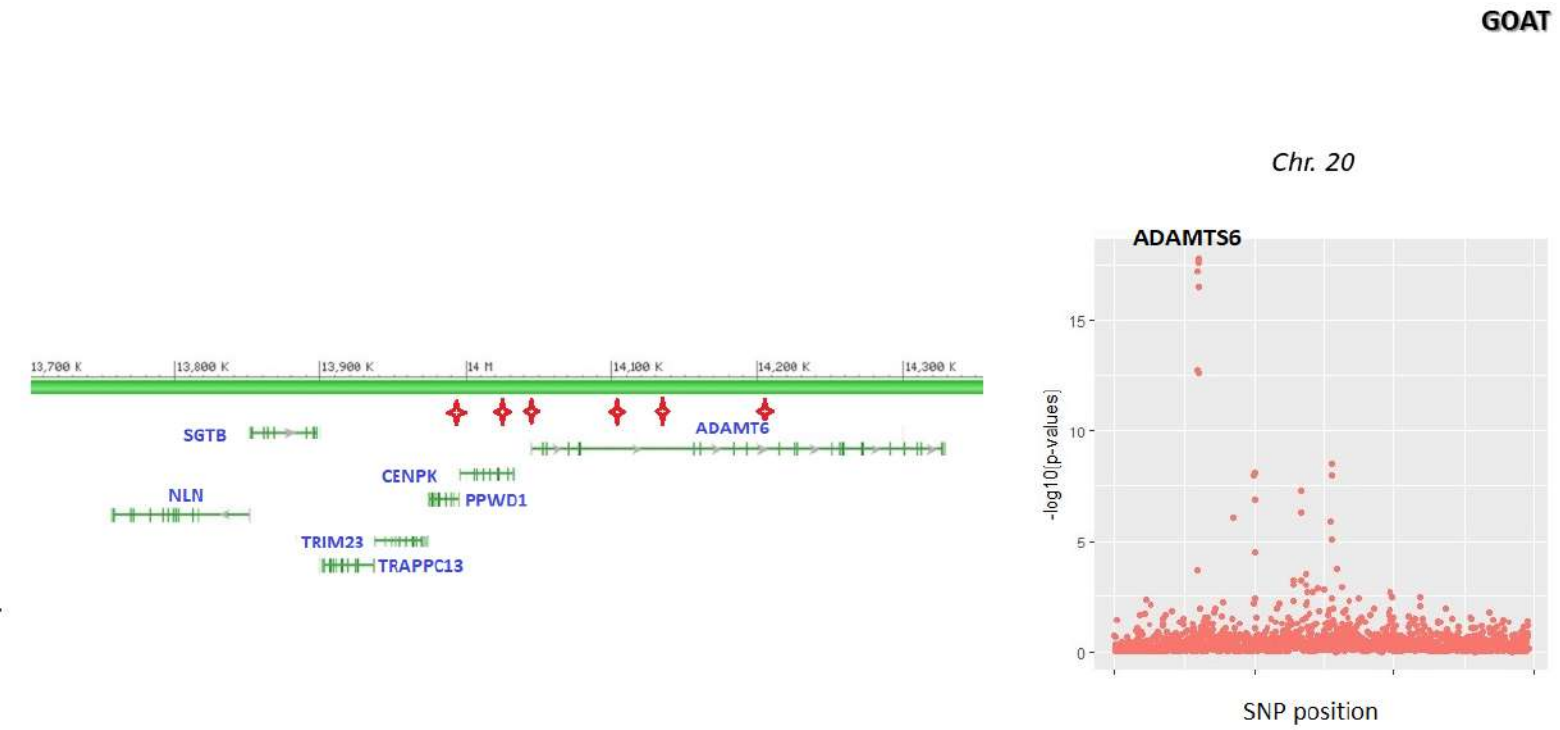
GOAT
Chr. 20
ADAMTS6
13,700 K
13,800 K
13,900 K
14 M
14,100 K
14,200 K
14,300 K
SGTB
ADAMT6
CENPK
PPWD1
NLN
TRIM23
TRAPPC13
-log10(p-values)
15
10
5
0
SNP position

**GOAT**

*Chr. 22*

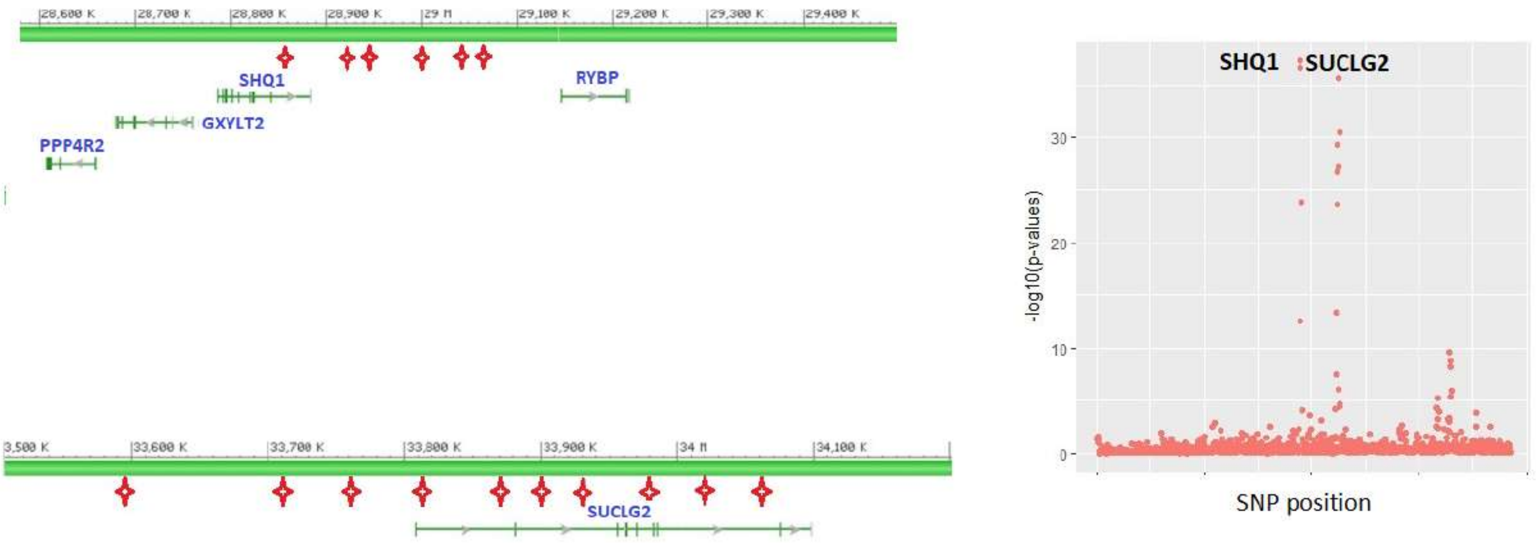

28,600 K
28,700 K
28,800 K
28,900 K
29 M
29,100 K
29,200 K
29,300 K
29,400 K
SHQ1
RYBP
GXYLT2
PPP4R2
33,600 K
33,700 K
33,800 K
33,900 K
34 M
34,100 K
SUCLG2
SHQ1
SUCLG2
-log10(p-values)
30
20
10
0
SNP position

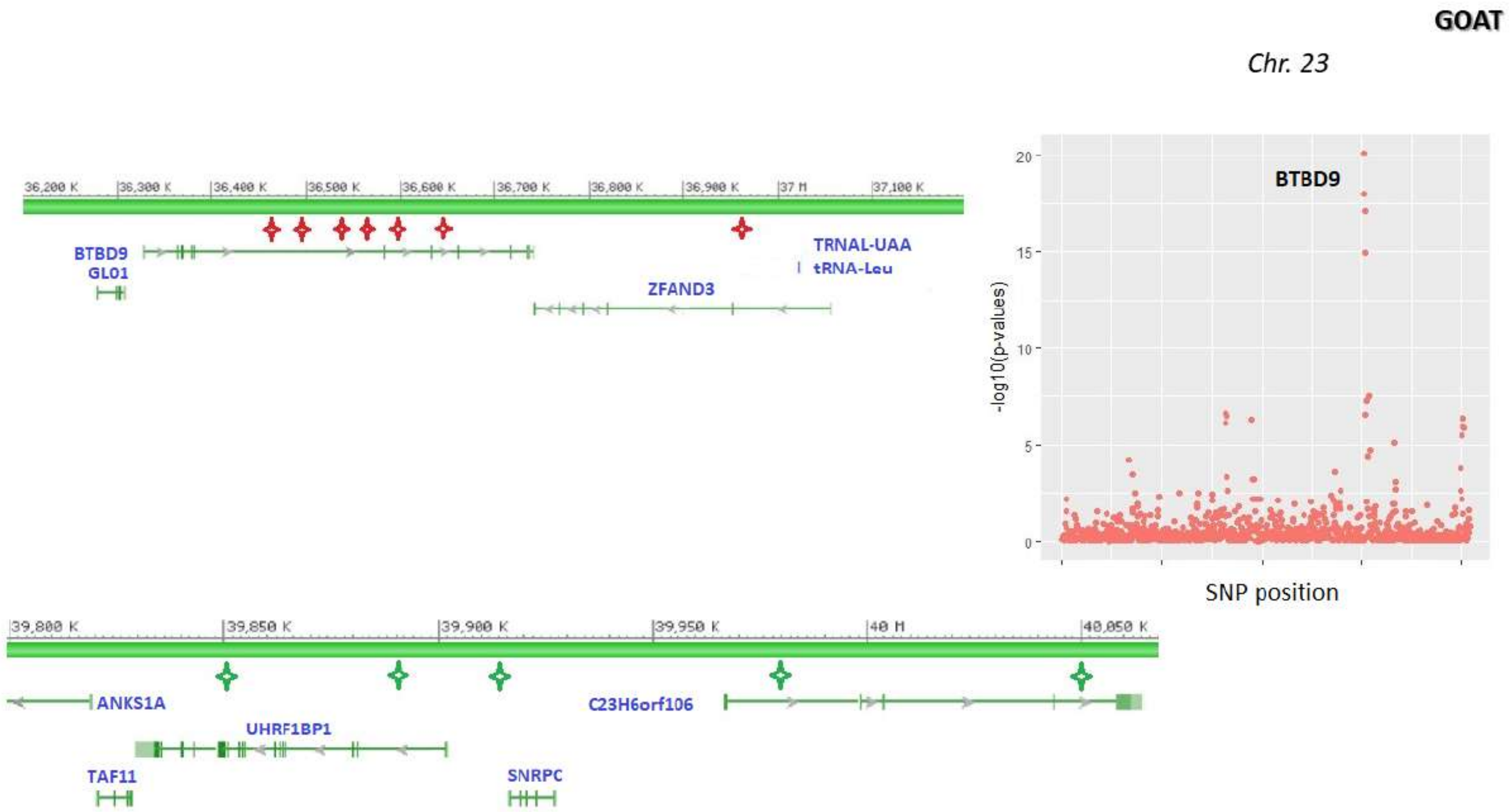
GOAT
Chr. 23
BTBD9
-log10(p-values)
SNP position
36,200 K
36,300 K
36,400 K
36,500 K
36,600 K
36,700 K
36,800 K
36,900 K
37 M
37,100 K
BTBD9
GLO1
TRNAL-UAA
tRNA-Leu
ZFAND3
39,800 K
39,850 K
39,900 K
39,950 K
40 M
40,050 K
ANKS1A
C23H6orf106
UHRF1BP1
TAF11
SNRPC

**GOAT**

*Chr. 25*

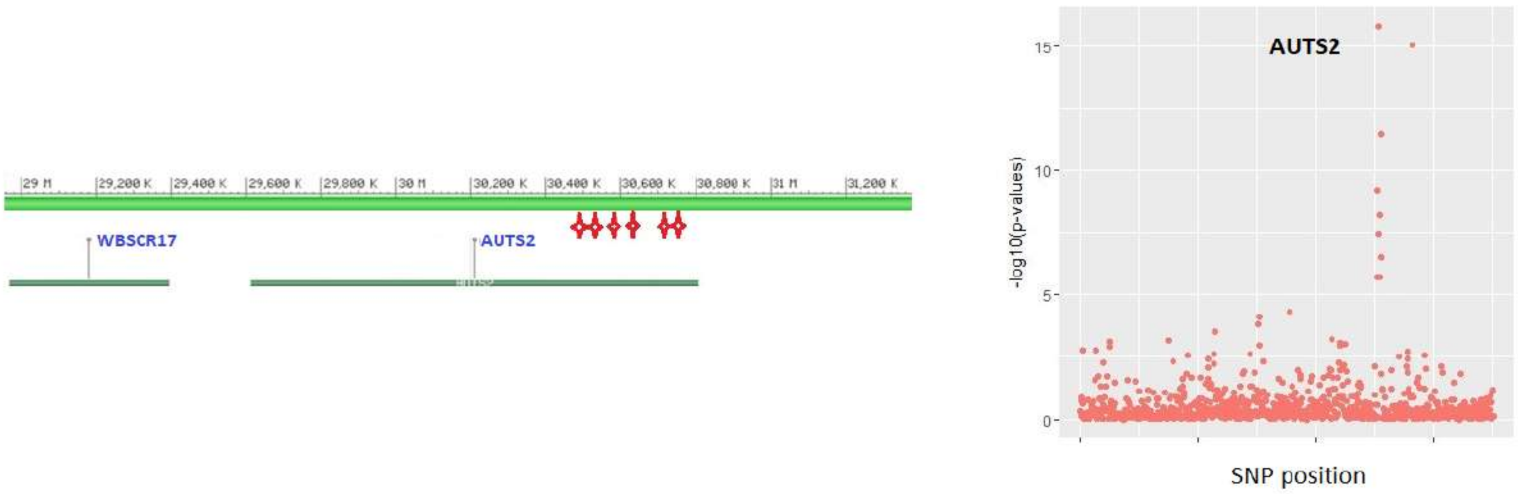

29 M
29,200 K
29,400 K
29,600 K
29,800 K
30 M
30,200 K
30,400 K
30,600 K
30,800 K
31 M
31,200 K
WBSCR17
AUTS2
AUTS2
-log10(p-values)
0
5
10
15
SNP position

**GOAT**

*Chr. 26*

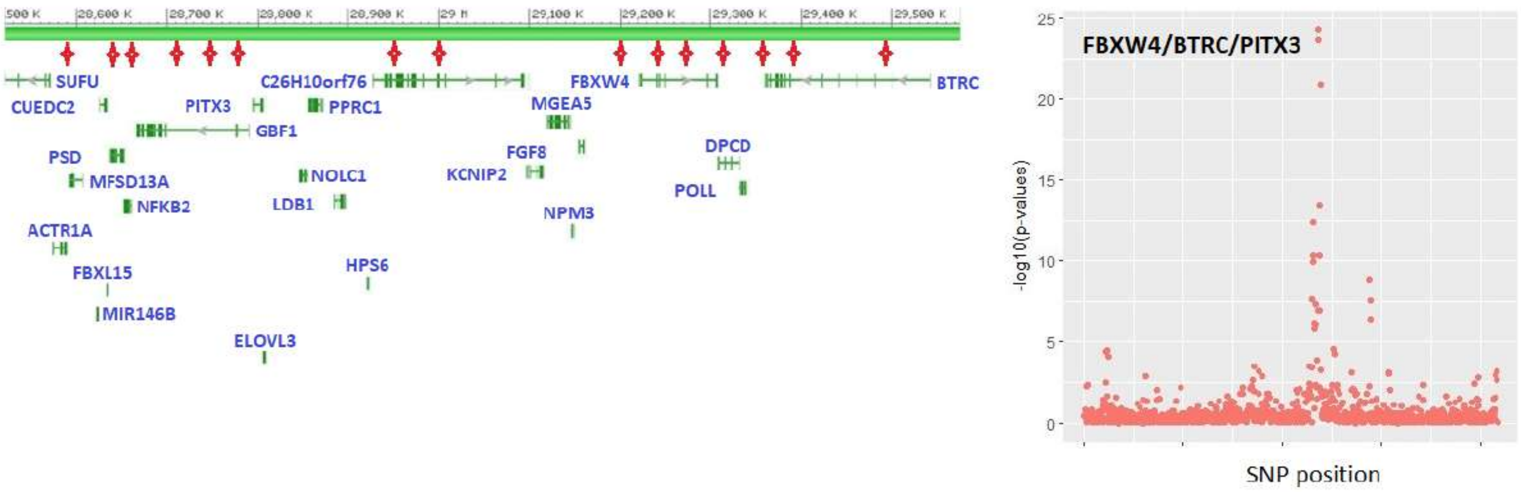
SUFU
CUEDC2
PITX3
GBF1
C26H10orf76
PPRC1
PSD
MFSD13A
NFKB2
ACTR1A
FBXL15
MIR146B
ELOVL3
NOLC1
LDB1
HPS6
FBXW4
MGEA5
FGF8
KCNIP2
NPM3
DPCD
POLL
BTRC
FBXW4/BTRC/PITX3
-log10(p-values)
SNP position
0
5
10
15
20
25

SHEEP

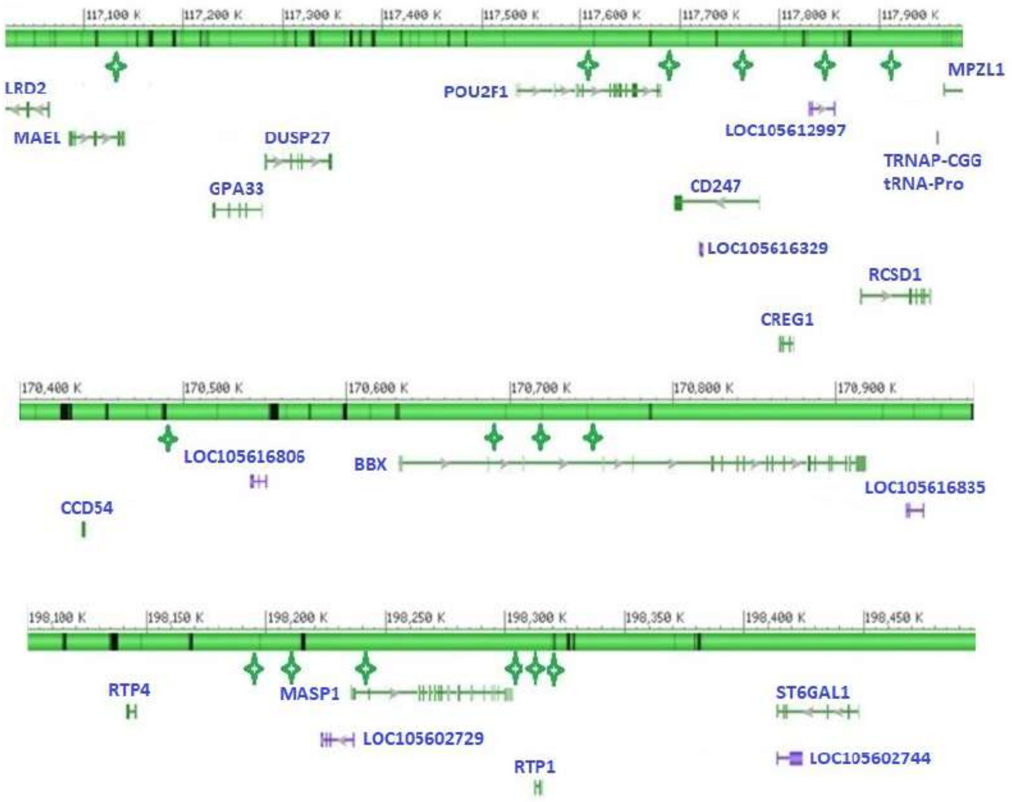
117,100 K
117,200 K
117,300 K
117,400 K
117,500 K
117,600 K
117,700 K
117,800 K
117,900 K
LRD2
MAEL
DUSP27
GPA33
POU2F1
LOC105612997
MPZL1
TRNAP-CGG
tRNA-Pro
CD247
LOC105616329
RCSD1
CREG1
170,400 K
170,500 K
170,600 K
170,700 K
170,800 K
170,900 K
LOC105616806
BBX
LOC105616835
CCD54
198,100 K
198,150 K
198,200 K
198,250 K
198,300 K
198,350 K
198,400 K
198,450 K
RTP4
MASP1
ST6GAL1
LOC105602729
LOC105602744
RTP1

*Chr. 1*

**SHEEP**

*Chr. 2*

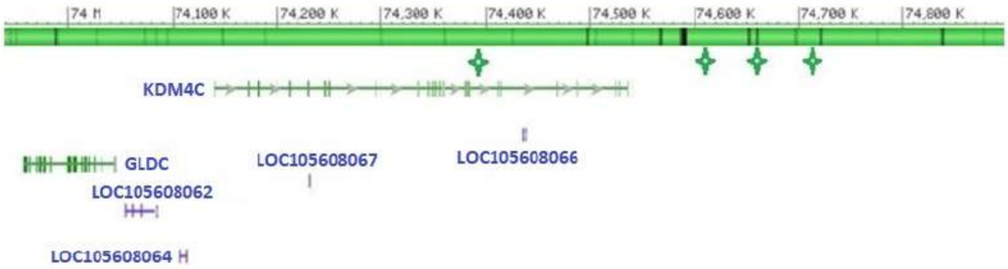
74 M
74,100 K
74,200 K
74,300 K
74,400 K
74,500 K
74,600 K
74,700 K
74,800 K
KDM4C
GLDC
LOC105608067
LOC105608066
LOC105608062
LOC105608064

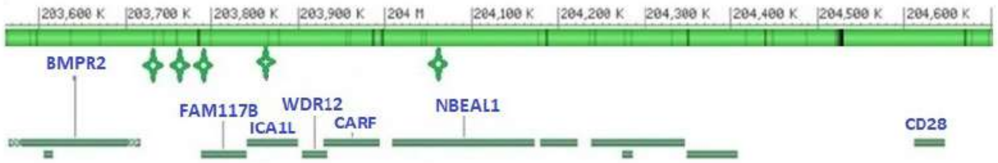
203,600 K
203,700 K
203,800 K
203,900 K
204 M
204,100 K
204,200 K
204,300 K
204,400 K
204,500 K
204,600 K
BMPR2
FAM117B
WDR12
ICA1L
CARF
NBEAL1
CD28

**SHEEP**

*Chr. 3*

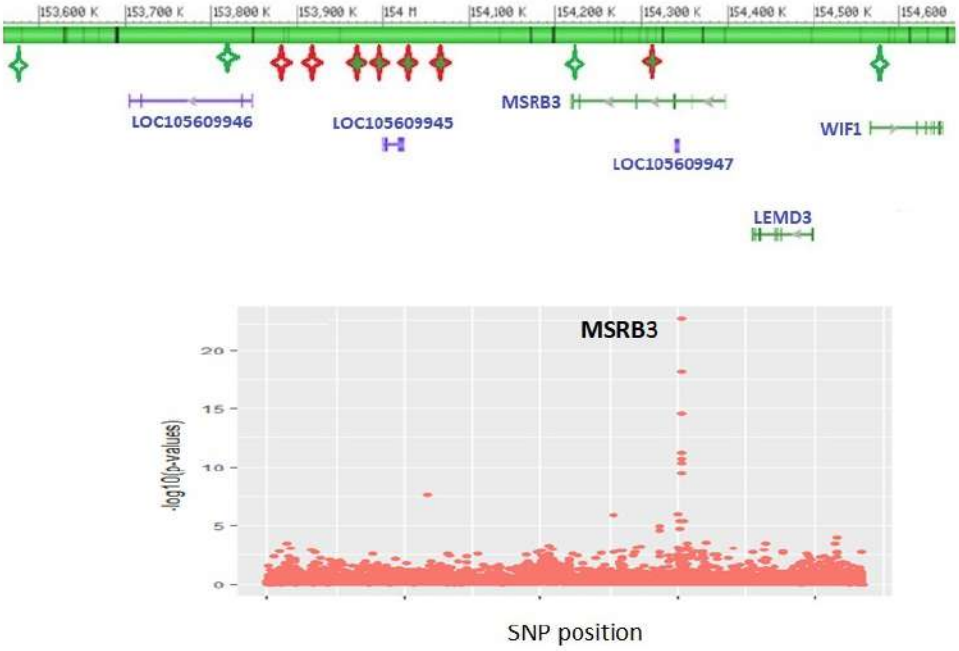

153,600 K
153,700 K
153,800 K
153,900 K
154 M
154,100 K
154,200 K
154,300 K
154,400 K
154,500 K
154,600
LOC105609946
LOC105609945
MSRB3
LOC105609947
WIF1
LEMD3
MSRB3
20
15
10
5
0
-log10(p-values)
SNP position

**SHEEP**

*Chr. 4*

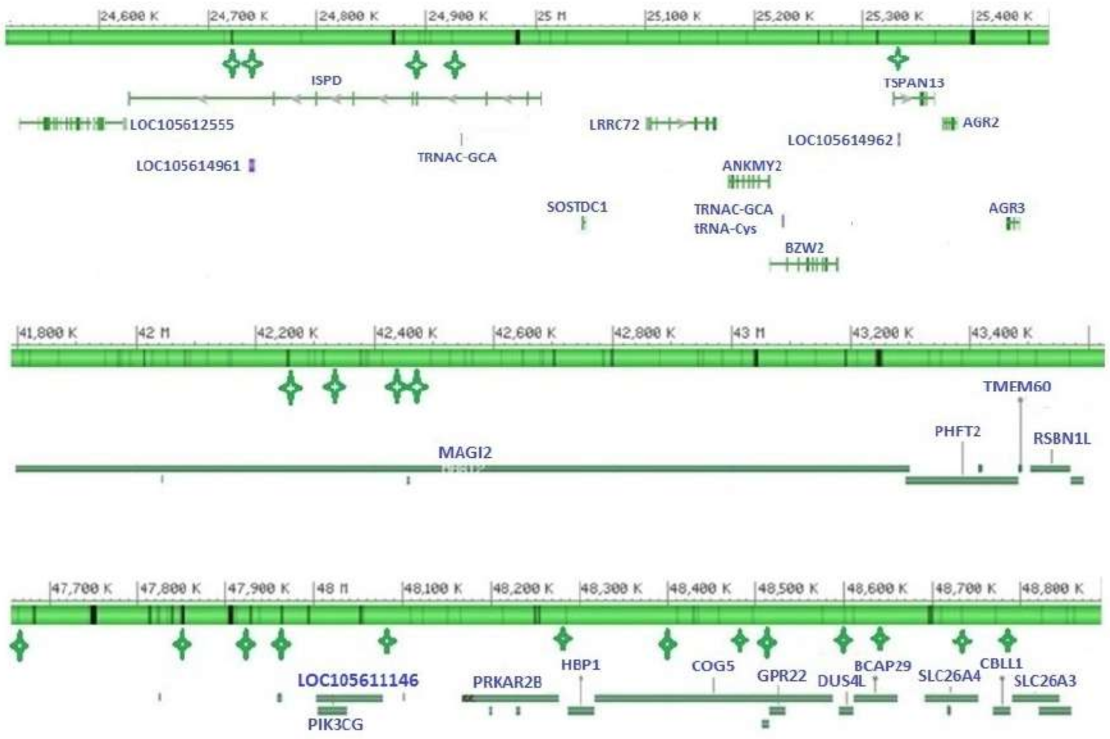
24,600 K
24,700 K
24,800 K
24,900 K
25 M
25,100 K
25,200 K
25,300 K
25,400 K
ISPD
TSPAN13
LOC105612555
LRRC72
AGR2
LOC105614962
TRNAC-GCA
LOC105614961
ANKMY2
SOSTDC1
TRNAC-GCA
tRNA-Cys
AGR3
BZW2
41,800 K
42 M
42,200 K
42,400 K
42,600 K
42,800 K
43 M
43,200 K
43,400 K
TMEM60
PHFT2
RSBN1L
MAGI2
47,700 K
47,800 K
47,900 K
48 M
48,100 K
48,200 K
48,300 K
48,400 K
48,500 K
48,600 K
48,700 K
48,800 K
LOC105611146
PRKAR2B
HBP1
COG5
GPR22
DUS4L
BCAP29
SLC26A4
CBLL1
SLC26A3
PIK3CG

**SHEEP**

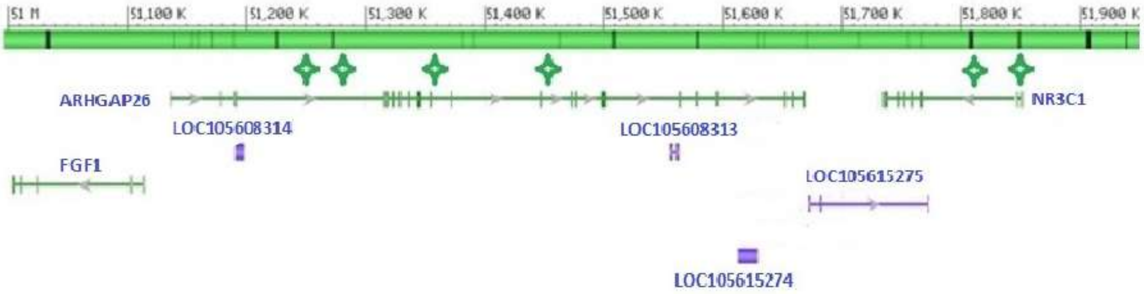

51 M
51,100 K
51,200 K
51,300 K
51,400 K
51,500 K
51,600 K
51,700 K
51,800 K
51,900 K
ARHGAP26
LOC105608314
LOC105608313
NR3C1
FGF1
LOC105615275
LOC105615274

*Chr. 5*

**SHEEP**

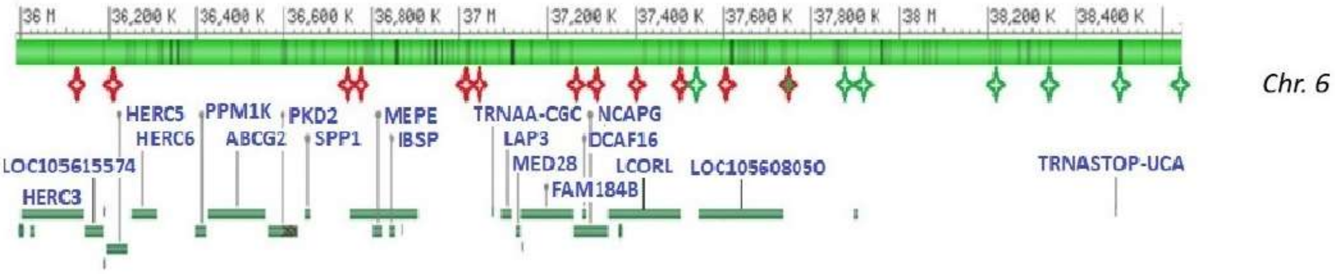
36 M
36,200 K
36,400 K
36,600 K
36,800 K
37 M
37,200 K
37,400 K
37,600 K
37,800 K
38 M
38,200 K
38,400 K
Chr. 6
HERC5
PPM1K
PKD2
MEPE
TRNAA-CGC
NCAPG
HERC6
ABCG2
SPP1
IBSP
LAP3
DCAF16
LOC105615574
MED28
LCORL
LOC105608050
TRNASTOP-UCA
HERC3
FAM184B

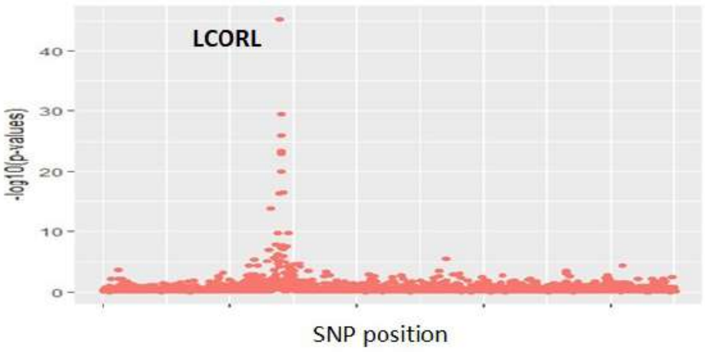
LCORL
40
30
20
10
0
-log10(p-values)
SNP position

**SHEEP**

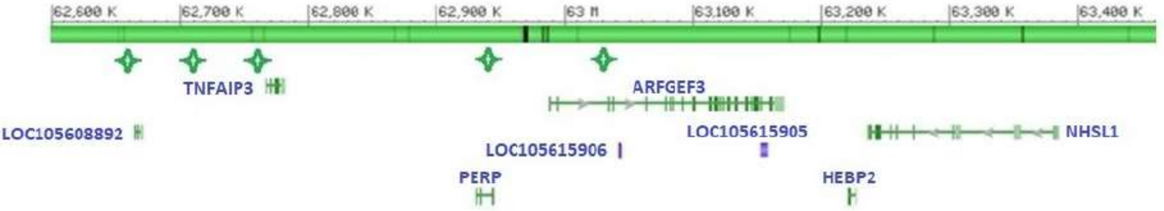

*Chr. 8*

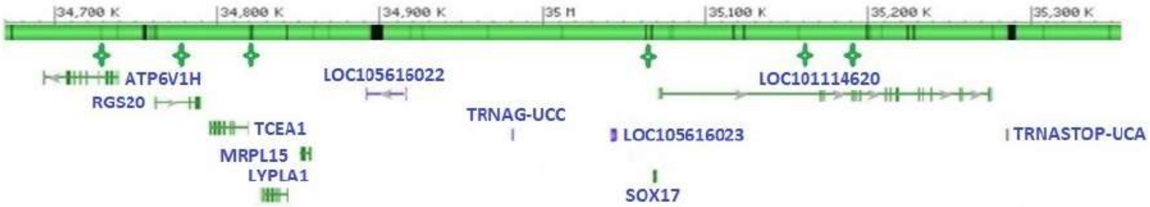

*Chr. 9*

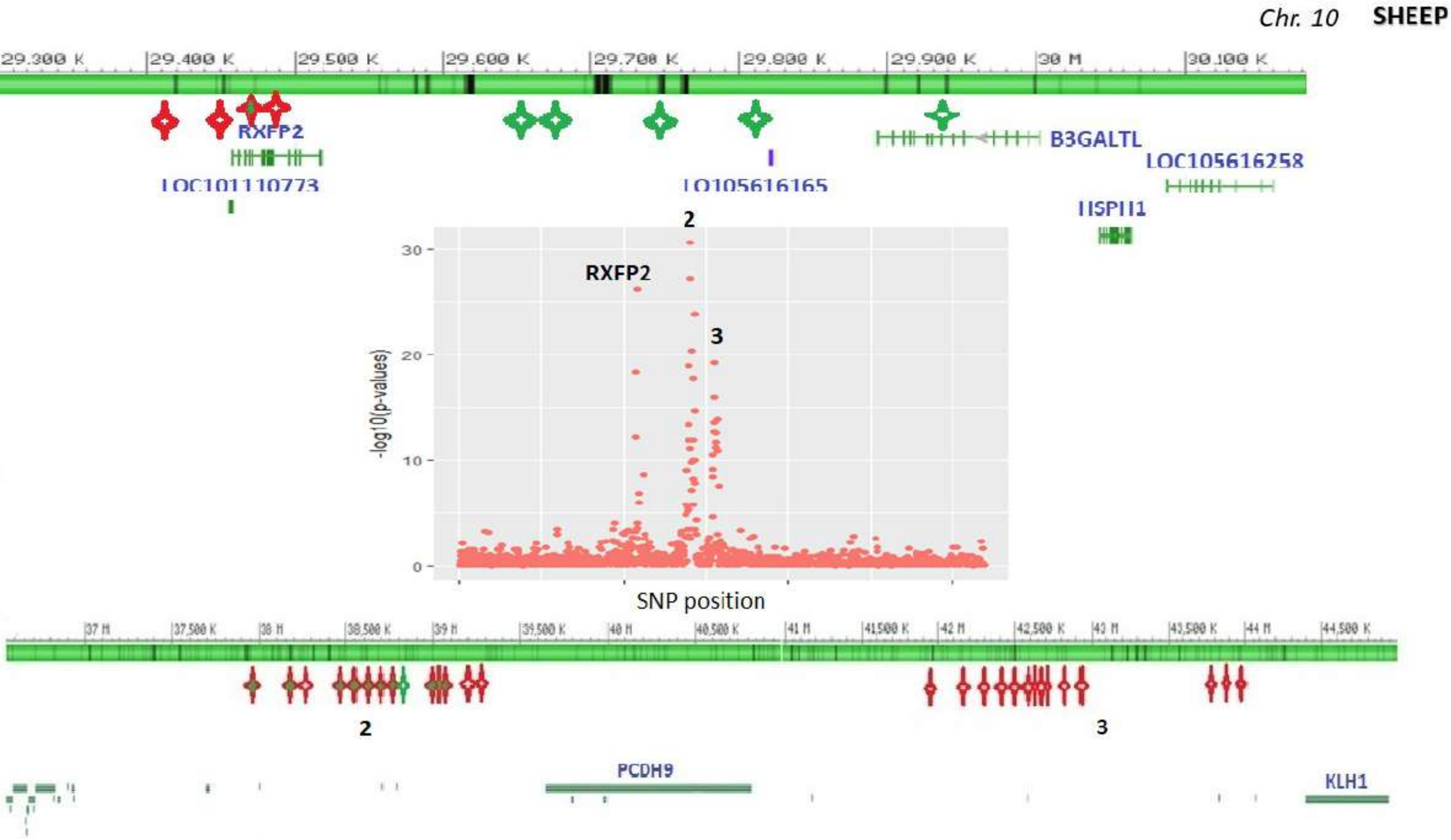
Chr. 10 SHEEP
29.300 K
29.400 K
29.500 K
29.600 K
29.700 K
29.800 K
29.900 K
30 M
30.100 K
RXFP2
LOC101110773
LO105616165
B3GALTL
LOC105616258
HSPH1
2
RXFP2
3
-log10(p-values)
30
20
10
0
SNP position
37 M
37,500 K
38 M
38,500 K
39 M
39,500 K
40 M
40,500 K
41 M
41500 K
42 M
42,500 K
43 M
43,500 K
44 M
44,500 K
2
3
PCDH9
KLH1

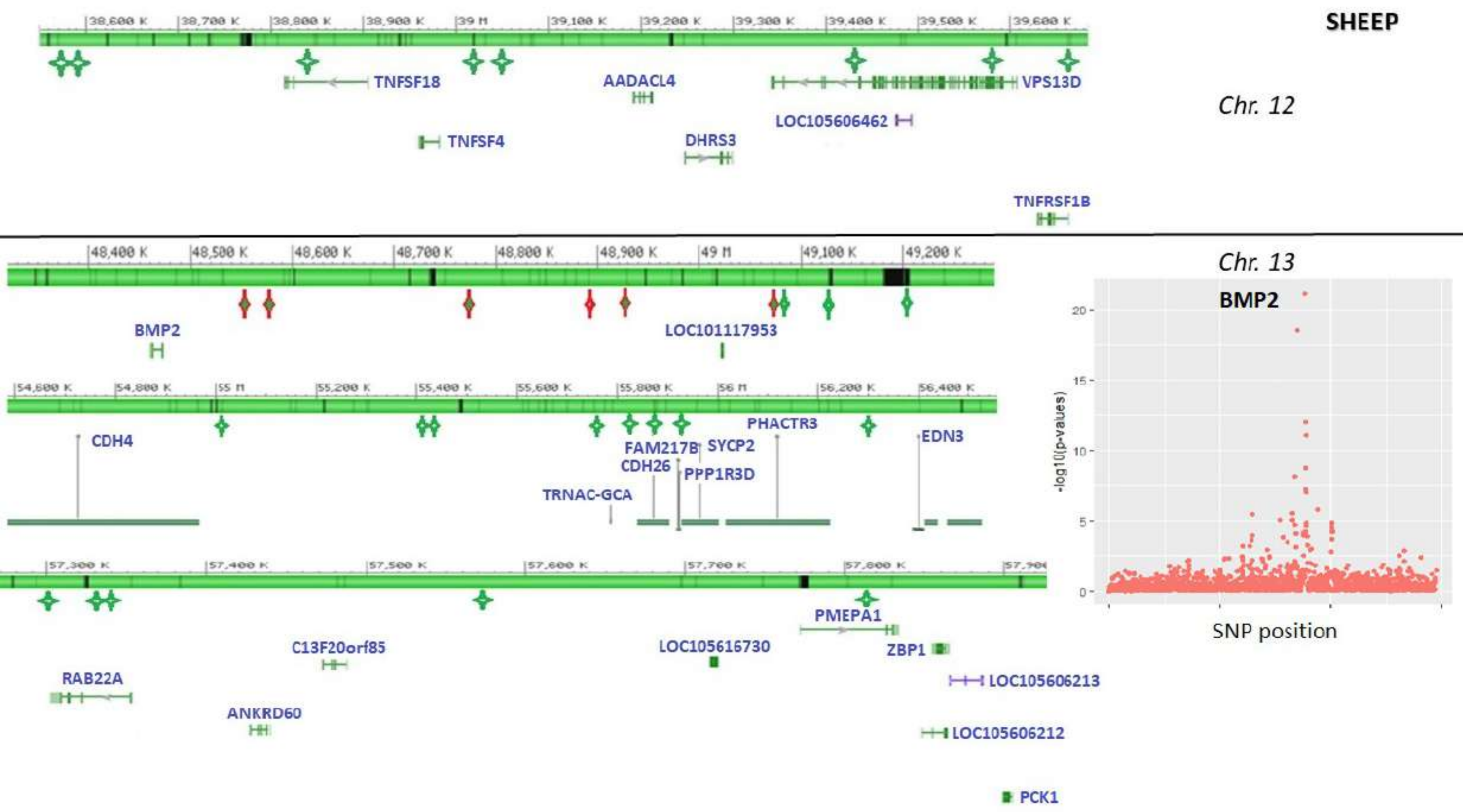
SHEEP
Chr. 12
TNFSF18
AADACL4
VPS13D
LOC105606462
TNFSF4
DHRS3
TNFRSF1B
Chr. 13
BMP2
LOC101117953
CDH4
FAM217B
SYCP2
PHACTR3
EDN3
CDH26
PPP1R3D
TRNAC-GCA
PMEPA1
C13F20orf85
LOC105616730
ZBP1
RAB22A
LOC105606213
ANKRD60
LOC105606212
PCK1
-log10(p-values)
SNP position

**SHEEP**

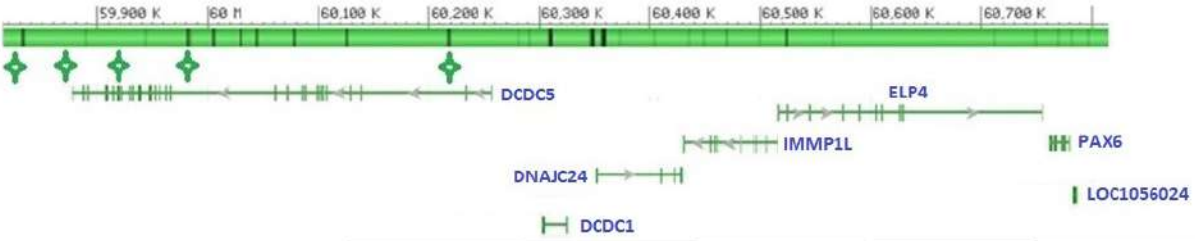

*Chr. 15*

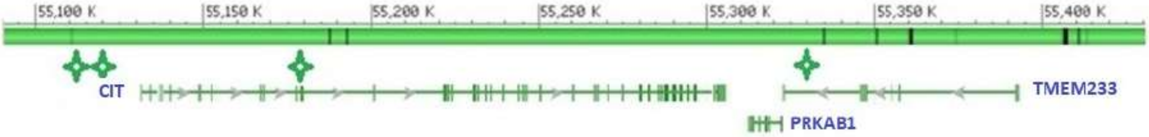

*Chr. 17*

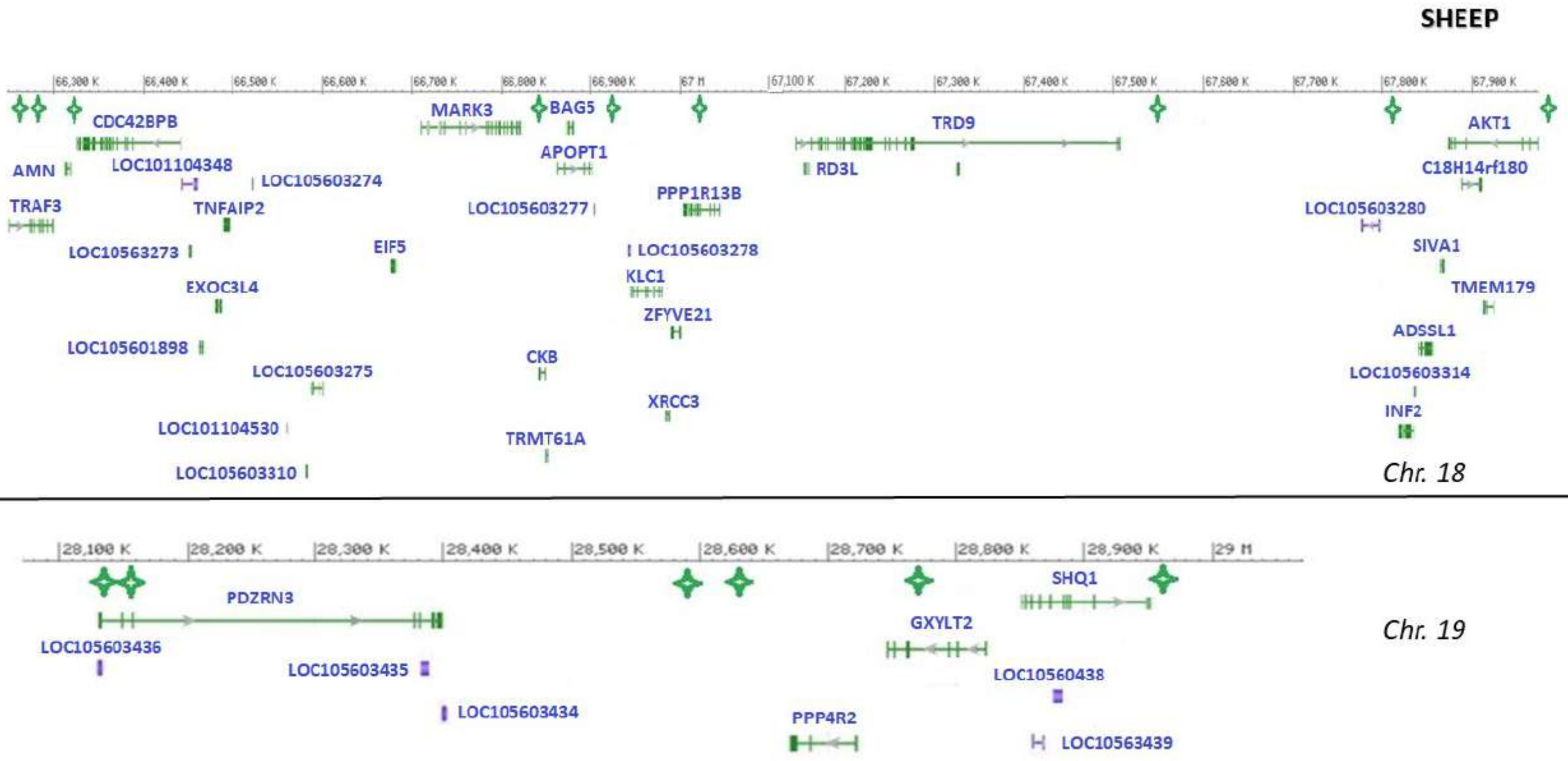
SHEEP
CDC42BPB
AMN
LOC101104348
LOC105603274
TRAF3
TNFAIP2
LOC10563273
EIF5
EXOC3L4
LOC105601898
LOC105603275
LOC101104530
LOC105603310
MARK3
BAG5
APOPT1
LOC105603277
CKB
TRMT61A
PPP1R13B
LOC105603278
KLC1
ZFYVE21
XRCC3
TRD9
RD3L
AKT1
C18H14rf180
LOC105603280
SIVA1
TMEM179
ADSSL1
LOC105603314
INF2
Chr. 18
28,100 K
28,200 K
28,300 K
28,400 K
28,500 K
28,600 K
28,700 K
28,800 K
28,900 K
29 M
PDZRN3
LOC105603436
LOC105603435
LOC105603434
PPP4R2
GXYLT2
SHQ1
LOC10560438
LOC10563439
Chr. 19

**SHEEP**

*Chr. 21*

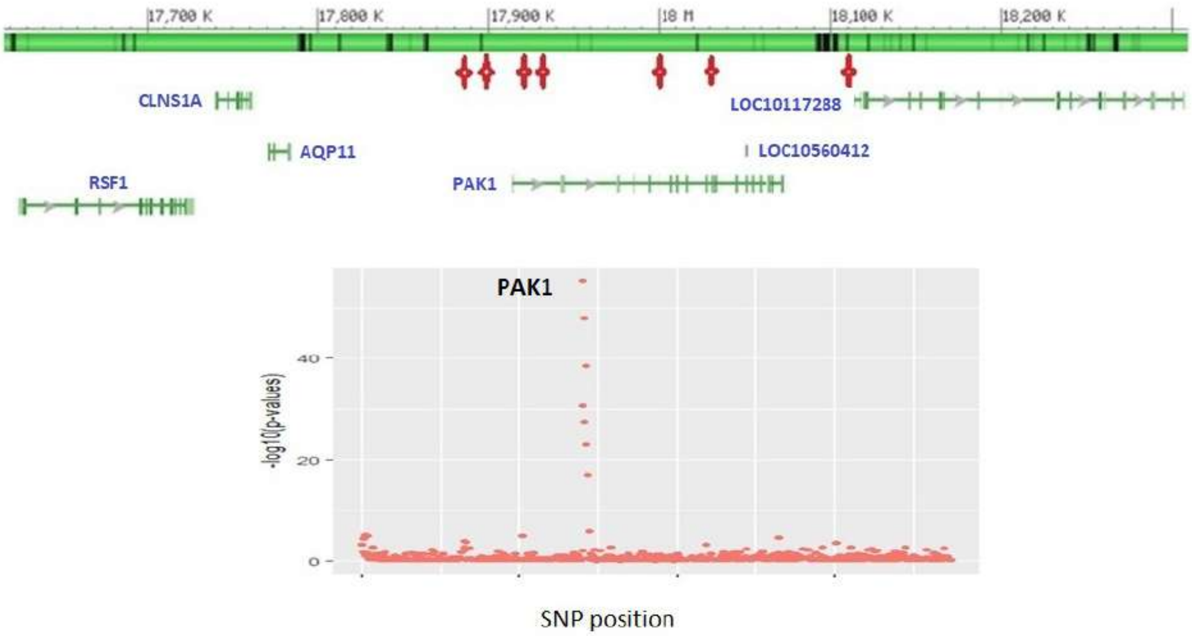
17,700 K
17,800 K
17,900 K
18 M
18,100 K
18,200 K
CLNS1A
LOC10117288
AQP11
LOC10560412
RSF1
PAK1
PAK1
-log10(p-values)
40
20
0
SNP position

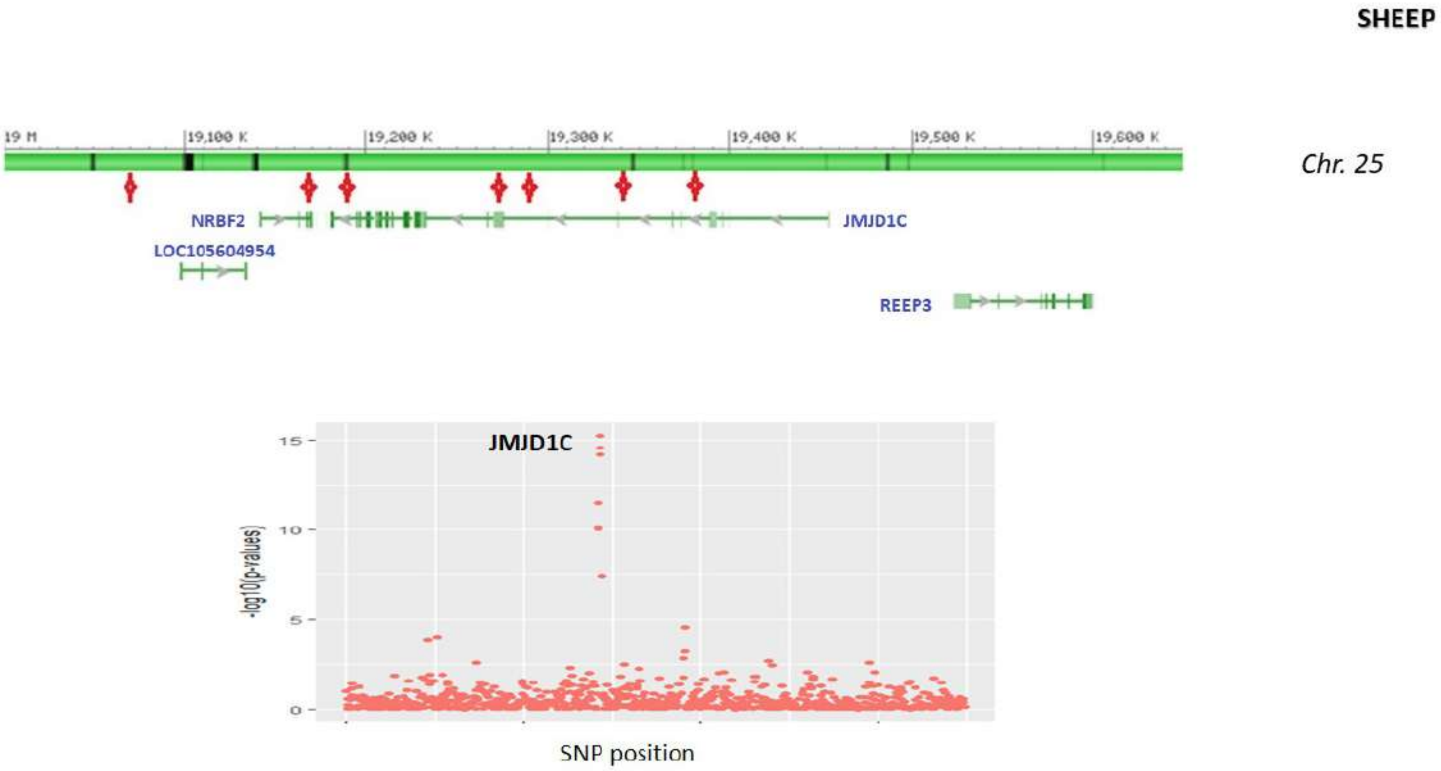
SHEEP
Chr. 25
19 M
19,100 K
19,200 K
19,300 K
19,400 K
19,500 K
19,600 K
NRBF2
JMJD1C
LOC105604954
REEP3
JMJD1C
15
10
5
0
-log10(p-values)
SNP position

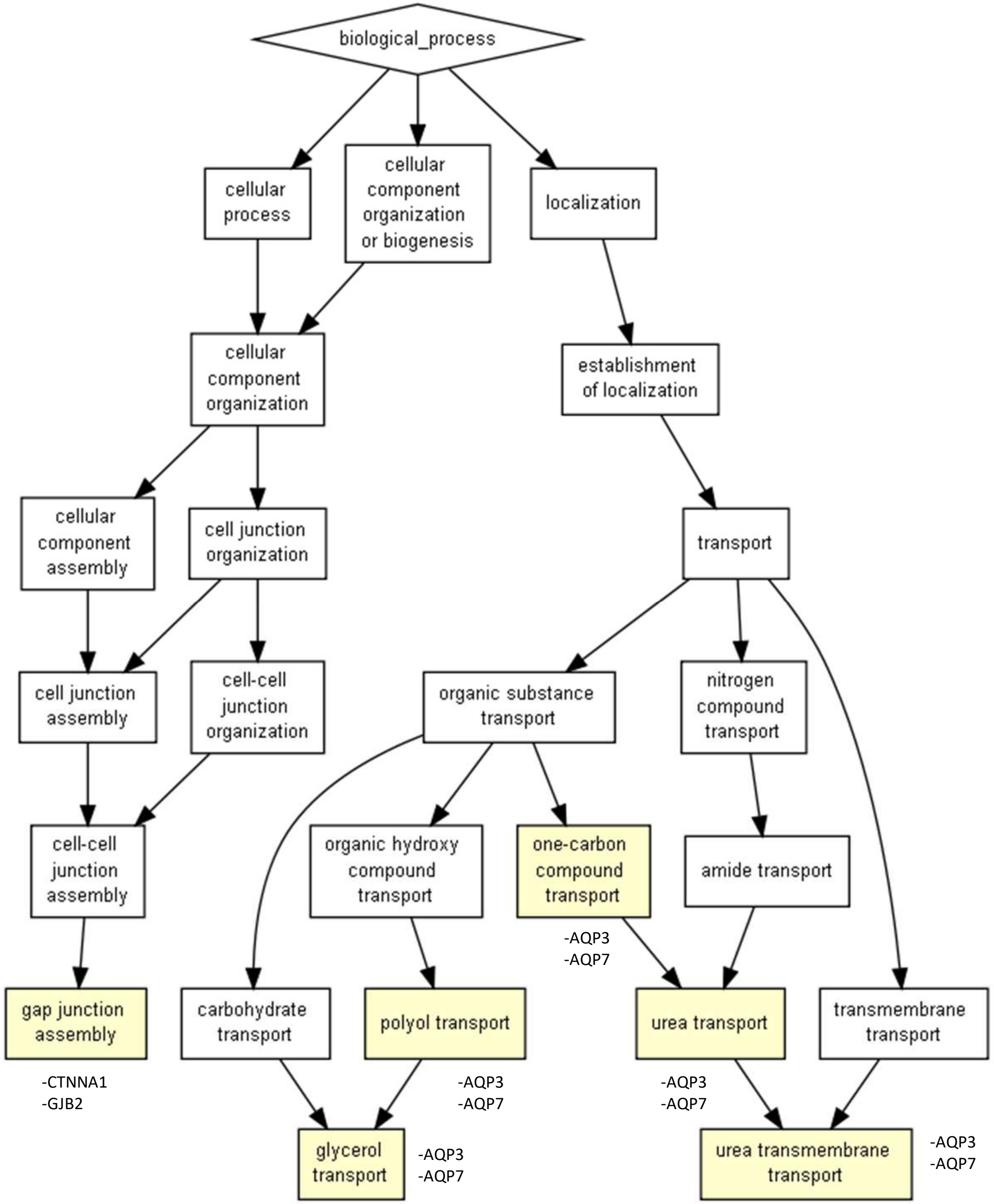
biological_process
cellular process
cellular component organization or biogenesis
localization
cellular component organization
establishment of localization
cellular component assembly
cell junction organization
transport
cell junction assembly
cell-cell junction organization
organic substance transport
nitrogen compound transport
cell-cell junction assembly
organic hydroxy compound transport
one-carbon compound transport
amide transport
-AQP3
-AQP7
gap junction assembly
carbohydrate transport
polyol transport
urea transport
transmembrane transport
-CTNNA1
-GJB2
-AQP3
-AQP7
-AQP3
-AQP7
glycerol transport
-AQP3
-AQP7
urea transmembrane transport
-AQP3
-AQP7

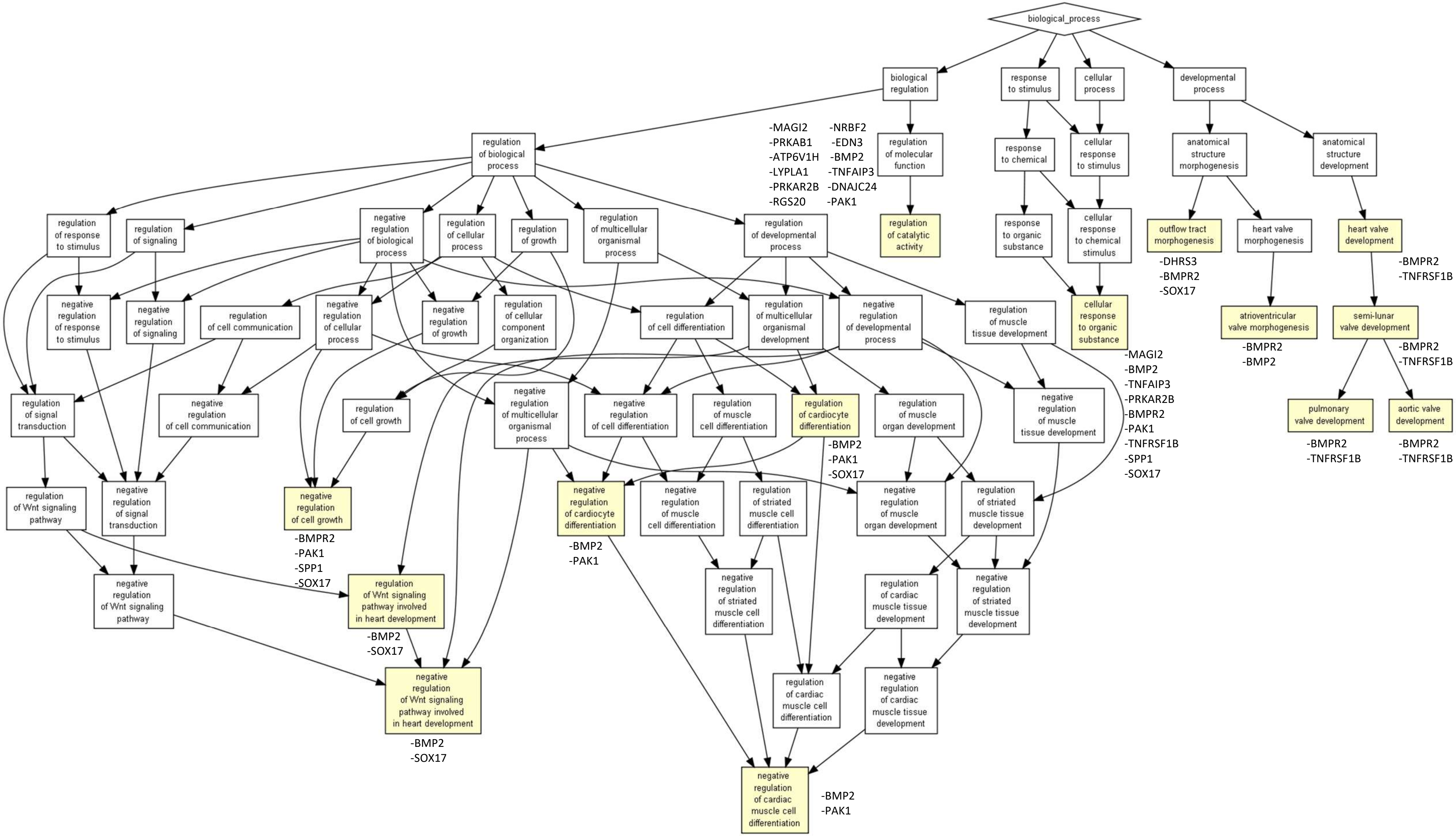
biological_process
biological regulation
response to stimulus
cellular process
developmental process
regulation of biological process
regulation of molecular function
response to chemical
cellular response to stimulus
anatomical structure morphogenesis
anatomical structure development
-MAGI2
-PRKAB1
-ATP6V1H
-LYPLA1
-PRKAR2B
-RGS20
-NRBF2
-EDN3
-BMP2
-TNFAIP3
-DNAJC24
-PAK1
regulation of response to stimulus
regulation of signaling
negative regulation of biological process
regulation of cellular process
regulation of growth
regulation of multicellular organismal process
regulation of developmental process
regulation of catalytic activity
response to organic substance
cellular response to chemical stimulus
outflow tract morphogenesis
heart valve morphogenesis
heart valve development
-DHRS3
-BMPR2
-SOX17
-BMPR2
-TNFRSF1B
negative regulation of response to stimulus
negative regulation of signaling
regulation of cell communication
negative regulation of cellular process
negative regulation of growth
regulation of cellular component organization
regulation of cell differentiation
regulation of multicellular organismal development
negative regulation of developmental process
regulation of muscle tissue development
cellular response to organic substance
atrioventricular valve morphogenesis
semi-lunar valve development
-BMPR2
-BMP2
-BMPR2
-TNFRSF1B
-MAGI2
-BMP2
-TNFAIP3
-PRKAR2B
-BMPR2
-PAK1
-TNFRSF1B
-SPP1
-SOX17
regulation of signal transduction
negative regulation of cell communication
regulation of cell growth
negative regulation of multicellular organismal process
negative regulation of cell differentiation
regulation of muscle cell differentiation
regulation of cardiocyte differentiation
regulation of muscle organ development
negative regulation of muscle tissue development
pulmonary valve development
aortic valve development
-BMP2
-PAK1
-SOX17
-BMPR2
-TNFRSF1B
-BMPR2
-TNFRSF1B
regulation of Wnt signaling pathway
negative regulation of signal transduction
negative regulation of cell growth
negative regulation of cardiocyte differentiation
negative regulation of muscle cell differentiation
regulation of striated muscle cell differentiation
negative regulation of muscle organ development
regulation of striated muscle tissue development
-BMPR2
-PAK1
-SPP1
-SOX17
-BMP2
-PAK1
negative regulation of Wnt signaling pathway
regulation of Wnt signaling pathway involved in heart development
negative regulation of striated muscle cell differentiation
regulation of cardiac muscle tissue development
negative regulation of striated muscle tissue development
-BMP2
-SOX17
negative regulation of Wnt signaling pathway involved in heart development
regulation of cardiac muscle cell differentiation
negative regulation of cardiac muscle tissue development
-BMP2
-SOX17
negative regulation of cardiac muscle cell differentiation
-BMP2
-PAK1

### Literature and classification of the genes identified

The highlighted selection signatures can be clustered according to different major adaptation mechanisms. Classifying genes according to the major functions in which they can be involved should be regarded with due caution. Indeed, a large number of genes are pleiotropic, like ADAMTS6 (see Wu et al., 2019). For the classification we tried to identify in the literature the major mechanisms associated with the genes under consideration. For example, the proposed classifications did not take into account the fact that MAGI2, KDM4C and JMJD1C are also related to spermatogenesis (Sujit et al., 2018); nor the fact that VPS13 family may play a role in the maintenance of neuronal processes in mammals; or that PCDH9 and KLHL1, were found associated with fat tail deposition by the study of Mastrangelo et al. (2019). Finally, a correlation between genetic variation and a trait does not imply a causal relationship.

- Selection signatures involved in lipid storage

A large number of the genes identified seems to be involved in lipid metabolism. Most of them are also involved in the production/composition of milk. (i) For goats we can mention: PTGER3 (Strong et al., 1992), CTBP1 (Xu et al., 2017), TCF12 (Ozdemir Ozgenturk et al., 2017; Dunner et al., 2013), SCFD2 (Yodklaew et al., 2017; Glastonbury et al., 2016), UHRF1BP1 (Justice et al., 2018), BTRC (Ishimoto et al., 2017), CTNNA1/SIL1(fat tail deposition in sheep: Ahbara et al., 2019), VPS13B (Liu et al., 2018; Capitan et al., 2014), WDPCP (Sazzini et al., 2016) and SUCLG2 (Silva-Vignato et al. 2019; Di Gerlando et al., 2019).

WDPCP has also been identified as being under selection in Ethiopian sheep of high altitude (Edea et al., 2019) as well as in Tibetan chicken (Wang et al., 2015), suggesting a probable mechanism common to these taxa of homeoterms. WDPCP is related to the BBSome, a protein complex in primary cilia with important role in leptin receptor (LepR) trafficking; in humans it is linked to obesity (Foucan et al., 2018).

(ii) The identified genes in sheep implied in lipid metabolism and milk production are the followings: ARHGAP26/NR3C1 (Edea et al., 2017), ABCD4/VSX2/LIN52 (fat tail deposition: Zhu et al., 2019), POU2F1 (Lewinsky et al., 2005; Zhao et al., 2002; Pérez-Montarelo et al., 2014), VPS13D, MSRB3 and BMP2 (these last two genes are discussed in the article).

We identified VPS13B in goats and VPS13D in sheep. The gene VPS13B was found under selection in Moroccan goats (Benjelloun, 2015), was associated to adaptation to dry areas in Chinese sheep (Yang et al., 2016), and with high altitude conditions in Yaks (Qi et al., 2019). Mammalian VPS13 proteins are involved in lipid transfer (Gao et al., 2018). VPS13D is involved in lactase persistence and lipid pathway in the Maasai (Wagh et al., 2012). The VPS13B protein could play an important role in the development of nerve cells (neurons) and in the growth and development of adipocytes as reported by Seifert et al. (2011).

- Selection signatures involved in seasonal patterns and circadian behaviors

The analysis detected a strong link between the environmental gradient and the BMPR1B gene in goats. This gene is involved in ovarian function and fertility (Shokrollahi & Morammazi, 2018; Mulsant et al., 2001; Souza et al., 2001; Wilson et al., 2001). The influence of the photoperiod on ovarian function is well known in mammals, where melatonin provides the hormonal signal transducing day length (Walton et al., 2011). This could explain the correlation between BMPR1B and the latitudinal gradient, i.e. a proxy for seasonality, found in our study. Moreover, seasonal patterns are frequently expressed at many levels (foraging, adipocity, growth, immune function *etc.*, see the review by Prendergast, Nelson, & Zucker, 2009). These elements could explain the link established by Cao et al. (2015) between BMPR1B and growth in sheep. The role of this gene in thermogenesis, has been demonstrated in murine adipose tissue (Whittle et al., 2015).

Similarly, SOX2, implicated in behavioral rhythms linked to environmental light cycles (Cheng et al., 2019) and DPH6, a circadian rhythm-related GO categories, were also identified by the LFMM approach as correlated with the environmental gradient in goats (see detailed discussion in the article).

- Selection signatures involved in hypoxia and/or heat stress response

Hypoxia is a major stimulus for physiological processes such as adaptation to life at high altitude, but it is also an important factor of many human pathologies, such as cancer (Kenneth & Rocha, 2008). We identified MAT2A in goats and JMJD1C and KDM4C (a JmjC domain-containing protein) in sheep. It is shown that hypoxia induces genomic DNA demethylation through the activation of HIF-1α and transcriptional upregulation of MAT2A (Liu et al., 2011). Moreover, the Jumonji C (JmjC) domain contains demethylases, allowing genes such as JMJD1C and KDM4C to trigger mechanisms to respond to hypoxia (Xia et al., 2009; Melvin & Rocha, 2012). In sheep, EDN3, RAB22A and DNAJC24 were identified by us. The gene EDN3, was previously associated with hypoxia tolerance in cetaceans as vasoconstrictor-related gene (Tian et al., 2016). RAB22A was found involved in hypoxia process in humans (Wang et al., 2014). DNAJC24 was associated with high altitude adaptation in sheep (Yang et al., 2016) and appeared under selection in Ugandan goats (Onzima et al., 2018). DNAJC24 belongs to the heat shock protein 70 family, that are produced in response to exposure to stressful conditions, as cold, UV light, *etc*.

The signature centered on PIGL/LOC102188626 in goats, and bordered by NCOR1/ADORA2B deserves special attention. PIGL was previously identified in Chineese sheep at high altitudes (Yang et al., 2016). NCOR1 belong to clock circadian gene network in cattle (Wang et al., 2015), whereas ADORA2B (an adenosine receptor, belonging to the hypoxia database) appears implicated in thermal adaptation (Wollenberg Valero et al., 2014).

Finally, in goats, one selection signature targeted UBE2R2/UBAP2, associated with hypoxia tolerance in humans (Udpa et al., 2014), and was bordered by AQP7/AQP3, which belong to the hypoxia database (see detailed discussion in the article).

- Selection signatures implicated in coat color or horns

First, considering morphological adaptations, two genes known to affect coat colour, ADAMTS20 (Oget, Servin, & Palhière 2019; Bertolini et al., 2018) and TYRP1 (Gratten et al., 2007; Becker et al., 2015), were identified in goats. The pigmentation (fiber and skin), by influencing the ability to absorb solar radiation, corresponds to a primordial adaptive mechanism that is highly submitted to natural selection in mammals (Caro, 2005). In Western Europe, sheep have been subjected to strong artificial selection, over the past 200 years, on the quality of the wool but also on its colour; the white colour being particularly prized (Lauvergne, 1969; Mellah, 2015). This could explain why we did not found any selection signatures in sheep for this type of gene in connection with the environment.

With regard to horns, the RXFP2 gene is known to be strongly involved in this process (Johnston et al., 2011; Allais-Bonnet et al., 2013) and appeared to be selected in sheep. Indeed, the production of polled animals has been a major objective of artificial breeding for decades. Also, we were not surprised to observe a majority of sheep breeds of the dataset without horn (see detailed discussion in the article). In goats, the gene involved in the presence or absence of horn is pleiotropic, so that selecting individuals without horn induces undesirable secondary phenotypes, often in association with reduced fertility (Liron, 2011).

- Selection signatures implicated in immunity

In sheep, MASP-1, a serine protease, was identified. It functions as a component of the lectin pathway of complement activation, and as such, plays an essential role in the innate and adaptive immune response (Ammitzbøll et al., 2013). In goat, the genes PRNP, NFATC3/LRRC36 appeared selected. A large literature links PRNP to susceptibility to prion disease, in small ruminants (see Greenlee, 2019 for a review). The NFAT (nuclear factor of activated T cells) family of transcription factors is well documented for its role in T lymphocytes (Minematsu et al., 2011). LRR-containing proteins are also well known for their role in innate immunity (Ng et al., 2011). Finally, JAZF1, identified in sheep, plays a role in adipose tissue inflammation by limiting macrophage populations and restricting their antigen presentation function. Thus, this gene seems to act at the interface between immune system and adipose metabolism (Meng et al., 2018).

- Signatures involved in lung function

The GSTCD gene, highlighted in goat, plays a role in cellular homeostasis associated with lung function (Artigas et al. 2015; Obeidat et al., 2013; Henry et al., 2019). The SLC34A2 gene, identified in the goat dataset as selected and very close to SEL1L3, was previously associated with high altitude conditions in cattle (Verma et al., 2018). Moreover, Edea et al. (2019) linked SEL1L3, with high altitude adaptation in sheep. In humans, SLC34A2 is mainly located in the lungs, and produces and recycle surfactant, that makes breathing easy (Ma et al., 2018). Finally, SEL1L3 belong to the hypoxia regulated proteins database (http://www.hypoxiadb.com).

- Signature selection involved in neuronal function

MAGI2, detected in sheep, may play a central role in adaptation as it was also identified by Edea et al. (2019) in high altitude Ethiopian sheep, by Qi et al. (2019) in high altitude yaks and by Yang et al. (2016) associated with desert and arid conditions in Chinese sheep. The MAGI2 gene that belongs to the hypoxia regulated proteins database, plays the critical role of maintaining synaptic strength (Danielson et al., 2012).

BTBD9, highlighted in goat, is mainly expressed in brain structure (Stefansson et al., 2007). It plays a role in the regulation of ion channels' tetramerization and gating, and hence is potentially involved in neuronal signal transduction (Stogios et al., 2005).

AUTS2, also detected in goat, is a critical neuronal gene implicated in many neurological disorders as autism. Interestingly, several predicted protein–protein interaction domains (SH2 and SH3) were observed for this protein (Oksenberg, Stevison, Wall, & Ahituv, 2013), whereas SH3 gene was found under selection in the sheep dataset. SH3 domains are eukaryotic protein domains that are involved in a plethora of cellular processes including signal transduction.

The MDGA2 gene, was identified in goats. Mutations in MDGA2 were associated with autism potentially through its ability to modulate excitatory/inhibitory transmission. Indeed, MDGA2 is able to block neuroligin-1 interaction with neurexins and then to suppresses excitatory synapse development (Connor et al., 2016).

The genes TRPC4, TRPC6 and NBEA are discussed in detailed in the manuscript.

- "Complex" selection signatures

The following selection signatures appear more difficult to analyze and are processed on a case-by-case basis, due to the difficulty of categorizing them. In goats, HERC6 was identified as being under selection at the level of a rather small signature that was framed by HERC5 on one side and ABCG2 on the other. In sheep, HERC6 was found in a particularly large signature (it should be noted that PCAdapt predicts a zone lagged with respect to LFMM). A total of 16 genes were involved in this signature, which also included HERC5 and ABCG2. The HERC family emerged more than 595 million years ago. HERC5 and HERC6 contain a domain implicated in antiviral activity, which gives them great importance in terms of intrinsic immunity (Paparisto et al., 2018). In terms of adaptation, HERC6 was found to be linked to arid environments adaptation (Yang et al., 2016; Mdladla, 2016) and also to stress grazing adaptation (Mwacharo et al., 2018). With regard to the other genes found in the sheep signature: a very large literature links ABCG2/SPP1/LAP3/NCAPG/MEPE to milk production, particularly ABCG2 (see Ruiz-Larrañaga et al., 2018); the remaining genes LCORL/LAP3/FAM184B may be related to growth traits (Al-Mamun et al., 2015; Lindholm-Perry et al., 2001; Bongiorni et al., 2012; Eberlein et al., 2009). Only more in-depth research will make it possible to understand the mechanisms underlying the adaptations in relation to these genome areas.

In goat, the large selection signature located on chromosome 12 and including amongst others genes RNF17, ZMYM2, PARP4 and XPO4, seems to play an important role in adaptation processes. Here again the situation is particularly complex and fine studies focused on this area are needed, while this area appears linked with lipid metabolic processes, growth, DNA repair, and also associated with adaptation to high altitude, stress grazing and arid environments (see Supplementary Table 3).
